# Exploiting the Cullin E3 Ligase Adaptor Protein SKP1 for Targeted Protein Degradation

**DOI:** 10.1101/2023.10.20.563371

**Authors:** Seong Ho Hong, Akane Osa, Oscar W. Huang, Ingrid E. Wertz, Daniel K. Nomura

## Abstract

Targeted protein degradation with Proteolysis Targeting Chimeras (PROTACs) is a powerful therapeutic modality for eliminating disease-causing proteins through targeted ubiquitination and proteasome-mediated degradation. Most PROTACs have exploited substrate receptors of Cullin-RING E3 ubiquitin ligases such as cereblon and VHL. Whether core, shared, and essential components of the Cullin-RING E3 ubiquitin ligase complex can be used for PROTAC applications remains less explored. Here, we discovered a cysteine-reactive covalent recruiter EN884 against the SKP1 adapter protein of the SKP1-CUL1-F-box containing SCF complex. We further showed that this recruiter can be used in PROTAC applications to degrade neo-substrate proteins such as BRD4 and the androgen receptor in a SKP1- and proteasome-dependent manner. Our studies demonstrate that core and essential adapter proteins within the Cullin-RING E3 ubiquitin ligase complex can be exploited for targeted protein degradation applications and that covalent chemoproteomic strategies can enable recruiter discovery against these targets.

## Introduction

Targeted protein degradation with heterobifunctional Proteolysis Targeting Chimeras (PROTACs) consisting of a protein-targeting ligand linked to E3 ubiquitin ligase recruiters has arisen as a powerful therapeutic modality for degrading disease-causing proteins through proteasome-mediated degradation. While there are >600 E3 ligases, most PROTACs have utilized a small number of E3 ligases for recruitment including cereblon and VHL ^1–4^. Many additional recruiters have been discovered in recent years against additional E3 ligases, including cIAP, MDM2, DCAF16, DCAF11, RNF114, RNF4, FEM1B, KEAP1, and DCAF1 ^5–16^. While some of these E3 ligases are essential for cancer cell viability, many of them are not, and as such, resistance mechanisms can potentially arise in cancer cells that impair E3 ligase recruitment and render the PROTAC ineffective. Developing recruiters against core components of the ubiquitin-proteasome machinery, such as against Cullin-RING E3 ligase complex adaptor proteins such as DDB1, SKP1, or ELOB/ELOC in CUL1-CUL7 E3 ligases that are essential to cell viability may avoid potential future resistance mechanisms that may arise from PROTACs ^17–19^.

Recruitment of Cullin-RING E3 ligase adaptor proteins for targeted protein degradation has precedence. Slabicki and Ebert *et al*. discovered that a cyclin-dependent kinase inhibitor CR8 degrades cyclin K through acting as a molecular glue between CDK12-cyclin K and the CUL4 adaptor protein DDB1 ^20^. Various viruses have also co-opted adaptor proteins within Cullin-RING E3 ligase complexes. For example, the human papilloma virus (HPV) E7 protein is responsible for host cellular oncogenic transformation and co-opts Elongin C in the CUL2 complex to ubiquitinate and degrade the RB tumor suppressor ^21^. Paramyxovirus and Simian Virus 5 V proteins both co-opt the CUL4 adaptor protein DDB1 ^21^. We have also recently discovered a covalent DDB1 recruiter that can be exploited for PROTAC applications ^22^.

In this study, we investigated whether the CUL1-RING E3 ligase adaptor protein SKP1 could be recruited for PROTAC-mediated degradation of neo-substrates. We used covalent chemoproteomic approaches to discover a SKP1 recruiter that can be used for PROTACs to degrade neo-substrates in cells.

## Results

### Discovering a Covalent Recruiter Against SKP1

We sought to identify a covalent binder for SKP1. We screened 1284 cysteine-reactive covalent ligands in a gel-based activity-based protein profiling (ABPP) screen competing covalent ligands against a fluorophore-conjugated cysteine-reactive iodoacetamide probe (IA-rhodamine) labeling of pure human SKP1-FBXO7-CUL1-RBX1 complex **(Figure 1a; Figure S1; Table S1)** ^14,23^. The top hit that arose from this screen was EN884 that showed dose-response displacement of cysteine-reactive probe labeling of pure human SKP1 protein in the SKP1-FBXO7-CUL1-RBX1 complex **(Figure 1b-1c)**. We next performed tandem mass spectrometry (MS/MS) analysis on the SKP1-FBXO7-CUL1-RBX1 complex incubated with EN884 to identify the site of modification. We identified one site that was modified—C160 on SKP1 **(Figure 1d)**. C160 resides at the C-terminus of SKP1, a region that is predicted to be an intrinsically disordered region (IDR) of the protein **(Figure S1)**. Previous studies have indicated its conformational flexibility plays a pivotal role in SKP1 interaction with various F-Box domains characterized by diverse sequences and structures ^24,25,25–28^. Consequently, the C-terminus of SKP1 adopts a unique conformation contingent upon its interaction with specific F-Box domains **(Figure 1e; Figure S3)**. Thus, we hypothesize that our covalent SKP1 recruiters may exploit and bind in between the collective interfaces between SKP1 and CUL1 substrate receptors. While structural information on FBXO7 remain elusive, analyses of SKP1 – Fbox domain protein complexes, such as SKP2, FBG3, FBXW7, or FBXO2, suggests that perhaps EN884 engages only a subset of SKP1 protein engaged with specific CUL1 substrate receptors **(Figure 1e; Figure S3)**.

**Figure 1.**
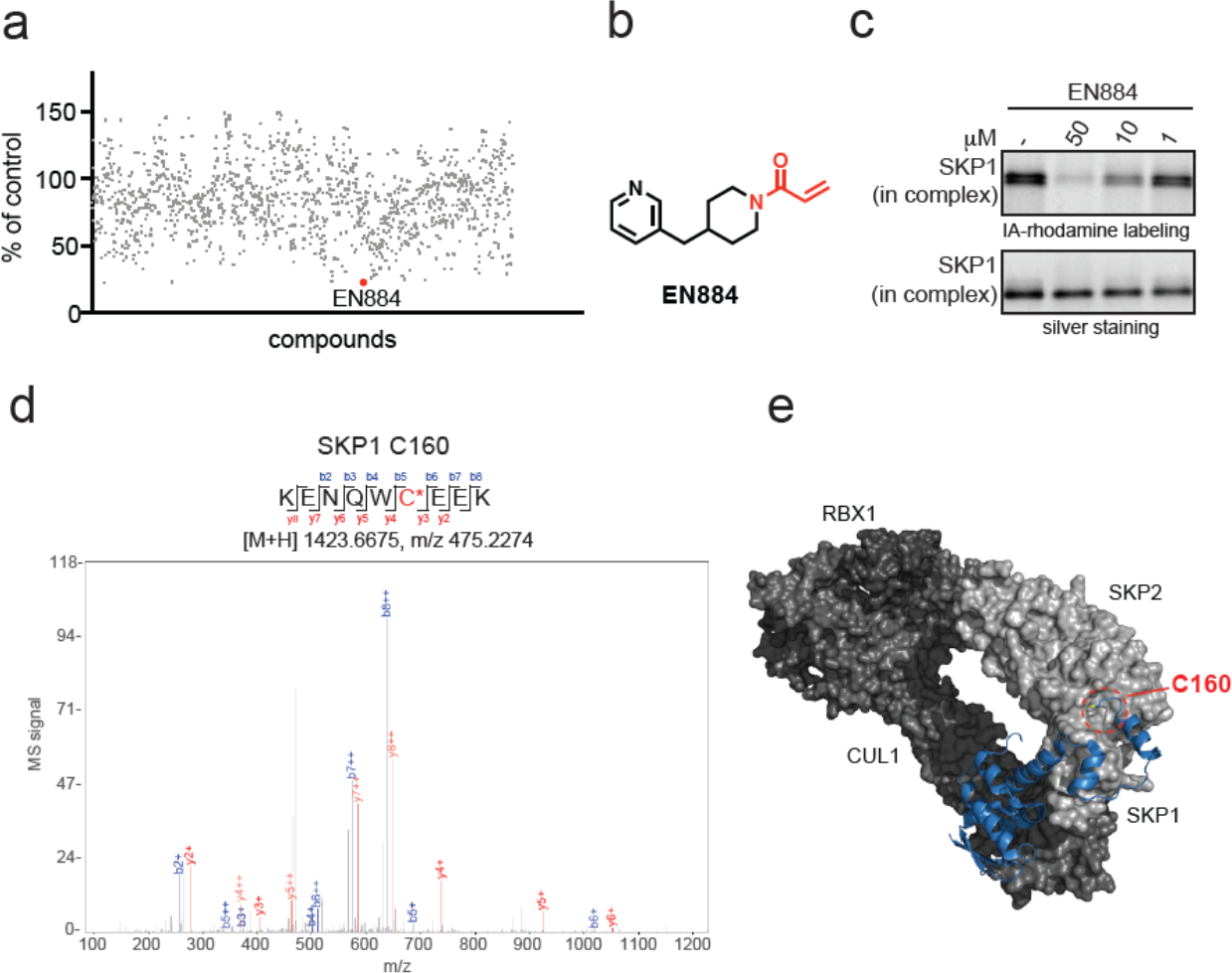
Discovery of a covalent SKP1 recruiter. **(a)** Gel-based ABPP screening of a cysteine-reactive covalent ligand library (50 μM) against a rhodamine-functionalized iodoacetamide (IA-rhodamine) labeling of pure SKP1 protein in the SKP1-FBXO7-CUL1-RBX1 complex. Individual compound values are noted as percent of labeling compared to that of DMSO vehicle control. EN884 was the top hit that showed the most inhibition of probe labeling. **(b)** Structure of EN884 with the cysteine reactive acrylamide warhead in red. **(c)** gel-based ABPP of EN884 competition against IA-rhodamine labeling. Pure SKP1 in the SKP1-FBXO7-CUL1-RBX1 complex was pre-incubated with EN884 1 h prior to labeling with IA-rhodamine (100 nM) for 30 min after which proteins were separated by SDS/PAGE and visualized by in-gel fluorescence. Protein loading was assessed by silver staining. **(d)** Mass spectrometry analysis of EN884 modification on SKP1. The SKP1-FBXO7-CUL1-RBX1 complex was incubated with EN884 (50 μM) for 1 h and tryptic digests from the complex were subsequently analyzed for the EN884 covalent adduct on a cysteine. Shown is the MS/MS data for EN884 modification on SKP1 C160. **(e)** Shown is the location in the circle of SKP1 C160 in the CUL1 complex with SKP1 in blue, in complex with SKP2, RBX1, and CUL1. Shown is an SCF complex model derived from superimposing the crystal structures of Skp1-Skp2-Cks1 with p27 peptide (PDB ID: 2AST) and Cul1-Rbx1-Skp1-Skp2 (PDB ID: 1LDK). Data shown in (**b)** are from n=3 biologically independent replicates per group.

We next synthesized two alkyne-functionalized probes based on the EN884 hit with either exit vectors off either the *meta* or *para*-position of the phenyl ring to yield SJH1-37-m or SJH1-37-p, respectively **(Figure 2a)**. Based on gel-based ABPP assessment of binding of each of these probes to the SKP1 complex, the *meta*-position was significantly more favored compared to *para* **(Figure 2b)**. Interestingly, and consistent with the position of C160 on SKP1 interfacing with CUL1 substrate receptors, EN884 and SJH1-37m did not bind to monomeric SKP1 **(Figure 2c)**. These results indicate that the rest of the CUL1 complex, or at least the substrate receptor, was required for ligand binding to SKP1. We further showed that SJH1-37m engages SKP1 in cells through pulldown of SKP1, but not unrelated proteins such as GAPDH, from probe treatment and subsequent appendage of a biotin enrichment handle by copper-catalyzed azide-alkyne cycloaddition (CuAAC) and avidin-enrichment **(Figure 2d)**. We next performed mass spectrometry-based ABPP to identify the cysteine engaged in SKP1 by EN884 in HEK293T cells using previously established isotopic desthiobiotin-ABPP (isoDTB-ABPP) ^29^. We found that EN884 significantly engaged C160 on SKP1 by 7% **(Figure 2e; Table S2)**. This minimal engagement observed with our covalent SKP1 recruiter is consistent with minimal engagement observed with other covalent E3 ligase recruiters that have been deployed in PROTAC applications, including for DCAF16, DCAF11, and RNF114^7,8,11^. This low degree of engagement is also consistent with EN884 potentially only engaging with a subset of SKP1 in the cell that is engaged with certain CUL1 substrate receptors. We also performed a pulldown proteomics experiment with the SJH1-37m probe and demonstrated 2.8-fold and statistically significant enrichment of SKP1 over vehicle-treated controls **(Figure S4; Table S3)**. The covalent hit is not yet selective with 414 other targets significantly enriched among 5928 proteins identified, but there were no other proteins involved in the CUL1 E3 ligase complex pulled down by SJH1-37m **(Table S3)**.

**Figure 2.**
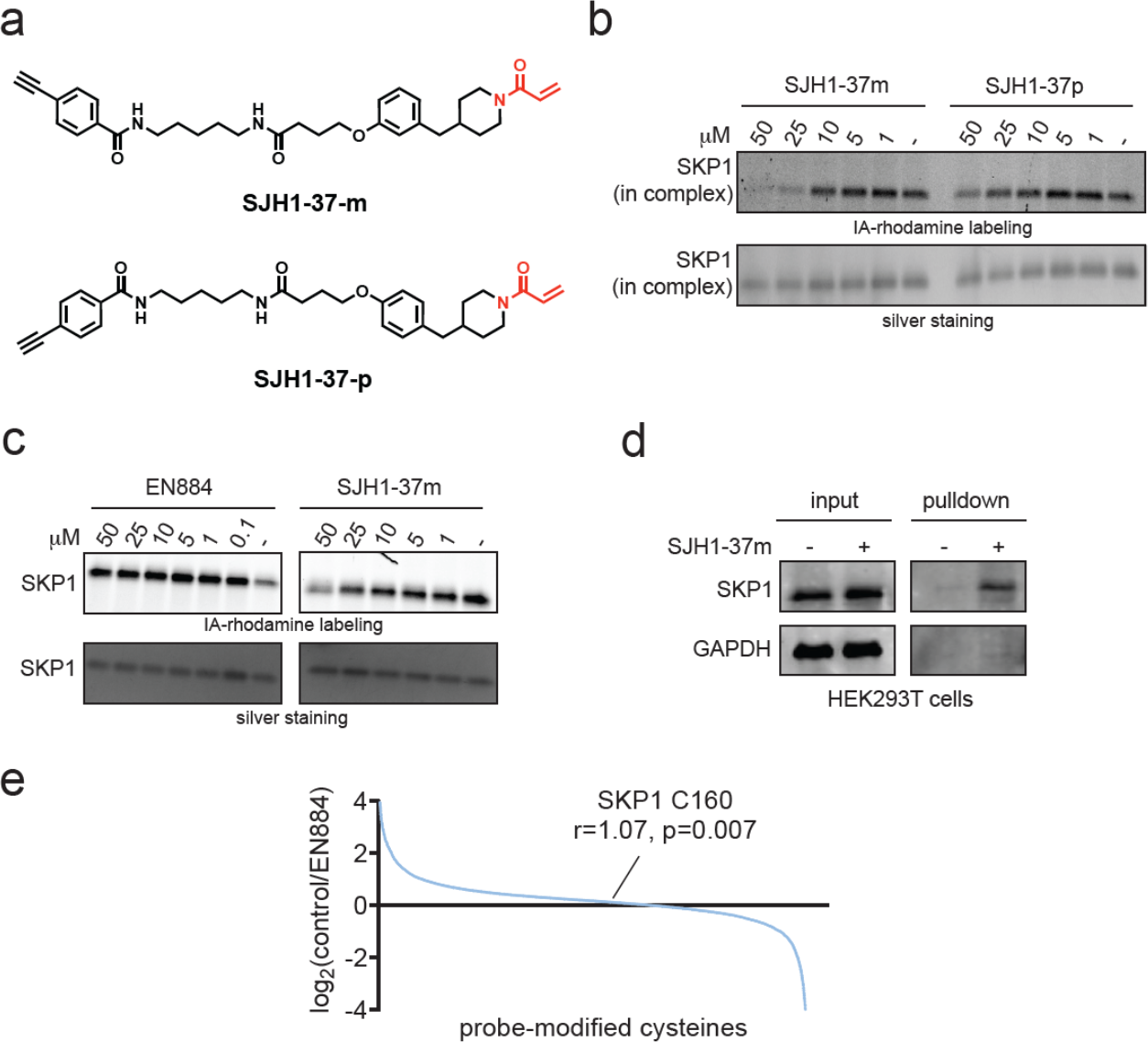
Characterization of covalent SKP1 recruiter. **(a)** Structures of two alkyne-functionalized probes based on EN884. **(b)** Gel-based ABPP of both probes against IA-rhodamine labeling of pure SKP1 in the SKP1-FBXO7-CUL1-RBX1 complex as described in **(a). (c)** Gel-based ABPP of EN884 and SJH1-37m against pure SKP1 not in complex assessed as described in **(b). (d)** SKP1 pulldown using SJH1-37m probe. HEK293T cells were treated with DMSO vehicle or SJH1-37m (50 μM) for 4 h. Lysates were subjected to copper-catalyzed azide-alkyne cycloaddition (CuAAC) to append on a biotin enrichment handle after which probe-modified proteins were avidin-enriched and separated on SDS/PAGE and SKP1 and an unrelated protein GAPDH were detected by Western blotting. **(e)** isoDTB-ABPP analysis of EN884 in HEK293T cells. HEK293T cells were treated with DMSO vehicle or EN884 (50 μM) for 4 h, after which lysates were labeled with an alkyne-functionalized iodoacetamide probe (IA-alkyne) and control and treated cells were subjected to CuAAC with a desthiobiotin-azide with an isotopically light or heavy handle, respectively. After the isoDTB-ABPP procedure, probe-modified peptides were analyzed by LC-MS/MS and control vs treated probe-modified peptide ratios were quantified. Data shown in (**b, c, d, e)** are from n=3 biologically independent replicates per group.

### Using Covalent SKP1 Recruiter in PROTACs to Degrade BRD4

To demonstrate that our covalent SKP1 ligand could be used in PROTAC applications, we synthesized four different PROTACs linking a derivative of EN884 to the BET family inhibitor JQ1 through either a C2, C4, C5, or PEG3 linker to generate SJH1-51A, SJH1-51B, SJH1-51C, and SJH1-51D, respectively **(Figure 3a)**. All four PROTACs degraded the short, but not long isoform, of BRD4 in HEK293T cells with the most robust degradation observed with SJH1-51B with a C4 alkyl linker **(Figure 3b-3d)**. While we observed isoform-specific BRD4 degradation in HEK293T cells, both isoforms were degraded by SJH1-51B in MDA-MB-231 breast cancer cells **(Figure 4a)**. Tandem mass tagging (TMT)-based quantitative proteomic profiling of SJH1-51B by in MDA-MB-231 cells showed moderately selective BRD4 degradation with 16 proteins significantly reduced in levels by >50 % **(Figure 4b**; **Table S4)**.

**Figure 3.**
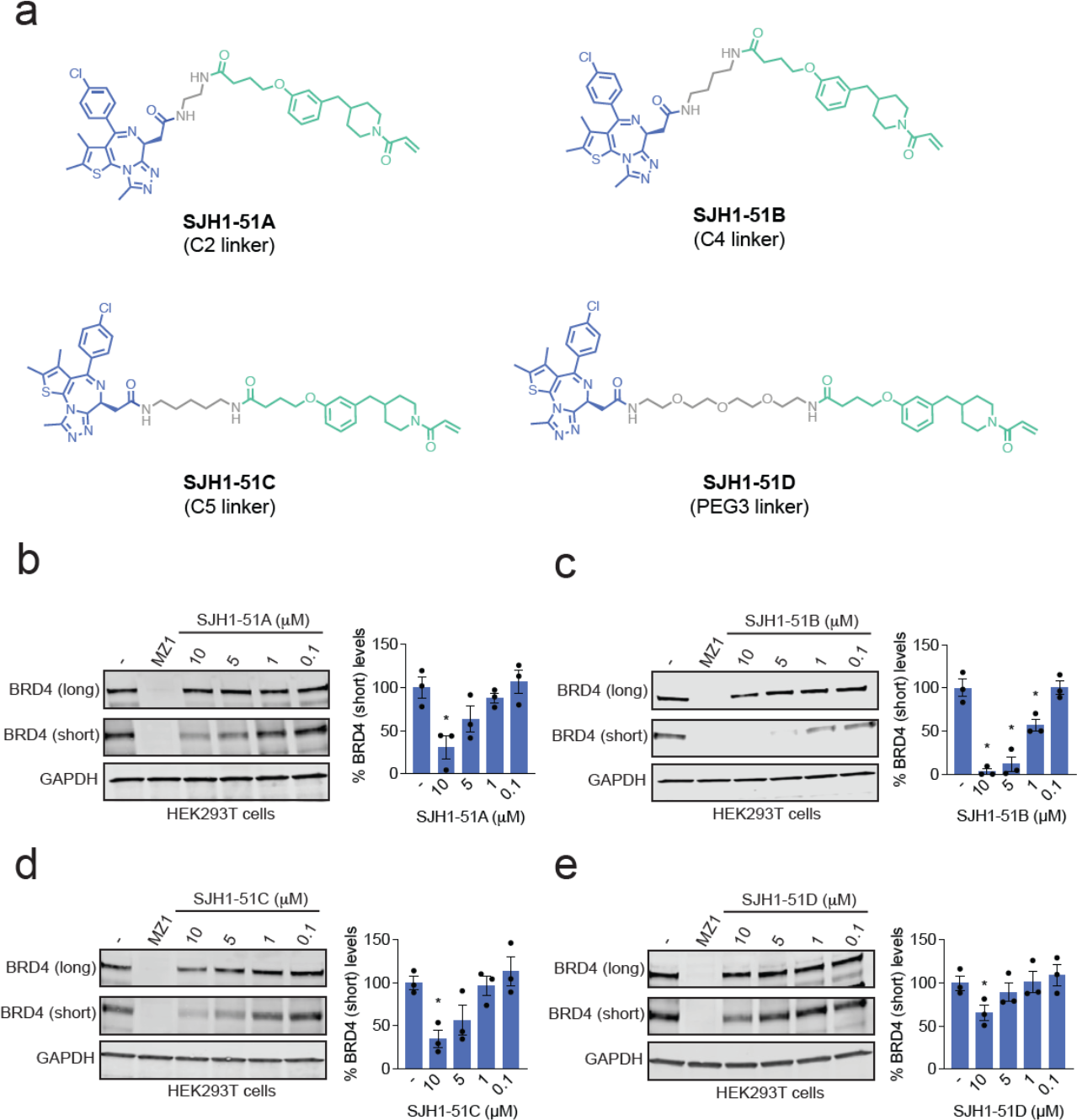
SKP1-based BRD4 degraders. **(a)** Structures of 4 BRD4 degraders using covalent SKP1 recruiters in green linked via a linker in grey to the BET family inhibitor JQ1 in blue via a C2, C4, or C5 alkyl linker or PEG3 linker. **(b-e)** BRD4 degradation in cells. HEK293T cells were treated with DMSO vehicle, MZ1 (1 μM), or each degrader for 24 h. The long and short isoforms of BRD4 and loading control GAPDH were assessed by Western blotting and bands were quantified by densitometry and normalized to GAPDH. Blots are representative of n=3 biologically independent replicates/group. Bar graphs show average ± sem and individual replicate values of the BRD4 short isoform. Significance is expressed as *p<0.05 compared to vehicle-treated controls.

**Figure 4.**
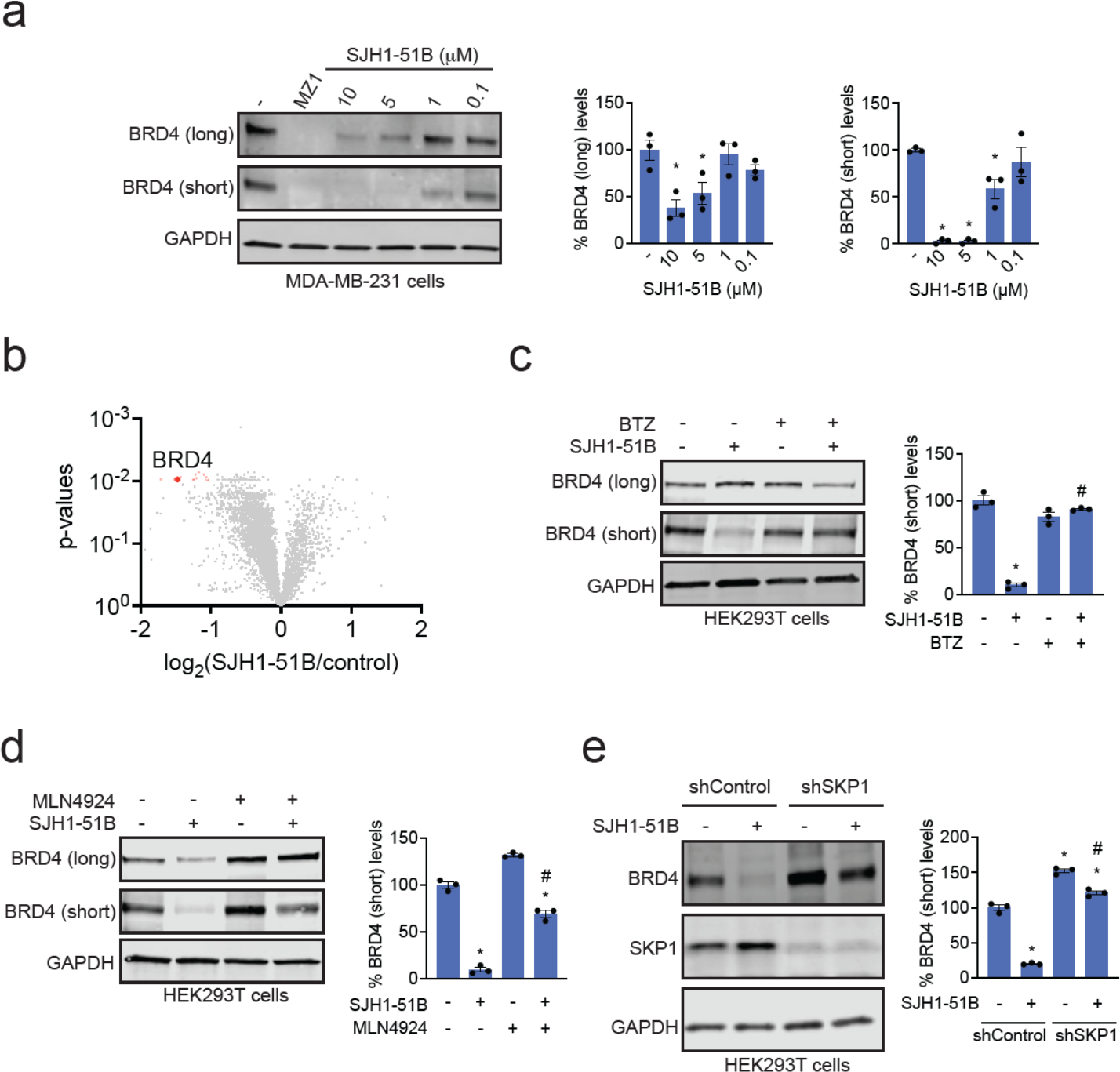
Characterization of BRD4 degrader. **(a)** BRD4 degradation in MDA-MB-231 cells. MDA-MB-231 cells were treated with DMSO vehicle, MZ1 (1 μM), or SJH1-51B for 24 h. The long and short isoforms of BRD4 and loading control GAPDH were assessed by Western blotting and bands were quantified by densitometry and normalized to GAPDH. **(b)** TMT-based quantitative proteomic profiling of SJH1-51B in MDA-MB-231 cells. MDA-MB-231 cells were treated with DMSO vehicle or SJH1-51B (10 μM) for 24 h. **(c, d)** Proteasome and NEDDylation-dependence of BRD4 degradation. HEK293T cells were pre-treated with DMSO vehicle or bortezomib (1 μM) in **(c)** or MLN4924 (1 μM) in **(d)** for 1 h prior to treatment of cells with DMSO vehicle or SJH1-51B (10 μM) for 24 h. BRD4 and loading control GAPDH levels were assessed by Western blotting and quantified. **(e)** Attenuation of BRD4 degradation by SKP1 knockdown. shControl and shSKP1 HEK293T cells were treated with DMSO vehicle or SJH1-51B (5 μM) for 24 h. Short BRD4 isoform, SKP1, and loading control GAPDH levels were assessed by Western blotting and quantified. Blots in **(a, c, d, e)** are representative of n=3 biologically independent replicates/group. Bar graphs show average ± sem and individual replicate values of BRD4 levels. Significance is expressed as *p<0.05 compared to vehicle-treated controls, and #p<0.05 compared to either SJH1-51B treatment alone in **(c, d)** or compared to SJH1-51B-treated shControl cells in **(e)**.

Consistent with proteasome-mediated degradation, SJH1-51B-mediated BRD4 degradation was attenuated by pre-treatment with the proteasome inhibitor bortezomib **(Figure 4c)**. We also showed dependence on NEDDylation required for active Cullin-RING E3 ligases with attenuation of BRD4 degradation by the NEDDylation inhibitor MLN4924 **(Figure 4d)**. Most importantly, we demonstrated that SKP1 knockdown completely attenuated BRD4 degradation by SJH1-51B in HEK293T cells **(Figure 4e)**. For BRD4, we showed that the non-reactive derivative of SJH1-51B, SJH1-51B-NC, did not degrade BRD4 in HEK293T cells **(Figure S5a-S5b)**. A time-course of BRD4 degradation with SJH1-51B showed degradation of the BRD4 short isoform starting at 6 h with progressive BRD4 loss through 24 h **(Figure S5c)**. Despite the non-reactive SJH1-51B-NC showing no BRD4 degradation, we observed equivalent binding of both SJH1-51B and SJH1-51B-NC to SKP1 in the SKP1-FBXO7-CUL1-RBX1 complex **(Figure S5d)**. Akin to what we observed with EN884 and SJH1-37-m, we did not observe binding of either compound to SKP1 alone **(Figure S5e)**.

### Using Covalent SKP1 Recruiter in PROTACs to Degrade Androgen Receptor

BRD4 is a relatively easy protein to degrade with PROTACs. As such, we next tested whether our covalent SKP1 recruiter could be used to degrade another neo-substrate protein—the androgen receptor (AR). We synthesized 4 AR PROTACs linking our covalent SKP1 recruiter to the Arvinas AR PROTAC-targeting ligand from ARV-110 with either no linker or a C4, C6, or C7 alkyl linker to yield SJH1-63, SJH1-62C, SJH1-62D, and SJH1-62B, respectively **(Figure 5a-5d)** ^30^. We observed AR degradation with all 4 PROTACs, wherein SJH1-62B with the C7 alkyl linker showed the most potent and robust AR degradation in LNCaP prostate cancer cells **(Figure 5a-5d)**. We further demonstrated that the SJH1-62B-mediated loss of AR was attenuated upon proteasome inhibitor pre-treatment **(Figure 5e)**. Quantitative proteomic profiling also demonstrated selective degradation of AR with 11 other proteins also significantly lower in levels among >5000 proteins quantified **(Figure 5f; Table S5)**. We synthesized a non-reactive analog of SJH1-62B, SJH1-62B-NC, and surprisingly observed equivalent AR degradation as SJH1-62B in LNCaP cells **(Figure S6a-S6b)**. As was observed with the BRD4 degraders, we found that both SJH1-62B and SJH1-62B-NC both bound equally to SKP1 in the SKP1-FBXO7-CUL1-RBX1 complex, but not to SKP1 alone **(Figure S6c)**.

**Figure 5.**
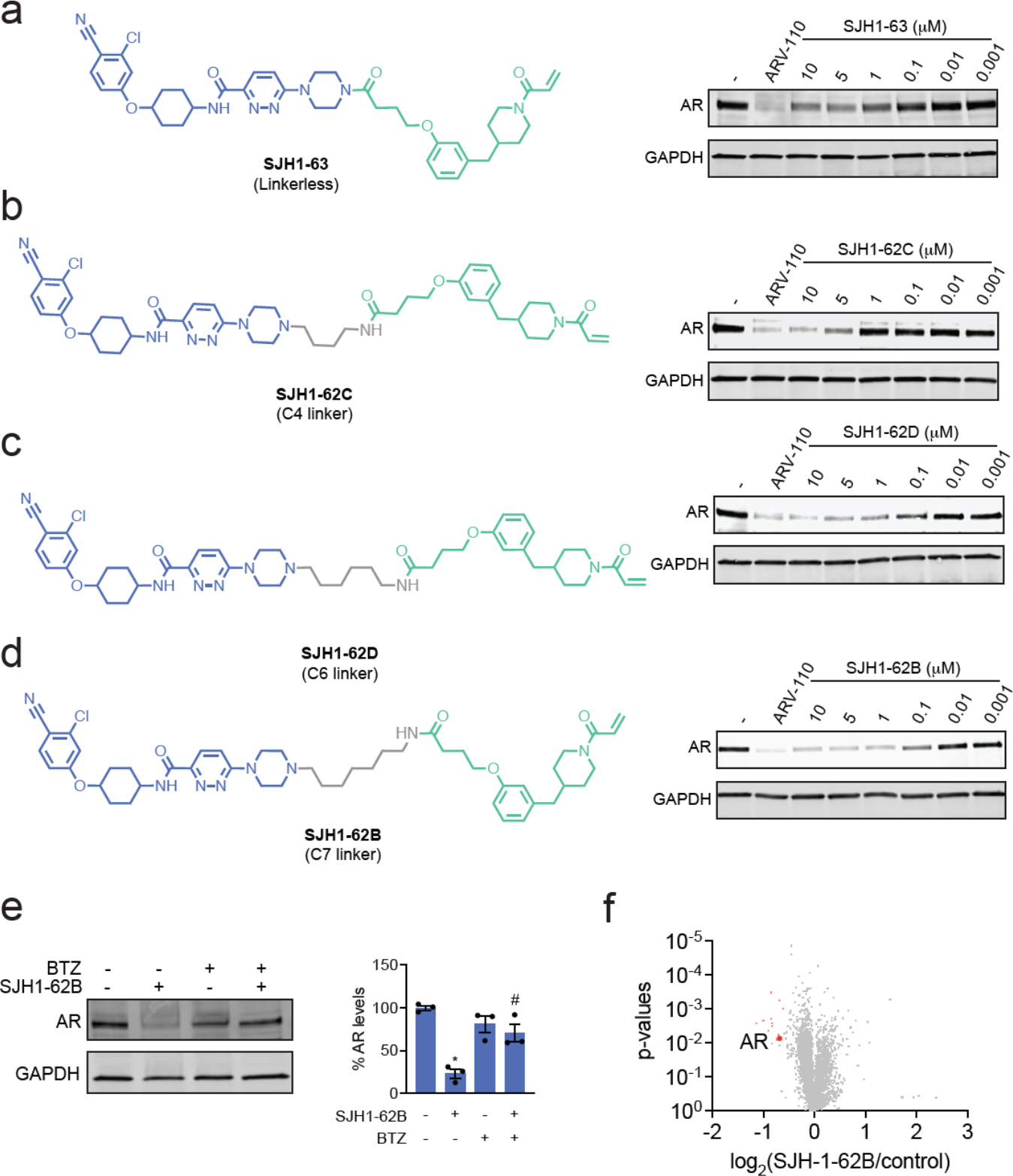
SKP1-based androgen receptor (AR) degraders. **(a-d)** Structures of AR degraders consisting of SKP1 covalent recruiters in green, linker in grey, and an AR-targeting ligand from ARV-110 in blue with either no linker, C4, C6, or C7 alkyl linkers. Also shown are AR degradation by each degrader in LNCaP prostate cancer cells. LNCaP cells were treated with DMSO vehicle, ARV-110 (1 μM), or each degrader for 24 h. AR and loading control GAPDH levels were assessed by Western blotting. **(e)** Proteasome-dependence of BRD4 degradation. LNCaP cells were pre-treated with DMSO vehicle or bortezomib (1 μM) for 1 h prior to treatment of cells with DMSO vehicle or SJH1-62B (1 μM) for 24 h. AR and loading control GAPDH levels were assessed by Western blotting and quantified. **(f)** TMT-based quantitative proteomic profiling of SJH1-62B in LNCaP cells. LNCaP cells were treated with DMSO vehicle or SJH1-62B (1 μM) for 24 h.

## Conclusions

Here, we discovered a SKP1 recruiter that covalently targets an intrinsically disordered C160 of SKP1 that sits at the interface of SKP1 and the CUL1 F-box substrate receptor. This observation represents a proof of concept for discovering ligandable sites that exclusively emerge when an IDR containing protein forms a complex with its interacting protein. We further demonstrated that this covalent SKP1 ligand can be used in PROTAC applications to degrade neo-substrates such as BRD4 and AR in cells. Given the essentiality of SKP1, covalent recruitment of SKP1 may potentially overcome resistance mechanisms of PROTACs that utilize non-essential E3 ligases in cancer treatment.

While we demonstrate proof-of-concept of using Cullin-RING E3 ligase adaptor proteins in PROTACs, our SKP1 ligand is still an early hit and will require significant medicinal chemistry optimization to improve potency, selectivity, and drug-like properties. Intriguingly, we observed that both the covalent and non-covalent BRD4 and AR PROTACs both bound to SKP1 in the SKP1-FBXO7-CUL1-RBX1 complex, suggesting that the SKP1 ligand may be a reversible binding component. While BRD4 degradation was not observed with the non-reactive degrader, we did observe AR degradation with the non-reactive AR degrader. We do not yet understand this discrepancy, but it may be that a tighter ternary complex may be necessary for BRD4 degradation compared to that of AR. Nonetheless, SKP1 represents another core component of the ubiquitin-proteasome machinery that can be exploited to degrade neo-substrate proteins. Other recently published examples include direct recruitment of the proteasome, recruitment of E2 ubiquitin conjugating enzymes such as UBE2D, as well as the CUL4 adaptor protein DDB1. Our results also highlight the utility of using covalent chemoproteomic approaches to identify recruiters for proteins within the ubiquitin proteasome system that can be utilized for targeted protein degradation applications.

## Acknowledgement

We thank the members of the Nomura Research Group and Bristol Myers Squibb for critical reading of the manuscript. This work was supported by Bristol Myers Squibb for all listed authors. This work was also supported by the Nomura Research Group and the Mark Foundation for Cancer Research ASPIRE Award for DKN. This work was also supported by grants from the National Institutes of Health (R01CA240981 and R35CA263814 for DKN) and the National Science Foundation Molecular Foundations for Biotechnology Award (2127788). We also thank Drs. Hasan Celik and UC Berkeley’s NMR facility in the College of Chemistry (CoC-NMR) for spectroscopic assistance. Instruments in the College of Chemistry NMR facility are supported in part by NIH S10OD024998.

## Author Contributions

SH, DKN, IEW conceived of the project idea. SH, DKN designed experiments, performed experiments, analyzed and interpreted the data, and wrote the paper. SH, DKN, AO, OWH performed experiments, analyzed and interpreted data, and provided intellectual contributions.

## Methods

### Covalent ligand library

Compound names that start with “EN” were purchased from Enamine.

### Gel-Based ABPP

Recombinant Human SKP1/FBXO7/CUL1/RBX1 Complex Protein, CF (Boston Biochem, E3-526) was employed for the initial Gel-Based ABPP screening. The SKP1 containing SCF complex was reconstituted by utilizing stoichiometric amount of the following recombinant proteins: Recombinant Human CUL1/RBX1 Neddylated Complex Protein (R&D systems, E3-411-025), Recombinant Human FBXO7 GST (N-Term) Protein (Novus Biologicals, H00025793-P01-2ug) /or/ Human Recombinant FBXO2 Protein (Novus Biologicals™, NBP2517640.05MG), and Recombinant Human SKP1 (Bristol Myers Squibb). SKP1 or SKP1 containing SCF complex (0.1 μg) in phosphate-buffered saline (PBS) was incubated with a covalent ligand (50 μM) or DMSO vehicle. After incubating at 23 °C for 1 hour, 0.1 μM of IA-rhodamine (Setareh Biotech, 6222) was subsequently added. The reaction mixture was incubated at 23 °C for an additional 30 min and quenched by adding 4x reducing Laemmli SDS sample loading buffer (Thermo Scientific™, J160015.AD). Samples were boiled at 95 °C for 5 minutes and separated on precast 4-20% Criterion TGX gels (Bio-Rad). The competition of IA-rhodamine was analyzed by in-gel fluorescent signal using a ChemiDoc MP (Bio-Rad). Imaged gels were stained using Pierce™ Silver Stain Kit (Thermo Scientific™, 24612) following manufacturer’s instruction.

### Cell Culture

HEK293T, LNCaP, and MDA-MB-231 cell lines were sourced from the UC Berkeley Cell Culture Facility. HEK293T and MDA-MB-231 cells were cultured in Dulbecco’s Modified Eagle Medium (DMEM) supplemented with 10% (v/v) fetal bovine serum (FBS) and 2.5 mM Gibco L-Glutamine (Q). LNCaP cells were cultured using Roswell Park Memorial Institute 1640 (RPMI-1640) medium with 10% (v/v) FBS and 2.5 mM Q. All cells were maintained in a standard incubation environment maintaining 37 °C with 5% CO_2_.

### Preparation of Cell Lysates

Cells were gently washed with cold PBS twice and subsequently scraped. The scraped cells were pelleted by centrifugation (1400 g, 4 min, 4 °C). After removing the supernatant, the pelleted cells were re-suspended in PBS containing protease inhibitor (cOmplete™, EDTA-free protease inhibitor Cocktail). Cells lysis was achieved via sonication, and cellular debris were separated using centrifugation (7500 g, 10 min, 4 °C). Cell lysates were then transferred to new tubes and the concentration of proteome was determined using Pierce™ BCA Protein Assay kits (Thermo Scientific™, 23227). Next, the cell lysates were further diluted to appropriate experiment condition. For the western blotting experiments, the pelleted scraped cells were re-dissolved in the lysis buffer (Thermo Scientific™, 89900). After incubating at 4 °C for 30 minutes with occasional vertexing, cellular debris was pelleted by centrifugation (7500 g, 10 min, 4°C). The supernatant was collected, and the proteome concentration was verified using BCA assay and further diluted to match the required experimental conditions.

### Western Blotting

Proteins were separated by precast 4-20% Criterion TGX gels (Bio-Rad) and transferred to nitrocellulose membrane using the Trans-Blot Turbo transfer system (Bio-Rad). After the transfer, membrane was blocked with Tris-buffered saline containing Tween 20 (TBST) containing 5% bovine serum albumin (BSA) for 2 hours at 23 °C. After the blocking, targeted proteome was probed with primary antibody in TBST with 5% BSA (Primary antibody dilution condition from manufacturer). Incubation with primary antibody was performed overnight (10 hours) in cold room (4 °C), and the primary antibody was washed three times with TBST. Membrane was then incubated in the dark with IR680 or IR800-conjugated secondary antibodies (1: 10000 dilution). After 1 hour incubation at 23 °C, blot was washed 3 times with TBST, and the blots were visualized using Odyssey Li-Cor fluorescent scanner. Following antibodies were used for this study: GAPDH (Proteintech, 60004-1-1G-150L), BRD4 (Abcam, ab128874), androgen receptor (Abcam, ab133273), SKP1 (Millipore-Sigma, SAB5300376-100UL).

### Pulldown of SKP1 Using SJH1-37m

HEK293T cells at 80% confluency were treated with DMSO and SJH1-37m (50 uM). After 4 hours of incubation, cells were harvested, and lysates were prepared as previously described. Each lysate was normalized to a concertation of 5 mg/mL, and 500 μL of each lysate was transferred to separate tube. To each tube containing 500 μL of cell lysate, 10 μL of 20 mM biotin picolyl azide (Sigma Aldrich, 900912) in DMSO, 10 μL of 50 mM TCEP in H_2_O, 10 μL of 50 mM CuSO_4_ in H_2_O, and 30 μL of TBTA ligand (1.3 mg/mL in 1:4 DMSO/tBuOH, Cayman chemical, 18816) were added. The reaction mixture was incubated at 23 °C for 90 minutes, and the reaction was quenched by proteins precipitation. After washing the protein pellets twice with cold MeOH (4 °C), the pellets were redissolved in 200 μL of 1.2 % SDS/PBS (w/v). After heating the samples at 95 °C for 10 minutes, 10 μL of each sample was set aside for the western blot analysis (input control). 1 mL of PBS was added to the remaining sample to reduce the total SDS concentration to less than 0.2 % SDS/PBS (w/v).

100 μL of streptavidin agarose beads (ThermoFisher, 20353) were added to the lysates containing tubes and the samples were incubated at 4 °C on a rotator for 10 hours. After incubation, samples were brought to 23 °C, and beads were spined down in a centrifuge (1300 g, 2 min). The supernatant was removed. and beads were washed three more times with 500 μL of PBS and 500 μL of H_2_O. The beads were then suspended in 30 μL of Laemmli SDS sample loading buffer and heated to 95 °C for 10 minutes. Along with the input control, the resulting samples were subjected to western blot analysis.

### Mapping of EN-884 Site of Modification on SKP1 SCF Complex by LC-MS/MS

SKP1 containing SCF complex (100 μg, reconstituted as described) was diluted in PBS (100 μL) and incubated with EN-884 (50 μM) for 90 minutes at 37 °C. The protein was then precipitated by adding 900 μL of acetone, and the precipitation continued at -20 °C for 2 hours. The sample was pelleted at 20,000 g for 10 minutes, 4 °C, and the supernatant was carefully aspirated. The precipitated pellet was washed with 200 μL of acetone. The sample was then resuspended in 30 μL of 8 M Urea and 30 μL of ProteaseMax surfactant (20 μg/mL in 100 mM ammonium bicarbonate, Promega, V2071).

After vigorous mixing for 30 seconds, 40 μL of 100 mM ammonium bicarbonate buffer was added. To the sample containing tube, 10 μL of 110 mM TCEP in H_2_O was added and the sample was incubated at 60 °C for 30 minutes. Next, 10 μL of 150 mM iodoacetamide in H_2_O was added, and the sample was incubated at 37 °C for 30 minutes. The resulting sample was diluted with 120 μL of PBS containing 1.2 μL of ProteaseMax surfactant (0.1 mg/mL in 100 mM ammonium bicarbonate). Treated proteins were then digested with sequencing grade modified trypsin (Promega, V5111) and incubated overnight (37 °C, 14 hours). Digestion was quenched at the following day by adding 25 μL of formic acid, and the sample was fractionated using high pH reversed-phase peptide fractionation kits (ThermoFisher, 84688) following manufacturer’s recommendation.

### IsoDTB-ABPP Cysteine Chemoproteomic Profiling and TMT Proteomic Profiling Methods

Methods are described in **Supporting Information**

