## Supporting Information for "Exploiting the Cullin E3 Ligase Adaptor Protein SKP1 for Targeted Protein Degradation"

**Supporting Table Legends**

**Table S1. Structures of compounds screened and screening data.** Shown are the structures of all the covalent ligands screened against SKP1. Cysteine-reactive covalent ligands (50 μM) were screened against a rhodamine-functionalized iodoacetamide (IA-rhodamine) labeling of pure SKP1 protein in the SKP1-FBXO7-CUL1-RBX1 complex. Individual gels are shown in **Figure S1.** Values are noted as percent of labeling compared to that of DMSO vehicle control.

**Table S2. Cysteine chemoproteomic profiling of EN884.** isoDTB-ABPP analysis of EN884 in HEK293T cells. HEK293T cells were treated with DMSO vehicle or EN884 (50 μM) for 1 h, after which lysates were labeled with an alkyne-functionalized iodoacetamide probe (IA-alkyne) and control and treated cells were subjected to CuAAC with a desthiobiotin-azide with an isotopically light or heavy handle, respectively. After the isoDTB-ABPP procedure, probe-modified peptides were analyzed by LC-MS/MS and control vs treated probe-modified peptide ratios were quantified. Individual probe-modified peptides, proteins, light/heavy ratios, and statistical analyses are shown in the table.

**Table S3. TMT-based quantitative pulldown of SJH1-37m probe.** SKP1 pulldown using SJH1-37m probe. HEK293T cells were treated with DMSO vehicle or SJH1-37m (50 μM) for 4 h. Lysates were subjected to copper-catalyzed azide-alkyne cycloaddition (CuAAC) to append on a biotin enrichment handle after which probe-modified proteins were avidin-enriched, digested with trypsin, and analyzed by TMT-based quantitative proteomics.

**Table S4. TMT-based quantitative proteomic profiling of SJH1-51B in MDA-MB-231 cells.** MDA-MB-231 cells were treated with DMSO vehicle or SJH1-51B (10 μM) for 24 h.

**Table S5. TMT-based quantitative proteomic profiling of SJH1-62B in LNCaP cells.** LNCaP cells were treated with DMSO vehicle or SJH1-62B (1 μM) for 24 h.

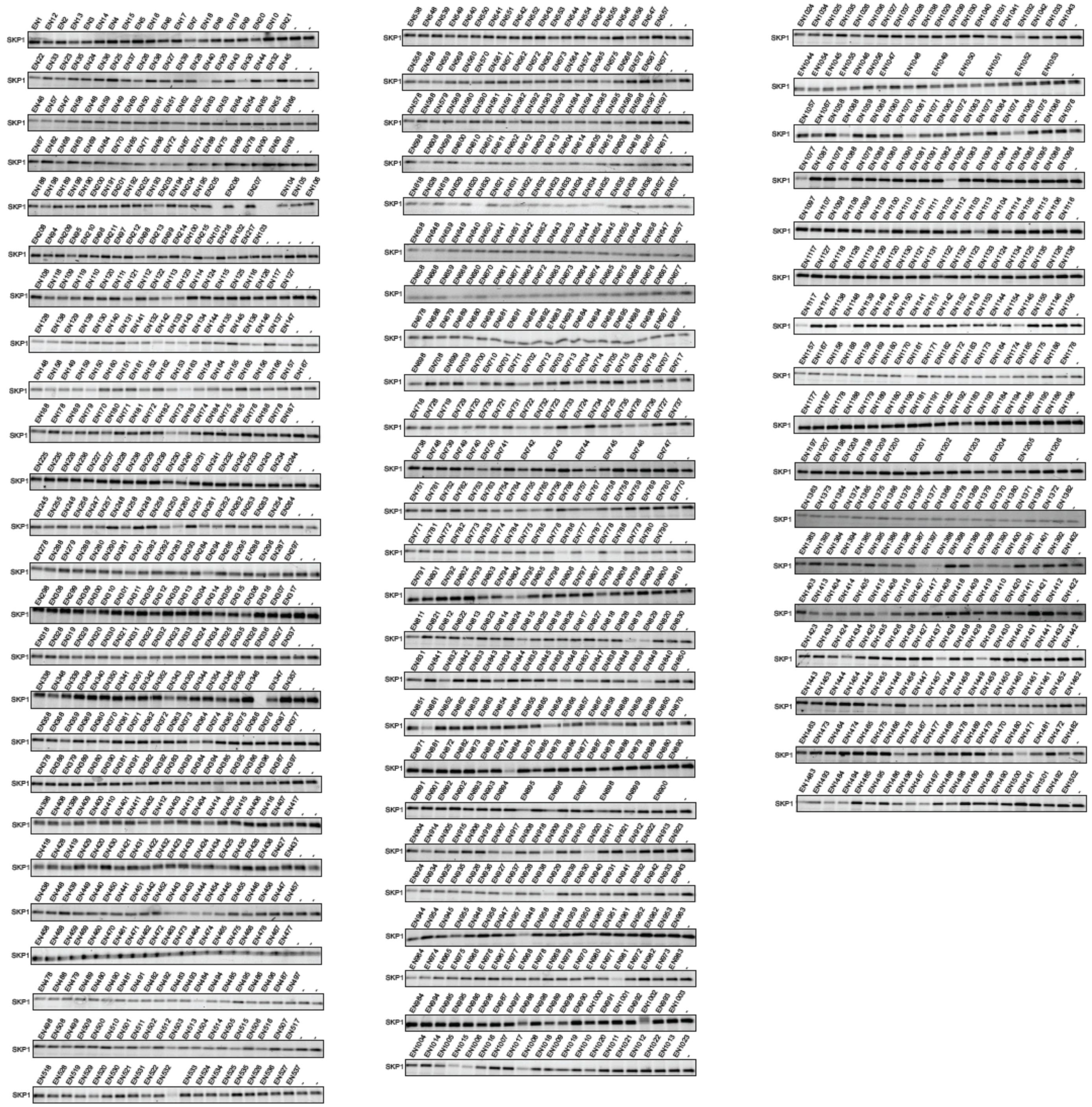

**Figure S1. Gel-based ABPP screen of cysteine-reactive covalent ligands against SKP1.** Cysteine-reactive covalent ligands (50 μM) were screened against a rhodamine-functionalized iodoacetamide (IA-rhodamine) labeling of pure SKP1 protein in the SKP1-FBXO7-CUL1-RBX1 complex. Proteins were separated by SDS/PAGE and IA-rhodamine labeling was read out by in-gel fluorescence.

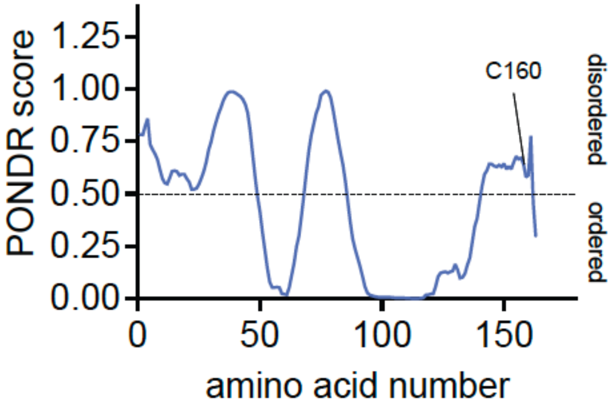


**Figure S2. Predictor of Natural Disordered Regions (PONDR[(R)]) prediction of order and disorder of the full-length human SKP1 amino acid sequence using http://www.pondr.com.**

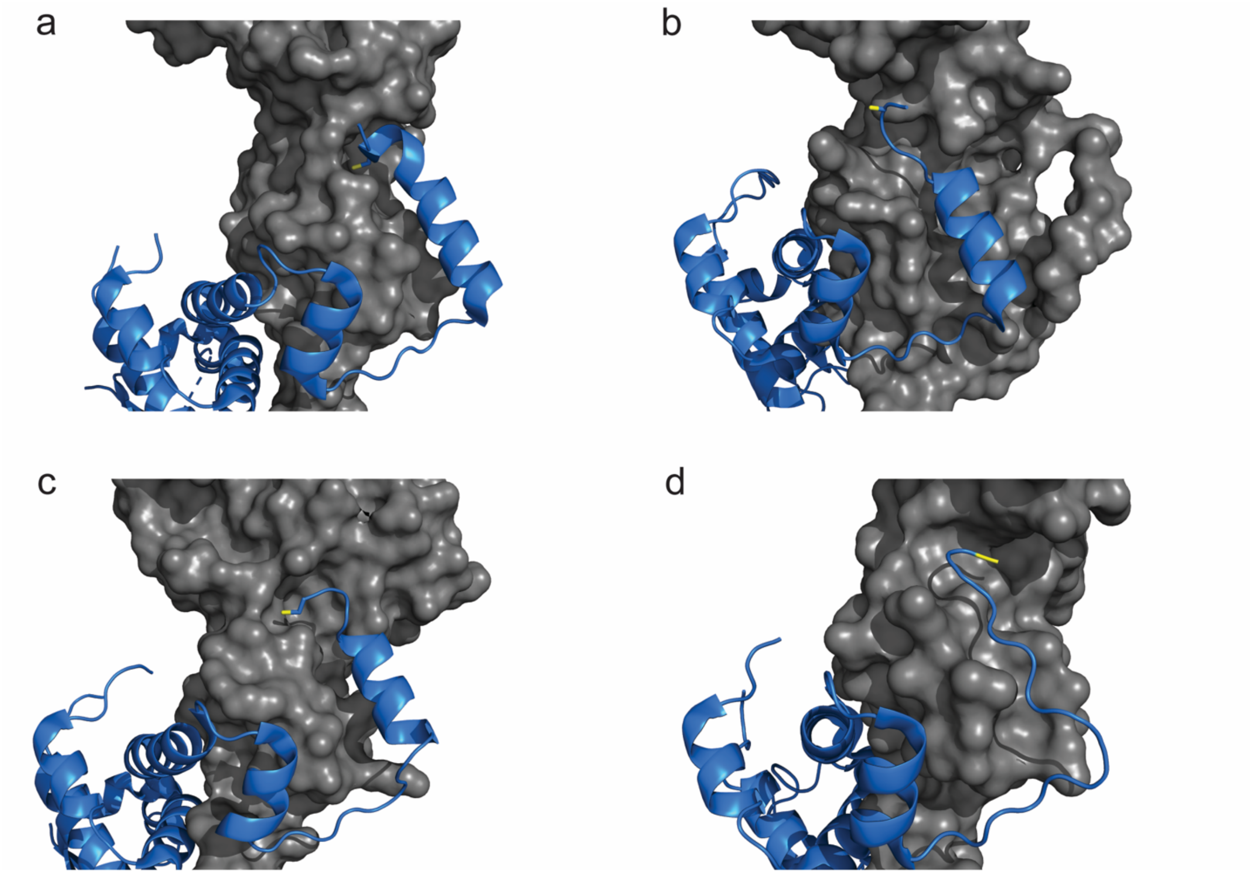


**Figure S3. Structure of protein complex highlighting conformational change in the C terminus of SKP1 upon the SCF complex formation. (a)** Crystal structure of Skp1-FBG3 (PDB ID:3WSO). FBG3 is shown in surface representation (grey) and Skp1 is shown in cartoon representation (blue)[1]. C160 of Skp1 is highlighted with yellow color. **(b)** Crystal structure highlighting Skp1-FBXW7 (PDB ID:5V4B)[2]. FBXW7 is shown in surface representation (grey) and Skp1 is shown in cartoon representation (blue). C160 of Skp1 is highlighted with yellow color. **(c)** Crystal structure of Skp1-Skp2 (PDB ID:2AST) [3]. Skp2 is shown in surface representation (grey) and Skp1 is shown in cartoon representation (blue). C160 of Skp1 is highlighted with yellow color. **(d)** Crystal structure of Skp1-FBXO2 (PDB ID:2E31)[4]. FBXO2 is shown in surface representation (grey) and Skp1 is shown in cartoon representation (blue). C terminus of Skp1 is highlighted with yellow color.

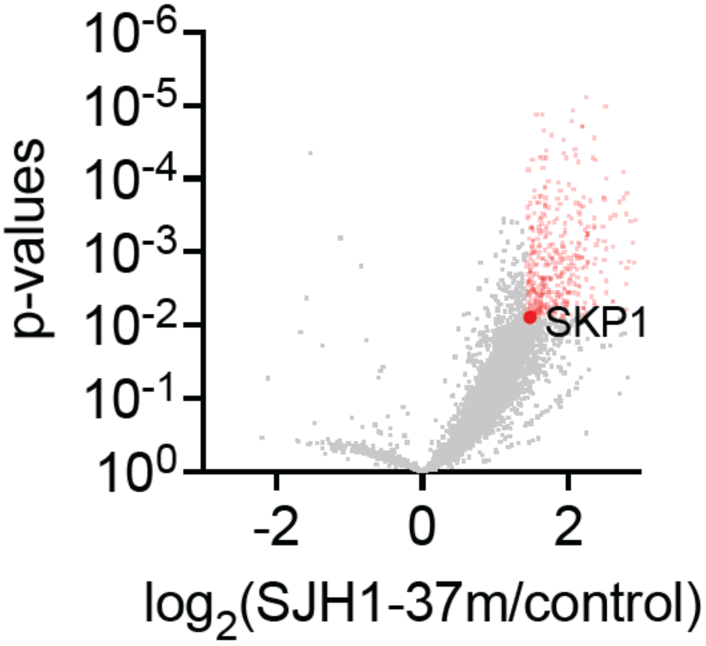


**Figure S4. SJH1-37m pulldown of SKP1.** SKP1 pulldown using SJH1-37m probe. HEK293T cells were treated with DMSO vehicle or SJH1-37m (50 μM) for 4 h. Lysates were subjected to copper-catalyzed azide-alkyne cycloaddition (CuAAC) to append on a biotin enrichment handle after which probe-modified proteins were avidin-enriched, digested with trypsin, and analyzed by TMT-based quantitative proteomics.

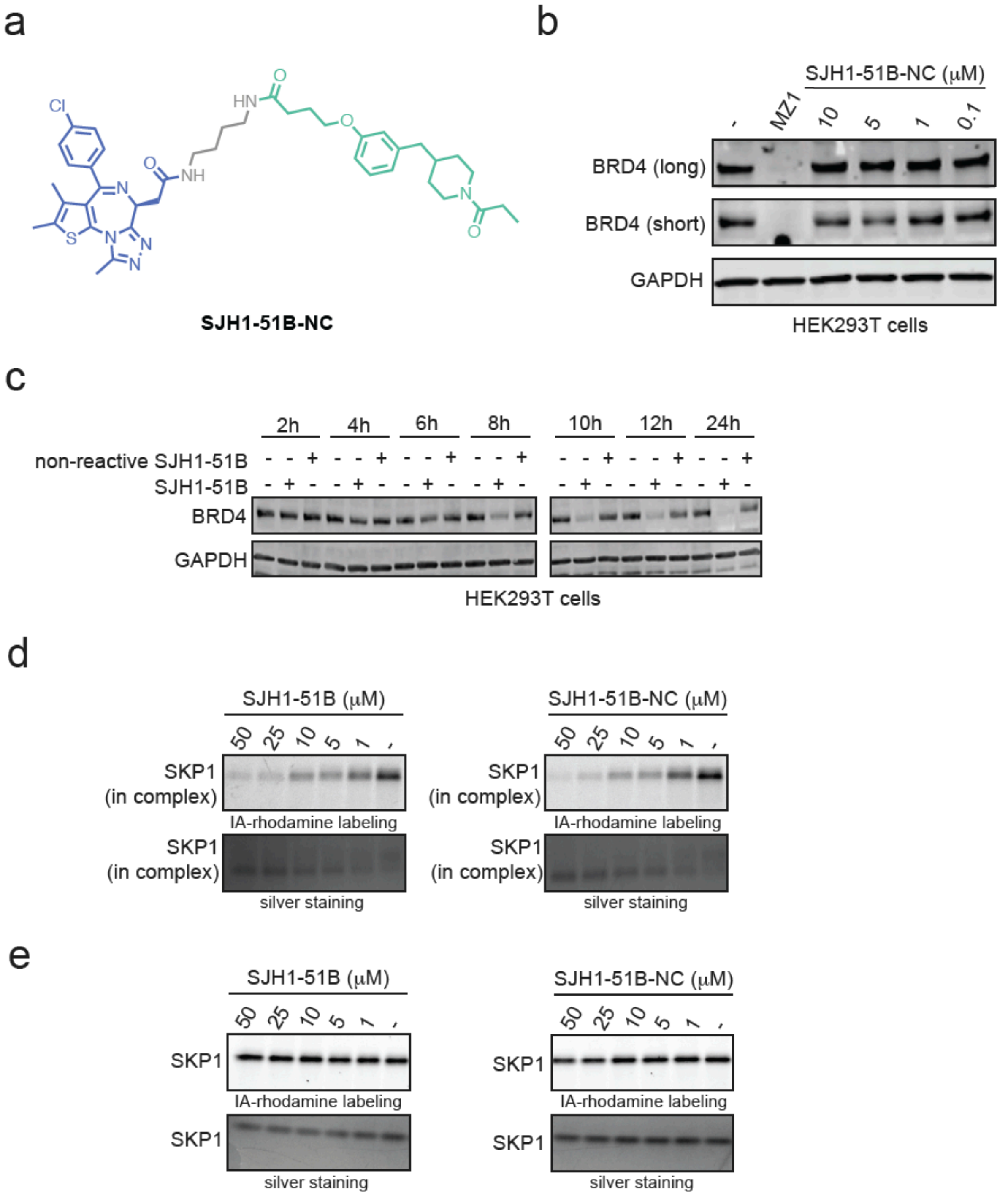


**Figure S5. Characterization of BRD4 degraders and their non-reactive analogs. (a)** Structure of non-reactive BRD4 degrader SJH1-51B-NC. **(b)** Lack of BRD4 degradation with SJH1-51B-NC. HEK293T cells were treated with DMSO vehicle, MZ1 (1 μM), or SJH1-51B-NC for 24 h and long and short BRD4 isoforms and loading control GAPDH were assessed by Western blotting. **(c)** Time course of BRD4 degraders. HEK293T cells were treated with DMSO vehicle or SJH1-51B, or SJH1-51B-NC for the noted time points and short BRD4 isoform levels and loading control GAPDH were assessed by Western blotting. **(d, e)** Gel-based ABPP of SJH1-51B or SJH1-51B-NC against IA-rhodamine labeling of either SKP1 in the SKP1-FBXO7-CUL1-RBX1 complex or with SKP1 alone. SKP1 in complex or SKP1 alone was pre-incubated with DMSO vehicle or degraders for 1 h prior to labeling with IA-rhodamine (100 nM) for 30 min, after which proteins were separated by SDS/PAGE and IA-rhodamine labeling was visualized by in-gel fluorescence and protein loading was assessed by silver staining. Blots and gels **(b, d,** and **e)** are representative of n=3 biologically independent replicates/group.

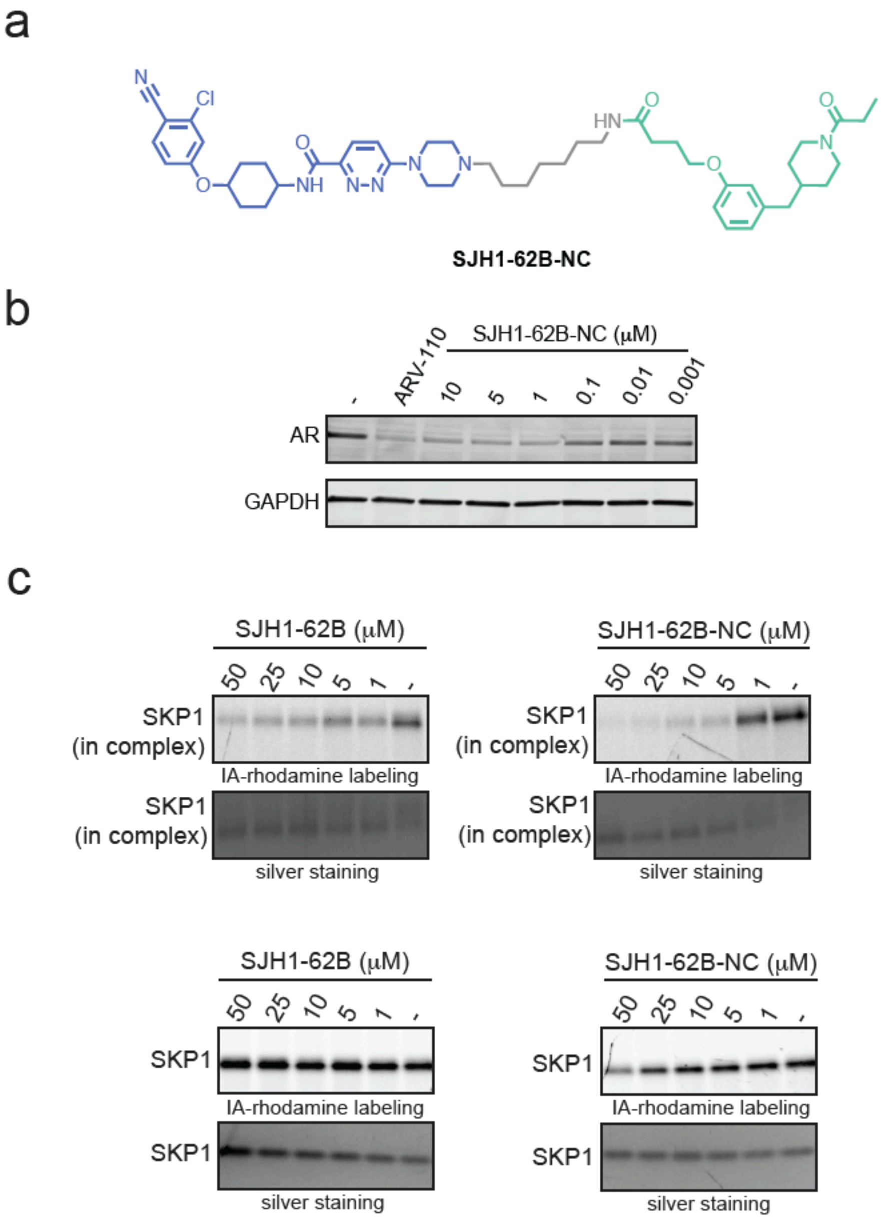


**Figure S6. Characterization of AR degraders and their non-reactive analogs. (a)** Structure of non-reactive BRD4 degrader SJH1-62B-NC. **(b)** AR degradation with SJH1-62B-NC. LNCaP cells were treated with DMSO vehicle, ARV-110 (1 μM), or SJH1-62B-NC for 24 h and AR and loading control GAPDH levels were assessed by Western blotting. **(c)** Gel-based ABPP of SJH1-62B or SJH1-62B-NC against IA-rhodamine labeling of either SKP1 in the SKP1-FBXO7-CUL1-RBX1 complex or with SKP1 alone. SKP1 in complex or SKP1 alone was pre-incubated with DMSO vehicle or degraders for 1 h prior to labeling with IA-rhodamine (100 nM) for 30 min, after which proteins were separated by SDS/PAGE and IA-rhodamine labeling was visualized by in-gel fluorescence and protein loading was assessed by silver staining. Blots and gels **(b, c)** are representative of n=3 biologically independent replicates/group.

### Methods

#### IsoDTB-ABPP cysteine chemoproteomic profiling of EN884

Cells were treated when they reached 80% confluency, with either DMSO vehicle or 50 μM of EN884 for 4 hours. Cell lysates were prepared as it is described, and proteome concentration was normalized to 2.5 mg/mL. For each biological replicate, two 1 mL aliquots of 2.5 mg/mL were used. Prior to tag labeling, each aliquot was treated with 20 μL of IA-alkyne (200 μM) for 1 hour in 23 °C. Subsequently, each sample set was treated with 120 μL of either heavy or light isoDTB tags containing master mix for 90 minutes at 23 °C. Master mix contains 1020 μL of TBTA (0.9 mg/mL in 1:4 in DMSO/tBuOH), 330 μL of $CuSO_4$ (12.5 mg/mL in $H_2O$), 330 μL of of TCEP (14.0 mg/mL in $H_2O$), and 160 μL of isoDTB tag (4 mg of either heavy or light tag in DMSO, Click chemistry Tools, 1565). After the reaction, one heavy tagged and one light tagged labeled samples were combined and precipitated in acetone at –20 °C.

After the overnight precipitation, the sample was centrifuged at 7,000 g for 10 minutes in 4 °C. The supernatant was removed, and the precipitated protein samples were resuspended in cold MeOH. Protein pellets were washed with cold MeOH (500 μL, 3 times) and dissolved in 600 μL of 8 M urea in PBS buffer. Pellets were completely dissolved in the urea solution by sonication, and the urea concentration was adjusted to 2 M by adding 1800 μL of PBS. Two replicates were then combined into a single container, and the combined sample sets were further diluted with 2400 μL of PBS containing 0.2% NP40 (w/v). Proteome was then incubated with high-capacity streptavidin agarose beads (200μL/sample, ThermoFisher, 20357) for 4 hours at 4 °C with an occasional mixing.

After incubation, the beads were centrifuged for 1 minute and the supernatant were removed. The remaining beads were then washed in following order: 3 x PBS with 0.1% NP40, 3 x PBS, and 3 x $H_2O$. Next, the protein-bound beads were resuspended in 400 μL of 2 M Urea in 0.1 M of tetraethylammonium bromide (TEAB) buffer. 8 μL of trypsin (0.5 mg/mL) was then added, and the bead bound proteins were digested for 14 hours at 37 °C. The resulting samples were then diluted with 800 μL of 0.1% NP40 in PBS, and the beads were washed with the same buffer three times and three more times with $H_2O$. Bead-bound peptides were then eluted with 0.1% formic acid in 50% acetonitrile (v/v). Eluted samples were dried using a vacufuge and re-dissolved in 300 μL of $H_2O$ with 0.1% TFA. Suspended samples were then fractionated using high pH reversed-phase peptide fractionation kits (ThermoFisher, 84688).

Data were extracted in the form of MS1 and MS2 files using Raw Converter (Scripps Research Institute) and searched against the Uniprot human database using ProLuCID search methodology in IP2 v.3-v.5 (Integrated Proteomics Applications, Inc.) [5]. Cysteine residues were searched with a static modification for carboxyaminomethylation (+57.02146).

For isoDTB-ABPP, we also searched for up to two differential modifications for methionine oxidation and either the light or heavy isoDTB tags (+561.33872 or +567.34621, respectively). Peptides were required to be fully tryptic peptides. ProLUCID data were filtered through DTASelect to achieve a peptide false-positive rate below 5%. Only those probe-modified peptides that were evident across three out of three biological replicates were interpreted for their isotopic light to heavy ratios. Light versus heavy isotopic probe-modified peptide ratios are calculated by taking the mean of the ratios of each replicate paired light versus heavy precursor abundance for all peptide-spectral matches associated with a peptide. The paired abundances were also used to calculate a paired sample *t*-test *P* value in an effort to estimate constancy in paired abundances and significance in change between treatment and control. *P* values were corrected using the Benjamini–Hochberg method.

#### TMT-based quantitative proteomic profiling

At 75% of their confluency, cells were treated with either DMSO vehicle or bifunctional compound of interest (SJH1-51B, 10 μM for MDA-MB-231 or SJH1-62B, 1 μM in LNCaP). After 24 hours of treatment, cell lysates were prepared as described. From each sample set, 100 μg of proteome was reduced, alkylated, and enzymatically digested using sequencing grade modified trypsin (Promega, V5111). The digestion took place overnight at 37 °C. Subsequently, each set of samples was subjected to labeling with TMTsixplex isobaric tag (ThermoFisher, 90061). 25 μg of each labeled set was combined, dried, and resuspended to fractionate. High pH reversed-phase peptide fractionation kits (ThermoFisher, 84868) were used based on the manufacturer's protocol to prepare each sample. Dried fractions were then resuspended in 25 μL of 0.1 % Formic acid/$H_2O$ (w/v) to be analyzed by LC-MS/MS.

Trypsin cleavage specificity was defined, allowing for up to 2 missed cleavages (cleavage at K, R except if followed by P). For the static modifications, carbamidomethylation of cysteine and methionine oxidation were assigned. Variable modifications were designated for N-termini and lysine residue. Reporter ion calculations were determined using summed abundances with the most confident centroid selected from 20 ppm window. Only peptide-to-spectrum matches that are unique assignments to a given identified protein with the total dataset are considered for protein quantitation. High confidence protein identification were reported with false discovery rate (FDR) cut-off set to < 1%. Differential abundance significance was assessed using ANOVA with Benjamini-Hochberg correction, yielding p-values for interpretation.

**SJH1-37-m alkyne pulldown quantitative proteomics.**
Cells were treated with either compound (SJH1-37-m, 50 µM) or DMSO vehicle at 75% of confluency for 4 hours. After treatment, cells were harvested, lysed, and the proteome concentration was adjusted to 5 mg/mL in 500 µL of PBS using the BCA assay. To each tube containing cell lysate, the following reagnets were added: 10 µL of 20 mM biotin picolyl azide (Sigma Aldrich, 900912) in DMSO, 10 µL of 50 mM TCEP in H2O, 10 µL of 50 mM $CuSO_4$ in H2O, and 30 µL of TBTA ligand (1.3 mg/mL in 1:4 DMSO/tBuOH, Cayman Chemical, 18816). The reaction mixture was incubated at 23 °C for 90 minutes, and the reaction was quenched by protein precipitation. Precipitated pellets were washed using 500 µL of MeOH, combined, and centrifuged again to yield 10 mg of white pellets per sample. Combined samples were resuspended in 1.2% SDS-PBS (1 mL), completely dissolved, and heated to 90 °C for 10 minutes. The soluble proteome was then diluted with 5 mL of PBS and further incubated with high-capacity streptavidin-agarose beads (200 µL/sample, ThermoFisher Scientific, 20357). Beads and lysates were incubated overnight at 4 °C with rotation. On the following day, beads were suspended and washed three times with 0.1% SDS-PBS, PBS, and H2O. Washed beads were resuspended in 6 M Urea/PBS (500 µL), and the samples were further treated with DTT and iodoacetamide. After removing the supernatant, beads were resuspended in 100 µL of 50 mM TEAB and enzymatically digested overnight using sequencing-grade trypsin (Promega, V5111). Digested peptides were eluted through centrifugation and subsequently with 500 µL of 50% ACN-H2O. The collected peptides containing samples were dried using a vacufuge and resuspended in 100 µL of 50 mM TEAB before labeling using commercially available TMTsixplex tags (ThermoFisher, P/N 90061). After labeling, 30 µg of each labeled sample was combined and dried using a vacufuge. Dried samples were redissolved with 300 µL of 0.1% TFA in H2O and further fractionated using high-pH reversed-phase peptide fractionation kits (ThermoFisher, P/N 84868) following the manufacturer's protocol. Analysis by LC-MS/MS and following procedure was performed as it is described for TMT-based quantitative proteomic profiling.

**Mass Spectrometry Analysis**
Mass spectrometry analysis was performed on an Orbitrap Eclipse Tribrid Mass Spectrometer with a High Field Asymmetric Waveform Ion Mobility (FAIMS Pro) Interface (Thermo Scientific) with an UltiMate 3000 Nano Flow Rapid Separation LCnano System (Thermo Scientific). Off-line fractionated samples (5 µl aliquot of 15 µl sample) were injected via an autosampler (Thermo Scientific) onto a 5 µl sample loop which was subsequently eluted onto an Acclaim PepMap 100 C18 HPLC column (75 µm x 50 cm, nanoViper). Peptides were separated at a flow rate of 0.3 µl/min using the following gradient: 2 % buffer B (100 % acetonitrile with 0.1 % formic acid) in buffer A (95:5 water:acetonitrile, 0.1 % formic acid) for 5 min, followed by a gradient from 2 to 40 % buffer B from 5 to 159 min, 40 to 95 % buffer B from 159 to 160 minutes, holding at 95 % B from 160-179 min, 95 % to 2 % buffer B from 179 to 180 min, and then 2 % buffer B from 180 to 200 min. Voltage applied to the nano-LC electrospray ionization source was 2.1 kV. Data was acquired through an MS1 master scan (Orbitrap analysis, resolution 120,000, 400-1800 m/z, RF lens 30 %, heated capillary temperature 250 °C) with dynamic exclusion enabled (repeat count 1, duration 60 s). Data-dependent data acquisition comprised a full MS1 scan followed by sequential MS2 scans based on 2 s cycle times. FAIMS compensation voltages (CV) of -35, -45, and -55 were applied. MS2 analysis consisted of: quadrupole isolation window of 0.7 m/z of precursor ion followed by higher energy collision dissociation (HCD) energy of 38 % with a orbitrap resolution of 50,000.

**Plasmid isolation**

Predesigned shRNA construct of SKP1 (The MISSION™, TRCN0000284791) was purchased as bacterial glycerol stock. Targeted clones containing E. coli were cultured, and the targeted plasmid were collected using QIAprep Spin Miniprep Kit by following the manufacturer's protocol (QIAGEN, ID: 27104).

**SKP1 Lentiviral Knockdown Studies**

For each replicate, two 1.5 mL tubes were used. To the first tube, 2.5 μg of following lentiviral plasmids – shRNA construct of SKP1, psPAX2 (carries GAG, REV, polgenes) and pMD2G (carries VSVG pseudotyping gene) – were dissolved in 750 μL of Gibco™ Opti-MEM™ I Reduced Serum Medium (catalogue no. 31985-062). To the second tube, lipofectamine™ 2000 (Invitrogen™, 11668019) was diluted in 750 μL of Gibco™ Opti-MEM™ I Reduced Serum Medium. Each tube was sat undisturbed for 5 minutes, and the two were mixed into one. After 30 minutes of incubation at a room temperature without mixing, each combined tube was added to HEK293T cells at 40% confluency. Media was replaced in following day, and the cells were incubated for 72 hours. At the infection day, the incubated viral soup was collected from HEK293T cells. After a round of filtration using 0.45 μm filter, the viral soup was combined with equal volume of target cell line's media, and polybrene (Sigma-Adrich, TR-1003-G) was added to the combined media in 1:1000 dilution. The lentiviral mixture was then added to the target cells, and the media was replaced with fresh media in the following day. After 48 hours, infected cells were selected using puromycin.

**Synthetic Methods and Characterization**

- Starting materials and reagents were purchased from commercial suppliers and used without additional purification step.

- All solvents are reagent grade or HPLC grade.

- Nuclear Magnetic Resonance (NMR) spectra were recorded on Bruker Avance 400, NEO 500, 600 MHz spectrometer. Chemical shift are reported in parts per million (ppm, $\delta$) and calibrated against tetramethylsilane (TMS) or residual solvent signal. $^{1}H$ NMR: $CDCl_3$ (7.26) and DMSO-d6 (2.50) & $^{13}C$ NMR: $CDCl_3$ (77.0) and DMSO-d6 (39.5). Multiplicity is reported as follows: singlet (s), doublet (d), doublet of doublet (dd) doublet of triplet (dt), triplet (t), triplet of doublet (td), quartet (q), and multiplet (m). Coupling constants (J) are reported in Hertz (Hz)

- All reactions were monitored by Thin Layer Chromatography (TLC) from Merck KGaA (TLC Silica gel 60 $F_{254}$). Visualization of TLC was done using UV (254 nm and 365 nm) or chemical stain ($KMnO_4$).

- Purification of product was performed by flash column chromatography using a Biotage Isolera. Elution of compound was monitored with the equipped UV detector, and following Biotage Sfar columns were used based on the scale of synthesis: 5 g, 10 g, or 25 g. If additional purification is needed, compound was further purified using Thermo scientific's semi-prep reversed phase high-performance liquid chromatography equipped (RP-HPLC: Ultimate 3000 HPLC ) equipped with C18 column (Luna® 10 $\mu$m c18(2), 100 Å, Serial #:5293-0084). Elution of sample was monitored using DIONEX UltiMate 3000. Eluting buffer A: 100% distilled water + 0.1% trifluoracetic acid (TFA). Eluting buffer B: 95% acetonitrile + 5% distilled water + 0.1% TFA.

- High-resolution mass spectra (HRMS) were obtained using Q Exactive™ Plus Hybrid Quadrupole-Orbtrap™ Mass Spectrometer.

**General Procedure A:** A carboxylic acid (i.e. JQ1-COOH) was dissolved in dimethylformamide (DMF). Subsequently, Mono-boc diamine linker (1.2 Eq.), hexafluorophosphate Azabenzotriazole Tetramethyl Uronium (HATU, 2 Eq.), and DIEA: N,N-Diisopropylethylamine (DIEA, 5 Eq.) were added to the reaction mixture. The reaction was stirred at room temperature (23 °C) for 4 hours. The reaction was then diluted with water (20 mL), and the organic layer was extracted with EtOAc five times. Combined organic layer was washed with brine and dried over anhydrous $Na_2SO_4$ before being concentration *in vacuo*. The concentrated organic mixture was further purified through flash column chromatography (0-10% MeOH/DCM) to yield Boc-protected product.

**General Procedure B:** A piperazine (i.e. AR-targeting ligand) and DIEA (2 Eq.) were dissolved in DMF. After 10 minutes of stirring at room temperature, N-Boc protected alkyl bromide linker (1.2 Eq.) was added. The reaction mixture was stirred at room temperature for 12 hours. Th reaction was quencedh by diluting with water, and the organic layer was extracted with EtOAc three times. The combined organic layer was subjected to washing with brine and dried over $Na_2SO_4$. Subsequently, the organic layer was concentrated *in vacuo* and the mixture was purified with flash column chromatography (0-13% MeOH/DCM) to yield Boc-protected products.

**General procedure C:** A primary or secondary amine was dissolved in DMF along with DIEA (5 Eq.). After stirring at room temperature for 10 minutes, HATU (2 Eq.) and required butanoic acid (**3**, **4**, or **7**; 1.1. Eq.) was added. The reaction was stirred at room temperature for 4 hours and further diluted with water. Organic mixture was extracted using 5 rounds of 4:1 $CHCl_3$:IPA extraction. Combined organic layer was then dried with anhydrous $Na_2SO_4$. After removing solvent *in vacuo*, compound was purified using flash column chromatography (80-100% EtOAc/Hexane followed by 2-15% MeOH/DCM). Additional round of RP-HPLC purification was performed only if needed (40-80% eluting buffer B : eluting buffer A).

**tert-butyl 4-(3-hydroxybenzyl)piperidine-1-carboxylate (1)**

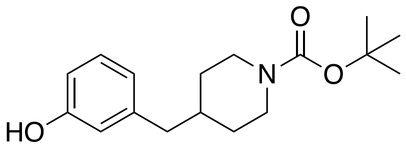

3-(piperidin-4-ylmethyl)phenol Hydrochloride (150 mg, 0.66 mmol) was dissolved in 5 mL of MeOH : $CH_2Cl_2$ (1 : 9) with DIEA (184 μL, 0.66 mmol). After 10 minutes of vigorous stirring at 23 °C, Di-tert-butyl decarbonate (314 mg, 0.86 mmol) was added to the stirring mixture, and the reaction was stirred for additional 12 hours. The reaction was quenched by removing the solvent under vacuum and the mixture was purified by flash column chromatography (hexane : EtOAc = 80 : 20 to 60 : 40). White crystalline solid **1** (130 mg, 68% yield) was obtained as an intermediate. $R_f$ = 0.6 (hexane : EtOAc = 50 : 50, UV). **$^1$H NMR** (600 MHz, $CDCl_3$) δ 7.13 (t, J = 7.8 Hz, 1H), 6.68 (dd, J = 7.9, 2.1 Hz, 2H), 6.64 (t, J = 2.0 Hz, 1H), 5.81 (d, J = 2.1 Hz, 1H), 4.06 (s, 2H), 2.64 (s, 2H), 2.47 (d, J = 7.0 Hz, 2H), 1.66 – 1.58 (m, 3H), 1.46 (s, 9H), 1.13 (tdd, J = 13.0, 11.1, 4.3 Hz, 2H). **LC/MS** $[M+Na]^+$ m/z calc: 314.2; found: 314.2

**4-(3-((1-(tert-butoxycarbonyl)piperidin-4-yl)methyl)phenoxy)butanoic acid (2)**

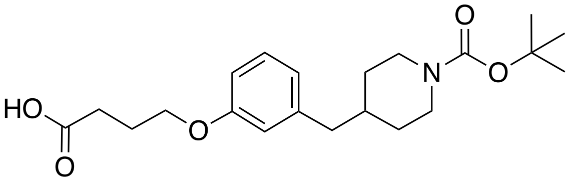

Intermediate **1** (100mg, 0.34 mmol) and $K_2CO_3$ (237 mg, 1.71 mmol) were dissolved in 5 mL DMF, and the reaction mixture was stirred at 55 °C for 30 minutes. Subsequently, ethyl 4-bromobutyrate (118 μL, 0.51 mmol) was added, and the reaction mixture was stirred for additional 6 hours at 55 °C with reflux setup. The solvent of the reaction was then removed under vacuum, and the reactants were re-dissolved in 50 mL of EtOAc. Dissolved reactant was washed with 10 mL of water twice, and the organic solvent was again removed *in vacuo*. Next, 10 mL of 2M NaOH : EtOH (5 : 5) was added to the resulting oil, and the reaction was vigorously stirred at 55 °C for 2 hours. After 2 hours, EtOH in the reaction mixture was removed as much as possible under vacuum, and the reaction containing flask was placed in an ice bath. 1M HCl was added dropwise to the reactant while stirring until the pH of the solution turn to 2 on a pH test strip. The precipitating compound was then extracted with EtOAc (3x). Collected organic layer was dried over $Na_2SO_4$ and further dried *in vacuo*. The intermediate was then purified using flash column chromatography (hexane : EtOAc = 60 : 40 to 20 : 80) to result white solid **2** (85 mg, 66 %) as a product. $R_f$ = 0.2 (hexane : EtOAc = 50:50, UV). **$^1$H NMR** (500 MHz, DMSO-d6) δ 12.13 (s, 1H), 7.16 (dd, J = 8.8, 7.0 Hz, 1H), 6.76 – 6.70 (m, 3H), 3.95 (t, J = 6.5 Hz, 2H), 3.88 (d, J = 11.6 Hz, 2H), 2.63 (s, 2H), 2.46 (d, J = 7.2 Hz, 2H), 2.38 (t, J = 7.3 Hz, 2H), 1.92 (p, J = 7.0 Hz, 2H), 1.65 (ddt, J = 11.4, 7.1, 3.7 Hz, 1H), 1.56 – 1.49 (m, 2H), 1.38 (s, 9H), 1.00 (qd, J = 12.4, 4.2 Hz, 2H). **13C NMR** (126 MHz, DMSO-d6) δ 174.1, 158.4, 153.8, 141.7, 129.1, 121.3, 115.1, 111.7, 78.4, 66.3, 54.9, 42.1, 37.2, 31.5, 30.1, 28.1, 24.3. **HRMS (ESI)** m/z: $[M+H]^+$ calc for $C_{21}H_{31}NO_5Na^+$ : 400.2100; found : 400.2059.

**4-(3-((1-acryloylpiperidin-4-yl)methyl)phenoxy)butanoic acid (3)**

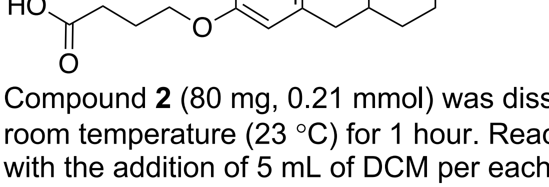

Compound **2** (80 mg, 0.21 mmol) was dissolved in 10 mL of DCM:TFA (50 : 50), and the reaction was stirred at room temperature (23 °C) for 1 hour. Reaction was then dried *in vacuo* and the drying step was repeated twice with the addition of 5 mL of DCM per each step. Next, 5 mL of THF was added to the dried reactant along with

DIEA (177 μL, 0.63 mmol). Reaction was stirred in the ice bath for 10 minute, and acryloyl chloride (40 μL, 0.31 mmol) was added. After 30 minutes of stirring with slowly increasing the reaction temperature to 23 °C, reaction mixture was concentrated and purified using flash column chromatography. Product **3** (33 mg, 47%) was obtained as clear oil after the flash column chromatography (Hexane : EtOAC = 80 : 20 to 0 : 100). $R_f$ = 0.5 (Hexane : EtOAc = 10 : 90, $KMnO_4$). **$^{1}$H NMR** (600 MHz, DMSO-d6) δ 12.12 (s, 1H), 7.20 – 7.14 (m, 1H), 6.80 – 6.75 (m, 1H), 6.75 – 6.71 (m, 3H), 6.06 (dd, J = 16.7, 2.4 Hz, 1H), 5.63 (dd, J = 10.5, 2.5 Hz, 1H), 4.38 (d, J = 13.2 Hz, 1H), 4.00 (d, J = 13.7 Hz, 1H), 3.95 (t, J = 6.4 Hz, 2H), 2.96 (t, J = 12.8 Hz, 1H), 2.60 – 2.52 (m, 1H), 2.47 (dd, J = 7.3, 2.5 Hz, 2H), 2.38 (t, J = 7.3 Hz, 2H), 1.92 (p, J = 6.9 Hz, 2H), 1.77 (ttt, J = 11.0, 7.3, 3.7 Hz, 1H), 1.60 (dd, J = 13.9, 3.6 Hz, 2H), 1.09 – 0.96 (m, 2H). **$^{13}$C NMR** (151 MHz, DMSO-d6) δ 174.1, 164.0, 158.4, 141.6, 129.1, 128.6, 126.8, 121.3, 115.1, 111.7, 66.3, 45.2, 42.0, 41.5, 37.3, 32.4, 31.4, 30.1, 24.3. **HRMS (ESI)** m/z: $[M+H]^+$ calc for $C_{19}H_{26}NO_4^+$ : 332.1862; found : 332.1846.

**4-(3-((1-propionylpiperidin-4-yl)methyl)phenoxy)butanoic acid (4)**

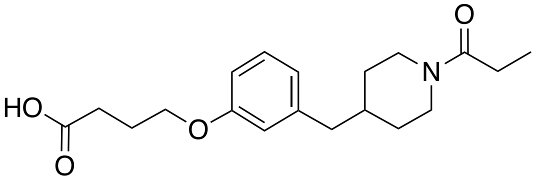


Compound **2** (80 mg, 0.21 mmol) was dissolved in 10 mL of DCM:TFA (50 : 50), and the reaction was stirred at the room temperature (23 °C) for 1 hour. Reaction was then dried *in vacuo* and the drying step was repeated twice with the addition of 5 mL of additional DCM. Next, 5 mL of THF was added to the dried reactant along with DIEA (177 μL, 0.63 mmol). Reaction was stirred in the ice bath for 10 minute, and propionyl chloride (40 μL, 0.31 mmol) was added. After 30 minutes of stirring at room temperature, reaction mixture was concentrated under vacuum and purified using flash column chromatography. Product **4** (17 mg, 24%) was obtained as a white solid after the flash column chromatography (Hexane : EtOAC = 80 : 20 to 0 : 100). $R_f$ = 0.4 (Hexane : EtOAc = 10 : 90, UV). **$^{1}$H NMR** (600 MHz, $CDCl_3$) δ 7.18 (t, *J* = 7.8 Hz, 1H), 6.75 – 6.69 (m, 2H), 6.67 (t, *J* = 2.0 Hz, 1H), 4.64 – 4.57 (m, 1H), 4.01 (t, *J* = 6.1 Hz, 2H), 3.81 (d, *J* = 13.3 Hz, 1H), 2.94 (td, *J* = 13.0, 2.6 Hz, 1H), 2.58 (t, *J* = 7.2 Hz, 2H), 2.56 – 2.50 (m, 2H), 2.50 – 2.44 (m, 2H), 2.34 (q, *J* = 7.5 Hz, 2H), 2.11 (p, *J* = 6.6 Hz, 2H), 1.76 (ddp, *J* = 11.2, 7.5, 3.6 Hz, 1H), 1.69 (dd, *J* = 13.4, 4.6 Hz, 2H), 1.18 – 1.09 (m, 5H). **$^{13}$C NMR** (151 MHz, $CDCl_3$) δ 177.7, 172.4, 158.8, 141.6, 129.2, 121.6, 115.5, 111.8, 66.5, 45.8, 43.0, 42.1, 38.14, 32.6, 31.8, 30.5, 26.6, 24.5, 9.7. **HRMS (ESI)** m/z: $[M+H]^+$ calc for $C_{19}H_{28}NO_4^+$ : 334.2018; found : 334.2011.

**tert-butyl 4-(4-hydroxybenzyl)piperidine-1-carboxylate (5)**

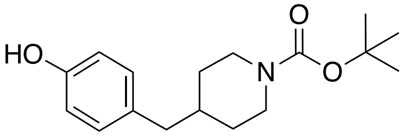


4-(piperidin-4-ylmethyl)phenol (150 mg, 0.66 mmol) was dissolved in 5 mL of MeOH : $CH_2Cl_2$ (5 : 95) with DIEA (184 uL, 0.66 mmol). After 10 minutes of vigorous stirring at 23 °C, Di-tert-butyl decarbonate (314 mg, 0.86 mmol) was added to the reaction mixture. The reaction was stirred at the same temperature (23 °C) for 14 hours. The solvent of the reactant was then removed *in vacuo* and the mixture was purified by column chromatography (hexane : EtOAc = 80 : 20 to 60 : 40) to afford white crystalline solid **5** in room temperature (133 mg, 69%) $R_f$ = 0.6 (hexane : EtOAc = 50 : 50, UV).**1H NMR** (600 MHz, CDCl3) δ 6.97 – 6.93 (m, 2H), 6.81 – 6.79 (m, 1H), 6.79 – 6.76 (m, 2H), 4.06 (s, 2H), 2.63 (s, 2H), 2.43 (d, J = 6.7 Hz, 2H), 1.67 – 1.53 (m, 3H), 1.46 (s, 9H), 1.17 – 1.05 (m, 2H). **LC/MS** $[M+Na]^+$ m/z calc: 314.2; found: 314.2

**4-(4-((1-(tert-butoxycarbonyl)piperidin-4-yl)methyl)phenoxy)butanoic acid (6)**

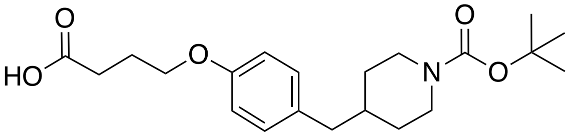

Intermediate **5** (100mg, 0.34 mmol) and $K_2CO_3$ (237 mg, 1.71 mmol) were dissolved in 5 mL DMF, and the reaction mixture was stirred at 55 °C for 30 minutes. Subsequently, ethyl 4-bromobutyrate (118 μL, 0.51 mmol) was added, and the reaction mixture was stirred for 6 hours at 55 °C with reflux setup. The solvent of the reaction was removed under vacuum, and the reactants were re-dissolved in 50 mL of EtOAc. Dissolved reactant was washed with 10 mL of water twice, and the EtOAc was again removed *in vacuo*. Next, 10 mL of 2M NaOH : EtOH (5 : 5) was added to the resulted oily intermediate, and the reaction was stirred at 55 °C vigorously for 2 hours. After 2 hours, EtOH in the reaction mixture was removed as much as possible *in vacuo*, and the reaction containing flask was placed in an ice bath. 1M HCl was added dropwise to the reactant with stirring until the pH of the solution turn to 2 on a pH test strip. The precipitating compound was then extracted with EtOAc (3x). Collected organic layer was dried over $Na_2SO_4$ and further concentrated *in vacuo*. The intermediate was then purified using flash column chromatography (hexane : EtOAc = 60 : 40 to 20 : 80) to result white solid **6** (85 mg, 66 %) $R_f$ = 0.2 (hexane : EtOAc = 50:50, UV). $^1$**H NMR** (400 MHz, $CDCl_3$) δ 7.03 (d, J = 8.5 Hz, 2H), 6.80 (d, J = 8.5 Hz, 2H), 4.23 – 3.90 (m, 4H), 2.78 – 2.52 (m, 4H), 2.46 (d, J = 6.7 Hz, 2H), 2.19 – 2.03 (m, 2H), 1.72 – 1.53 (m, 3H), 1.45 (s, 9H), 1.20 – 0.97 (m, 2H). $^{13}$**C NMR** (151 MHz, $CDCl_3$) δ 178.7, 157.1, 155.0, 132.5, 130.0, 114.3, 79.4, 66.5, 42.2, 38.4, 38.3, 31.9, 30.5, 28.5, 24.5. **HRMS (ESI)** m/z: $[M+H]^+$ calc for $C_{21}H_{31}NO_5Na^+$ : 400.2100; found : 400.2059.

**4-(4-((1-acryloylpiperidin-4-yl)methyl)phenoxy)butanoic acid (7)**

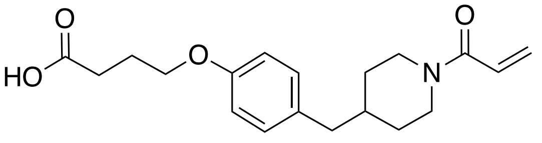

Compound **6** (80 mg, 0.21 mmol) was dissolved in 10 mL of DCM:TFA (50 : 50), and the reaction was stirred at the room temperature (23 °C) for 1 hour. Reaction was then dried *in vacuo* and the drying step was repeated twice with additional DCM. 5 mL of THF was then added to the dried reactant along with DIEA (177 μL, 0.63 mmol). Reacting vessel was then re-placed in an ice bath, and the reaction was stirred for 10 minute. Next, acryloyl chloride (40 μL, 0.31 mmol) was added and stirred for 30 minutes. Solvent was removed *in vacuo* and resulting oil was purified using flash column chromatography. Product **7** (33 mg, 47%) was obtained as clear oil. (Hexane : EtOAC = 50 : 50 to 0 : 100). $R_f$ = 0.4 (Hexane : EtOAc = 10 : 90, UV). $^1$**H NMR** (400 MHz, DMSO-d6) δ 7.13 – 6.98 (m, 2H), 6.93 – 6.71 (m, 3H), 6.06 (dd, J = 16.7, 2.5 Hz, 1H), 5.63 (dd, J = 10.5, 2.5 Hz, 1H), 4.38 (d, J = 13.0 Hz, 1H), 4.00 (d, J = 13.8 Hz, 1H), 3.94 (t, J = 6.4 Hz, 2H), 2.95 (t, J = 12.6 Hz, 1H), 2.58 – 2.51 (m, 1H), 2.44 (d, J = 7.1 Hz, 2H), 2.37 (t, J = 7.3 Hz, 2H), 1.92 (p, J = 6.9 Hz, 2H), 1.78 – 1.65 (m, 1H), 1.65 – 1.54 (m, 2H), 1.17 – 0.86 (m, 2H). $^{13}$**C NMR** (151 MHz, DMSO-d6) δ 174.6, 164.5, 157.2, 132.3, 130.4, 129.1, 127.2, 114.6, 66.9, 45.7, 42.1, 41.6, 38.0, 32.8, 31.8, 30.6, 24.8. **HRMS (ESI)** m/z: $[M+H]^+$ calc for $C_{19}H_{26}NO_4^+$ : 332.1862; found : 332.1846.

**N-(5-(4-(3-((1-acryloylpiperidin-4-yl)methyl)phenoxy)butanamido)pentyl)-4-ethynylbenzamide (SJH1-37m)**

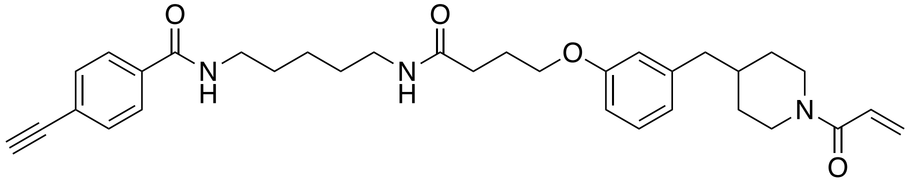

N-(5-aminopentyl)-4-ethynylbenzamide was synthesized based on previously reported synthetic route.[1] **3** (30 mg, 0.09 mmol) was dissolved in 2 mL of DMF. To a stirring mixture, N-(5-aminopentyl)-4-ethynylbenzamide (31 mg, 0.13 mmol), HATU (69 mg, 0.18 mmol), and DIEA (63 μL, 0.36 mmol) were subsequently added. Reaction proceeded at 23 °C for 4 hours, and the reaction was quenched by adding 5 mL of water. Reactants were extracted with EtOAc five times, and the organic layer was dried over $Na_2SO_4$ and concentrated under vacuum. The resulting oil was purified using flash column chromatography (DCM : MeOH = 100 : 0 to 95 : 5) over two step, and white powder was acquired as a product (19 mg, 39%). $R_f$ = 0.8 (DCM : MeOH = 90:10, UV) **$^1$H NMR** (600 MHz, DMSO-d6) δ 8.50 (t, J = 5.6 Hz, 1H), 7.85 – 7.82 (m, 2H), 7.80 (t, J = 5.6 Hz, 1H), 7.57 – 7.53 (m, 2H), 7.17 (t, J = 7.7 Hz, 1H), 6.80 – 6.69 (m, 4H), 6.05 (dd, J = 16.7, 2.4 Hz, 1H), 5.63 (dd, J = 10.5, 2.4 Hz, 1H), 4.37 (d, J = 12.9 Hz, 1H), 4.34 (s, 1H), 3.99 (d, J = 13.8 Hz, 1H), 3.91 (t, J = 6.4 Hz, 2H), 3.23 (q, J = 6.7 Hz, 2H), 3.04 (q, J = 6.6 Hz, 2H), 2.96 (t, J = 12.8 Hz, 1H), 2.56 (t, J = 12.4 Hz, 1H), 2.47 (dd, J = 7.2, 1.8 Hz, 2H), 2.21 (t, J = 7.4 Hz, 2H), 1.90 (p, J = 6.8 Hz, 2H), 1.81 – 1.72 (m, 1H), 1.62 – 1.56 (m, 2H), 1.51 (p, J = 7.3 Hz, 2H), 1.41 (p, J = 7.2 Hz, 2H), 1.33 – 1.24 (m, 2H), 1.03 (h, J = 11.5 Hz, 2H). **$^{13}$C NMR** (151 MHz, DMSO-d6) δ 171.3, 165.2, 164.0, 158.5, 141.6, 134.7, 131.5, 129.1, 128.6, 127.4, 126.8, 124.2, 121.2, 115.2, 111.7, 82.9, 82.6, 66.7, 45.1, 42.0, 41.5, 38.3, 37.2, 32.4, 31.7, 31.4, 28.8, 28.7, 25.0, 23.8. **HRMS (ESI)** m/z: $[M+H]^+$ calc for $C_{33}H_{42}N_3O_4^+$: 544.3164; found: 544.3119

**N-(5-(4-(4-((1-acryloylpiperidin-4-yl)methyl)phenoxy)butanamido)pentyl)-4-ethynylbenzamide (SJH1-37p)**

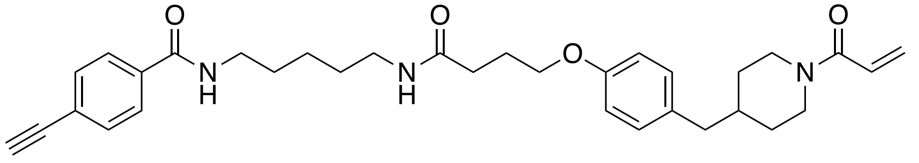

**7** (30 mg, 0.09 mmol) was dissolved in 2 mL of DMF. Next, N-(5-aminopentyl)-4-ethynylbenzamide (31 mg, 0.13 mmol), HATU (69 mg, 0.18 mmol), and DIEA (63 μL, 0.36 mmol) were subsequently added to the stirring reaction mixture. Reaction proceeded at 23 °C for 4 hours, and the reaction was diluted in water. Reactants were extracted with EtOAc five times, and the organic layer was dried over $Na_2SO_4$ and concentrated *in vacuo*. Resulting oil was purified using flash column chromatography (DCM : MeOH = 100 : 0 to 95 : 5), and white powder was acquired as a product (20mg, 41%). $R_f$ = 0.8 (DCM : MeOH = 90:10, UV) **1H NMR** (400 MHz, DMSO-d6) δ 8.52 (t, J = 5.6 Hz, 1H), 7.93 – 7.76 (m, 3H), 7.56 (d, J = 8.1 Hz, 2H), 7.05 (d, J = 8.2 Hz, 2H), 6.90 – 6.67 (m, 3H), 6.06 (dd, J = 16.7, 2.5 Hz, 1H), 5.63 (dd, J = 10.4, 2.5 Hz, 1H), 4.48 – 4.29 (m, 2H), 3.99 (d, J = 13.7 Hz, 1H), 3.89 (t, J = 6.4 Hz, 2H), 3.23 (q, J = 6.6 Hz, 2H), 3.04 (q, J = 6.6 Hz, 2H), 2.94 (t, J = 12.9 Hz, 1H), 2.63 – 2.52 (m, 1H), 2.43 (d, J = 7.1 Hz, 2H), 2.21 (t, J = 7.4 Hz, 2H), 1.90 (p, J = 6.8 Hz, 2H), 1.69 (ddt, J = 12.8, 9.2, 4.6 Hz, 1H), 1.65 – 1.54 (m, 2H), 1.54 – 1.44 (m, 2H), 1.44 – 1.35 (m, 2H), 1.34 – 1.22 (m, 2H), 1.06 – 0.91 (m, 2H). **$^{13}$C NMR** (151 MHz, DMSO-d6) δ 171.3, 165.2, 164.0, 156.7, 134.7, 131.7, 131.5, 129.8, 128.5, 127.3, 126.7, 124.1, 114.0, 82.8, 82.6, 79.1, 66.8, 45.1, 41.5, 41.0, 38.3, 37.5, 32.3, 31.7, 31.2, 28.8, 28.6, 24.9, 23.8. **HRMS (ESI)** m/z: $[M+H]^+$ calc for $C_{33}H_{42}N_3O_4^+$: 544.3164; found: 544.3119.

**tert-butyl (S)-(2-(2-(4-(4-chlorophenyl)-2,3,9-trimethyl-6H-thieno[3,2-f][1,2,4]triazolo[4,3-a][1,4]diazepin-6-yl)acetamido)ethyl)carbamate (8)**

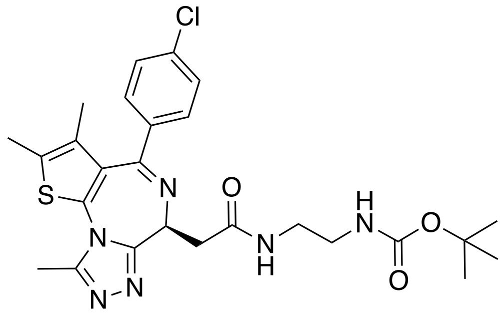

JQ-1 (50 mg, 0.12 mmol) was reacted with tert-butyl (2-aminoethyl)carbamate (20 mg, 0.12 mmol), HATU (95 mg, 0.25 mmol), and DIEA (109 μL, 0.62 mmol) in 2 mL DMF. Reaction was carried out based on the general procedure **A**. After flash column chromatography, intermediate **8** was obtained as an oil. (38 mg, 56%). $R_f$ =

0.4 (DCM : MeOH = 95 : 5, UV). $^{1}$**H NMR** (600 MHz, $CDCl_3$) δ 7.44 (s, 1H), 7.37 (d, J = 8.4 Hz, 2H), 7.30 (d, J = 8.6 Hz, 2H), 5.47 (s, 1H), 3.53 (dd, J = 14.7, 7.6 Hz, 1H), 3.40 (td, J = 14.2, 6.4 Hz, 3H), 3.35 – 3.18 (m, 3H), 2.67 (s, 3H), 2.39 (s, 3H), 1.66 (s, 3H), 1.39 (s, 9H). **HRMS** (ESI) m/z: $[M+H]^+$ calc for $C_{26}H_{32}ClN_6O_3S^+$: 543.1945; found: 543.1946.

**tert-butyl (S)-(4-(2-(4-(4-chlorophenyl)-2,3,9-trimethyl-6H-thieno[3,2-f][1,2,4]triazolo[4,3-a][1,4]diazepin-6-yl)acetamido)butyl)carbamate (9)**

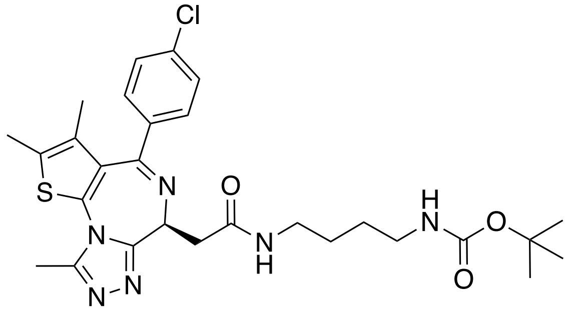


JQ-1 (50 mg, 0.12 mmol) was dissolved in 2 mL of DMF, and tert-butyl (4-aminobutyl)carbamate (24 mg, 0.12 mmol), HATU (95 mg, 0.25 mmol) and DIEA (109 μL, 0.62 mmol) were added to reactant. Reaction was carried out based on the general procedure **A**. After purification, intermediate **9** was obtained as an oil. (41 mg, 58%). $R_f$ = 0.4 (DCM : MeOH = 95 : 5, UV). $^{1}$**H NMR** (400 MHz, DMSO-d6) δ 8.19 (s, 1H), 7.50 (d, J = 8.1 Hz, 2H), 7.42 (d, J = 8.2 Hz, 2H), 6.81 (s, 1H), 4.53 – 4.46 (m, 1H), 3.24 – 3.01 (m, 3H), 2.99 – 2.81 (m, 3H), 2.60 (s, 3H), 2.41 (s, 3H), 1.62 (s, 3H), 1.46 – 1.34 (m, 13H). **HRMS** (ESI) m/z: $[M+H]^+$ calc for $C_{28}H_{36}ClN_6O_3S^+$: 571.2258; found: 571.2199.

**tert-butyl (S)-(5-(2-(4-(4-chlorophenyl)-2,3,9-trimethyl-6H-thieno[3,2-f][1,2,4]triazolo[4,3-a][1,4]diazepin-6-yl)acetamido)pentyl)carbamate (10)**

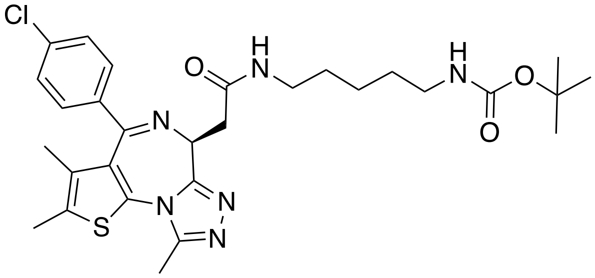


Following reaction was carried out based on general procedure **A**. JQ-1 (50 mg, 0.12 mmol) was reacted with tert-butyl (5-aminopentyl)carbamate (25 mg, 0.12 mmol) in the presence of HATU (95 mg, 0.25 mmol), and DIEA (109 μL, 0.62 mmol) in 2 mL DMF. **10** was obtained as an oil after the flash column chromatography. (61 mg, 83%). $R_f$ = 0.5 (DCM : MeOH = 95 : 5, UV). $^{1}$**H NMR (**500 MHz, DMSO-d6) δ 8.16 (t, J = 5.7 Hz, 1H), 7.49 (d, J = 8.5 Hz, 2H), 7.45 – 7.38 (m, 2H), 6.78 (t, J = 5.7 Hz, 1H), 4.50 (dd, J = 8.3, 5.9 Hz, 1H), 3.25 (dd, J = 15.0, 8.3 Hz, 1H), 3.21 – 3.09 (m, 3H), 3.05 (dt, J = 13.1, 6.4 Hz, 1H), 2.93 – 2.86 (m, 3H), 2.59 (s, 3H), 2.41 (s, 3H), 1.62 (s, 3H), 1.51 – 1.33 (m, 13H). **HRMS** (ESI) m/z: $[M+H]^+$ calc for $C_{29}H_{38}ClN_6O_3S^+$: 585.2414; found: 585.2412

**tert-butyl (S)-(1-(4-(4-chlorophenyl)-2,3,9-trimethyl-6H-thieno[3,2-f][1,2,4]triazolo[4,3-a][1,4]diazepin-6-yl)-2-oxo-6,9,12-trioxa-3-azatetradecan-14-yl)carbamate (11)**

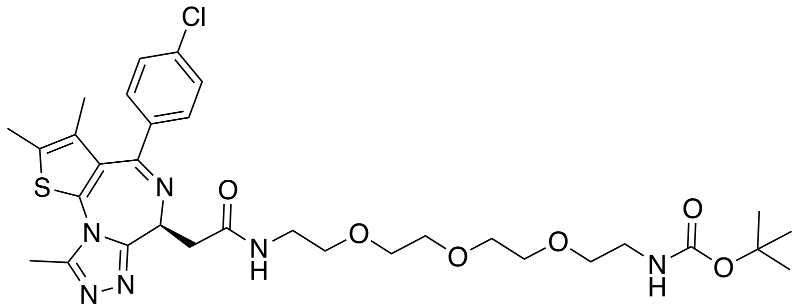

JQ-1 (50 mg, 0.12 mmol) was dissolved in 2 mL of DMF along with tert-butyl (2-(2-(2-(2-aminoethoxy)ethoxy)ethoxy)ethyl)carbamate (31 mg, 0.12 mmol), HATU (95 mg, 0.25 mmol), and DIEA (109 μL, 0.62 mmol). After 12 hours of stirring at 23 °C, the reaction was diluted with water. Reactants were then extracted using 5 rounds of 4:1 $CHCl_3$:IPA. Organic layer was dried over $Na_2SO_4$ and further concentrated *in vacuo*. Resulting oil was then purified using flash column chromatography (DCM : MeOH = 100 : 0 to 85 : 11) to afford **11** as a oil. (29 mg, 34%). $R_f$ = 0.4 (DCM : MeOH = 95 : 5, UV). **$^1$H NMR** (600 MHz, $CDCl_3$) δ 7.40 – 7.35 (m, 2H), 7.33 – 7.28 (m, 2H), 7.05 (s, 1H), 5.23 (s, 1H), 3.69 – 3.45 (m, 16H), 3.37 (dd, J = 14.6, 7.0 Hz, 1H), 3.29 (d, J = 6.1 Hz, 2H), 2.65 (s, 3H), 2.38 (s, 3H), 1.65 (s, 3H), 1.40 (s, 9H). **HRMS** (ESI) m/z: $[M+H]^+$ calc for $C_{32}H_{44}ClN_6O_6S^+$: 675.2731; found: 675.2660

**(S)-4-(3-((1-acryloylpiperidin-4-yl)methyl)phenoxy)-N-(2-(2-(4-(4-chlorophenyl)-2,3,9-trimethyl-6H-thieno[3,2-f][1,2,4]triazolo[4,3-a][1,4]diazepin-6-yl)acetamido)ethyl)butanamide (SJH1-51A)**

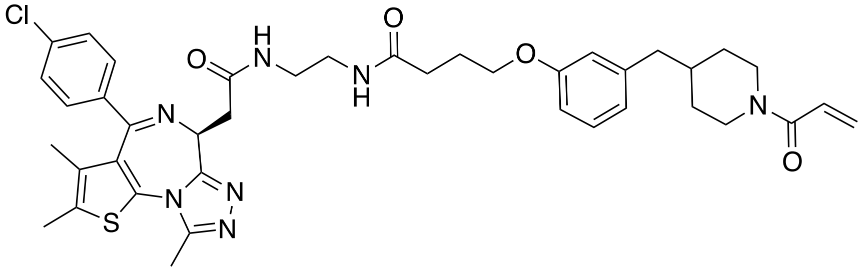

**8** (30 mg, 0.05 mmol) was dissolved in 10 mL of DCM:TFA (50 : 50), and the reaction was stirred at 23 °C for 1 hour. Reaction was then dried *in vacuo* and the drying step was repeated twice with the addition of 5 mL of additional DCM. Resulting oil was then dissolved in 1.5 mL of DMF, and HATU (42 mg, 0.11 mmol), DIEA (48 μL, 0.27 mmol), and **3** (18 mg, 0.06 mmol) were added to the stirring mixture. Following reaction was carried based on general procedure **C**. Obtained yellow oil was then purified using flash column chromatography (DCM : MeOH = 100 : 0 to 90 : 10) over two step, and the final product **SJH1-51A** was obtained as a white powder (11 mg, 26%). $R_f$ = 0.7 (DCM : MeOH = 90 : 10, $KMnO_4$). **$^1$H NMR** (600 MHz, DMSO-d6) δ 8.23 (t, J = 5.7 Hz, 1H), 7.86 (t, J = 5.1 Hz, 1H), 7.50 – 7.45 (m, 2H), 7.45 – 7.40 (m, 2H), 7.15 (t, J = 8.0 Hz, 1H), 6.80 – 6.69 (m, 4H), 6.05 (dd, J = 16.7, 2.4 Hz, 1H), 5.62 (dd, J = 10.5, 2.4 Hz, 1H), 4.51 (t, J = 6.6 Hz, 1H), 4.37 (d, J = 13.1 Hz, 1H), 3.99 (d, J = 13.5 Hz, 1H), 3.92 (t, J = 6.5 Hz, 2H), 3.23 (dd, J = 7.2, 2.7 Hz, 2H), 3.20 – 3.12 (m, 4H), 2.95 (t, J = 12.9 Hz, 1H), 2.63 – 2.51 (m, 4H), 2.46 (dd, J = 7.2, 2.7 Hz, 2H), 2.40 (s, 3H), 2.27 – 2.19 (m, 2H), 1.92 (p, J = 6.9 Hz, 2H), 1.76 (ddh, J = 11.0, 7.2, 3.5 Hz, 1H), 1.63 – 1.56 (m, 5H), 1.02 (t, 2H). **$^{13}$C NMR** (151 MHz, DMSO-d6) δ 171.6, 169.6, 164.0, 163.0, 158.4, 158.1, 157.8, 155.0, 149.8, 141.5, 136.6, 135.1, 132.1, 130.6, 130.1, 129.8, 129.5, 129.0, 128.5, 128.4, 126.7, 121.1, 115.1, 111.6, 66.6, 53.7, 45.1, 41.9, 41.5, 38.4, 38.3, 37.5, 37.2, 32.3, 31.7, 31.3, 24.8, 13.9, 12.6, 11.2. **HRMS** (ESI) m/z: $[M+H]^+$ calc for $C_{40}H_{47}ClN_7O_4S^+$ : 756.3099; found: 756.3095

**(S)-4-(3-((1-acryloylpiperidin-4-yl)methyl)phenoxy)-N-(4-(2-(4-(4-chlorophenyl)-2,3,9-trimethyl-6H-thieno[3,2-f][1,2,4]triazolo[4,3-a][1,4]diazepin-6-yl)acetamido)butyl)butanamide (SJH1-51B)**

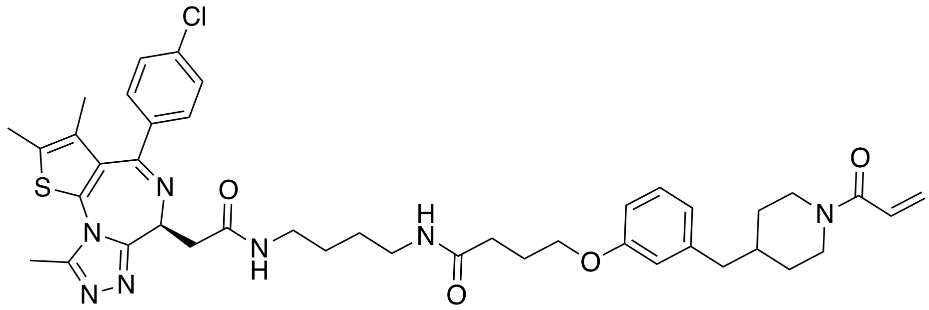

**9** (30 mg, 0.05 mmol) was dissolved in 10 mL of DCM:TFA (50 : 50), and the reaction was stirred at 23 °C for 1 hour. Reaction was then dried under vacuum and the drying step was repeated twice with the addition of 5 mL DCM. Next, resulting oil was dissolved in 1 mL of DMF, and HATU (40 mg, 0.11 mmol), DIEA (46 μL, 0.26 mmol), and **3** (17 mg, 0.05 mmol) were added to the stirring mixture. Following reaction was carried by following the general procedure **C**. Obtained yellow oil was then purified using flash column chromatography (DCM : MeOH = 100 : 0 to 90 : 10) over two step, and the final product **SJH1-51B** was obtained as a white powder (11 mg, 27%). $R_f$ = 0.7 (DCM : MeOH = 90 : 10, UV). **$^1$H NMR** (600 MHz, DMSO-d6) δ 8.16 (t, *J* = 5.7 Hz, 1H), 7.81 (t, *J* = 5.6 Hz, 1H), 7.51 – 7.47 (m, 2H), 7.44 – 7.39 (m, 2H), 7.16 (t, *J* = 7.9 Hz, 1H), 6.79 – 6.69 (m, 4H), 6.05 (dd, *J* = 16.7, 2.4 Hz, 1H), 5.62 (dd, *J* = 10.5, 2.4 Hz, 1H), 4.51 (dd, *J* = 8.3, 5.9 Hz, 2H), 4.37 (d, *J* = 13.1 Hz, 1H), 3.99 (d, *J* = 13.7 Hz, 1H), 3.92 (t, *J* = 6.4 Hz, 2H), 3.25 (dd, *J* = 15.0, 8.3 Hz, 1H), 3.21 – 3.10 (m, 3H), 3.09 – 3.03 (m, 3H), 2.96 (t, *J* = 12.9 Hz, 1H), 2.59 (s, 4H), 2.48 – 2.44 (m, 2H), 2.41 (s, 3H), 2.22 (t, *J* = 7.4 Hz, 2H), 1.91 (p, *J* = 6.9 Hz, 2H), 1.77 (ddq, *J* = 11.3, 7.6, 3.8 Hz, 1H), 1.65 – 1.54 (m, 5H), 1.47 – 1.39 (m, 4H), 1.06 – 0.97 (m, 2H). **$^{13}$C NMR** (151 MHz, DMSO-d6) δ 171.3, 169.2, 164.0, 162.9, 158.4, 158.1, 157.9, 155.0, 149.8, 141.5, 136.6, 135.1, 132.1, 130.6, 130.0, 129.7, 129.5, 129.0, 128.5, 128.4, 126.7, 121.1, 115.1, 111.6, 66.7, 53.8, 45.1, 41.9, 41.5, 38.2, 38.1, 37.5, 37.2, 32.3, 31.7, 31.3, 26.6, 26.5, 24.9, 13.9, 12.6, 11.2. **HRMS** (ESI) m/z: $[M+H]^+$ calc for $C_{42}H_{51}ClN_7O_4S^+$ : 784.3412; found: 784.3395

**(S)-N-(4-(2-(4-(4-chlorophenyl)-2,3,9-trimethyl-6H-thieno[3,2-f][1,2,4]triazolo[4,3-a][1,4]diazepin-6-yl)acetamido)butyl)-4-(3-((1-propionylpiperidin-4-yl)methyl)phenoxy)butanamide (SJH1-51B-NC)**

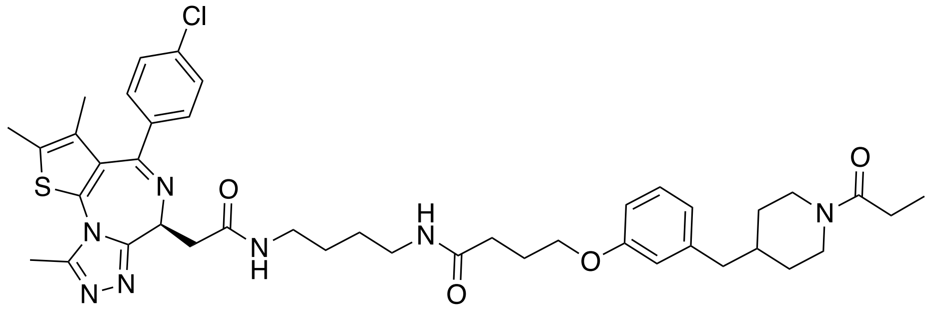

**9** (30 mg, 0.05 mmol) was dissolved in 10 mL of DCM:TFA (50 : 50), and the reaction was stirred at the room temperature (23 °C) for 1 hour. Reaction was then dried *in vacuo* and the drying step was repeated twice with the addition of 5 mL of additional DCM. Resulting oil was then dissolved in 1 mL of DMF, and HATU (40 mg, 0.11 mmol), DIEA (46 μL, 0.26 mmol), and **4** (17 mg, 0.05 mmol) were added to the stirring mixture. Following reaction was carried based on general procedure **C**. Obtained yellow oil was then purified using flash column chromatography (DCM : MeOH = 100 : 0 to 90 : 10) and another round of purification was performed using HPLC (buffer B : buffer A = 30:70 to 80:20). Fractions were collected and freeze-dried. Final product **SJH1-51B-NC** was obtained as a white powder (5.5 mg, 14%). $R_f$ = 0.5 (DCM : MeOH = 90 : 10, UV). **$^1$H NMR** (600 MHz, DMSO-d6) δ 8.16 (t, *J* = 5.7 Hz, 1H), 7.81 (t, *J* = 5.6 Hz, 1H), 7.51 – 7.46 (m, 2H), 7.44 – 7.39 (m, 2H), 7.16 (t, *J* = 7.9 Hz, 1H), 6.74 – 6.69 (m, 3H), 4.50 (dd, *J* = 8.3, 5.9 Hz, 1H), 4.34 (d, *J* = 13.1 Hz, 1H), 3.92 (t, *J* = 6.4 Hz, 2H), 3.79 (d, *J* = 13.6 Hz, 1H), 3.27 – 3.22 (m, 1H), 3.20 – 3.10 (m, 3H), 3.09 – 3.03 (m, 3H), 2.89 (t, *J* = 12.7 Hz, 1H), 2.59 (s, 3H), 2.48 – 2.42 (m, 3H), 2.41 (s, 3H), 2.29 – 2.25 (m, 2H), 2.22 (t, *J* = 7.4 Hz, 2H), 1.91 (p, *J* = 6.8 Hz, 2H), 1.76 – 1.69 (m, 1H), 1.62 (s, 3H), 1.56 (t, *J* = 14.4 Hz, 2H), 1.43 (p, *J* = 3.2 Hz, 4H), 1.10 – 1.01 (m, 2H), 0.96 (t, *J* = 7.4 Hz, 3H). **$^{13}$C NMR** (151 MHz, DMSO-d6) δ 171.3, 170.8, 169.3, 162.9, 158.4, 155.0, 149.7, 141.5, 136.7, 135.1, 132.1, 130.6, 130.0, 129.7, 129.5, 129.0, 128.4, 121.1, 117.6, 115.1,

111.6, 107.4, 66.6, 53.8, 44.8, 42.0, 41.0, 38.1, 38.1, 37.5, 37.2, 32.1, 31.7, 31.3, 26.6, 26.5, 25.5, 24.9, 13.9, 12.6, 11.2, 9.4. **HRMS** (ESI) m/z: $[M+H]^+$ calc for $C_{42}H_{53}ClN_7O_4S^+$ : 786.3568; found: 786.3546

**(S)-4-(3-((1-acryloylpiperidin-4-yl)methyl)phenoxy)-N-(5-(2-(4-(4-chlorophenyl)-2,3,9-trimethyl-6H-thieno[3,2-f][1,2,4]triazolo[4,3-a][1,4]diazepin-6-yl)acetamido)pentyl)butanamide (SJH1-51C)**

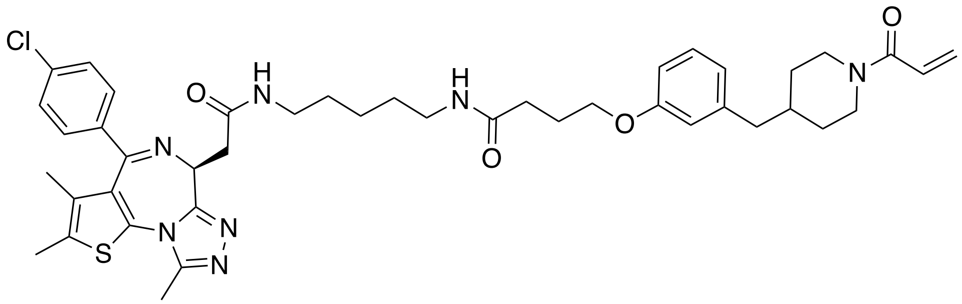

**10** (30 mg, 0.05 mmol) was dissolved in 10 mL of DCM:TFA (50 : 50), and the reaction was stirred at 23 °C for 1 hour. Reaction was then dried *in vacuo* and the drying step was repeated twice with the addition of 5 mL of additional DCM. Resulting oil was then dissolved in 1 mL of DMF, and HATU (39 mg, 0.10 mmol), DIEA (45 μL, 0.20 mmol), and **3** (17 mg, 0.05 mmol) were added to the stirring mixture. Following reaction was carried based on general procedure **C**. Obtained yellow oil was then purified using flash column chromatography (DCM : MeOH = 100 : 0 to 90 : 10) over two step to yield the final product **SJH1-51C** as a white powder (8 mg, 20%). $R_f$ = 0.7 (DCM : MeOH = 90 : 10, $KMnO_4$). **$^1$H NMR** (600 MHz, DMSO-d6) δ 8.16 (t, *J* = 5.7 Hz, 1H), 7.81 (t, *J* = 5.5 Hz, 1H), 7.51 – 7.46 (m, 2H), 7.44 – 7.39 (m, 2H), 7.16 (t, *J* = 8.0 Hz, 1H), 6.80 – 6.69 (m, 4H), 6.05 (dd, *J* = 16.7, 2.5 Hz, 1H), 5.62 (dd, *J* = 10.5, 2.4 Hz, 1H), 4.51 (dd, *J* = 8.2, 6.0 Hz, 1H), 4.37 (d, *J* = 13.1 Hz, 1H), 3.99 (d, *J* = 13.6 Hz, 1H), 3.92 (t, *J* = 6.4 Hz, 2H), 3.25 (dd, *J* = 15.0, 8.2 Hz, 1H), 3.21 – 3.08 (m, 3H), 3.08 – 3.00 (m, 3H), 2.95 (t, *J* = 12.8 Hz, 1H), 2.59 (s, 3H), 2.54 (m , 1H), 2.46 (dd, *J* = 7.1, 2.1 Hz, 2H), 2.42 – 2.39 (m, 3H), 2.22 (t, *J* = 7.4 Hz, 2H), 1.95 – 1.88 (m, 2H), 1.76 (dtp, *J* = 14.8, 7.2, 3.6 Hz, 1H), 1.63 – 1.56 (m, 5H), 1.42 (dq, *J* = 18.7, 7.3 Hz, 4H), 1.29 (td, *J* = 8.3, 4.2 Hz, 2H), 1.02 (t, *J* = 12.8 Hz, 2H). **$^{13}$C NMR** (151 MHz, DMSO-d6) δ 171.2, 169.2, 164.0, 162.9, 158.4, 158.0, 157.8, 155.0, 149.7, 141.5, 136.6, 135.1, 132.1, 130.6, 130.0, 129.7, 129.5, 129.0, 128.5, 128.4, 126.7, 121.1, 115.1, 111.6, 66.7, 53.8, 45.1, 41.9, 41.5, 38.3, 37.5, 37.2, 32.3, 31.6, 31.3, 28.8, 28.7, 24.9, 23.7, 13.9, 12.6, 11.2. **HRMS** (ESI) m/z: $[M+H]^+$ calc for $C_{43}H_{53}ClN_7O_4S^+$ : 797.3568; found: 797.3557

**(*S*)-4-(3-((1-acryloylpiperidin-4-yl)methyl)phenoxy)-*N*-(1-(4-(4-chlorophenyl)-2,3,9-trimethyl-6*H*-thieno[3,2-*f*][1,2,4]triazolo[4,3-*a*][1,4]diazepin-6-yl)-2-oxo-6,9,12-trioxa-3-azatetradecan-14-yl)butanamide (SJH1-51D)**

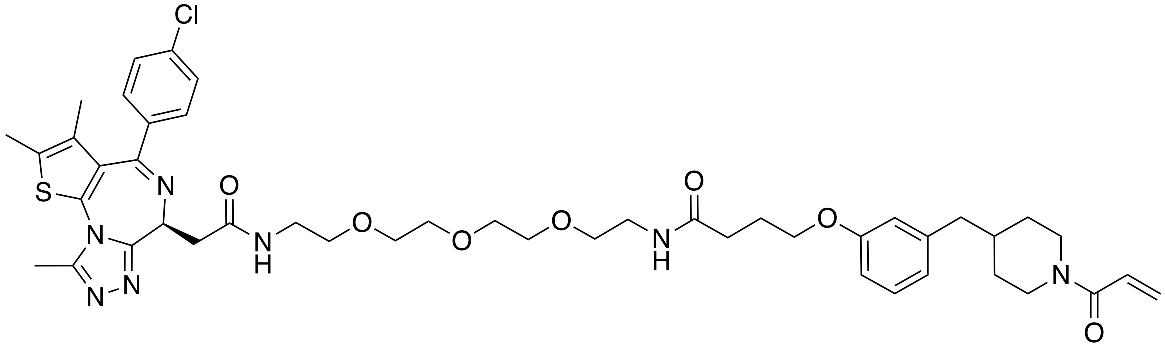

**11** (30 mg, 0.04 mmol) was dissolved in 10 mL of DCM:TFA (50 : 50), and the reaction was stirred at 23 °C for 1 hour. Reaction was then dried *in vacuo* and the drying step was repeated twice with the addition of 5 mL of additional DCM. Resulting oil was then dissolved in 1 mL of DMF, and HATU (39 mg, 0.10 mmol), DIEA (38 μL, 0.20 mmol), and **3** (15 mg, 0.04 mmol) were added to the stirring mixture. Following reaction was carried based on general procedure **C**. Obtained yellow oil was then purified using flash column chromatography (DCM : MeOH = 100 : 0 to 85 : 15) and further purified using RP-HPLC (Buffer A : Buffer B = 70 : 30 to 30 to 70). Collected fractions were freeze dried and the final product **SJH1-51D** was obtained as a white powder (6 mg, 15%). **$^1$H NMR** (600 MHz, DMSO-d6) δ 8.27 (t, *J* = 5.7 Hz, 1H), 7.90 (t, *J* = 5.6 Hz, 1H), 7.48 (d, *J* = 8.6 Hz, 2H), 7.45 – 7.40 (m, 2H), 7.20 – 7.13 (m, 1H), 6.80 – 6.69 (m, 4H), 6.05 (dd, *J* = 16.7, 2.4 Hz, 1H), 5.62 (dd, *J* = 10.5, 2.4 Hz, 1H), 4.52 (dd, *J* = 8.1, 6.0 Hz, 1H), 4.37 (d, *J* = 13.1 Hz, 1H), 4.00 (d, *J* = 13.6 Hz, 1H), 3.92 (t, *J* = 6.4 Hz, 2H), 3.56 – 3.48 (m, 8H), 3.46 (t, *J* = 5.9 Hz, 2H), 3.40 (t, *J* = 5.9 Hz, 2H), 3.34 – 3.24 (m, 3H), 3.23 – 3.18 (m, 3H), 2.95 (t, *J* = 13.0 Hz, 1H), 2.61 – 2.52 (m, 4H), 2.46 (dd, *J* = 7.2, 2.0 Hz, 2H), 2.41 (s, 3H), 2.24 (t, *J* = 7.4 Hz, 2H), 1.95 – 1.85 (m, 2H), 1.76 (dqt, *J* = 11.2, 7.5, 3.8 Hz, 1H), 1.64 – 1.56 (m, 5H), 1.02 (t, *J* = 13.1 Hz, 2H). **$^{13}$C NMR** (151 MHz, DMSO-d6) δ 170.5, 168.5, 162.9, 161.9, 157.4, 157.1, 156.9, 153.9, 148.7, 140.5, 135.6, 134.1, 131.1, 129.6, 129.0, 128.7, 128.4, 128.0, 127.5, 127.3, 125.6, 120.1, 114.0, 110.6, 68.6, 68.6, 68.5, 68.4, 68.1, 68.0, 65.6, 52.7, 44.0, 40.9, 40.4, 37.5, 37.4, 36.3, 36.1, 31.3, 30.5, 30.3, 23.8, 12.9, 11.5, 10.1. **HRMS** (ESI) m/z: $[M+H]^+$ calc for $C_{46}H_{59}ClN_7O7_S^+$ : 888.3885; found: 888.3875

**tert-butyl (4-(4-(6-((4-(3-chloro-4-cyanophenoxy)cyclohexyl)carbamoyl)pyridazin-3-yl)piperazin-1-yl)butyl)carbamate (12)**

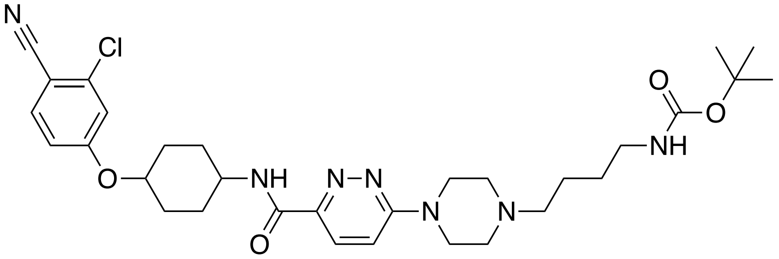

AR targeting ligand, N-(4-(3-chloro-4-cyanophenoxy)cyclohexyl)-6-(piperazin-1-yl)pyridazine-3-carboxamide, was synthesized based on previously reported synthetic route.[2] Reaction was carried based on the general procedure **B**, and following amount of each reagent was used: N-(4-(3-chloro-4-cyanophenoxy)cyclohexyl)-6-(piperazin-1-yl)pyridazine-3-carboxamide (50 mg, 0.11 mmol), DIEA (40 μL, 0.23 mmol), and tert-butyl (4-bromobutyl)carbamate was added (35 mg, 0.14 mmol). Intermediate **12** (35 mg, 50%) was obtained after the purification using flash column chromatagraphy (DCM : MeOH = 100 : 0 to 90 : 10). $R_f$ = 0.7 (DCM : MeOH = 90 : 10, UV). **$^1$H NMR** (500 MHz, DMSO-d6) δ 8.61 (d, *J* = 8.2 Hz, 1H), 7.84 (dd, *J* = 13.0, 9.2 Hz, 2H), 7.39 (d, *J* = 2.4 Hz, 1H), 7.35 (d, *J* = 9.6 Hz, 1H), 7.14 (dd, *J* = 8.9, 2.4 Hz, 1H), 6.85 (t, *J* = 5.6 Hz, 1H), 4.54 (td, *J* = 10.2, 5.2 Hz, 1H), 3.92 – 3.80 (m, 1H), 3.72 – 3.66 (m, 4H), 2.93 (t, *J* = 6.3 Hz, 2H), 2.47 (t, *J* = 5.1 Hz, 4H), 2.31 (t, *J* = 6.7 Hz, 2H), 2.14 – 2.07 (m, 2H), 1.93 – 1.86 (m, 2H), 1.70 – 1.58 (m, 2H), 1.57 – 1.39 (m, 7H), 1.37 (s, 9H). **HRMS** (ESI) m/z: $[M+H]^+$ calc for $C_{31}H_{43}ClN_7O_4^+$ : 612.3065; found: 612.3089

***tert*-butyl (6-(4-(6-((4-(3-chloro-4-cyanophenoxy)cyclohexyl)carbamoyl)pyridazin-3-yl)piperazin-1-yl)hexyl)carbamate (13)**

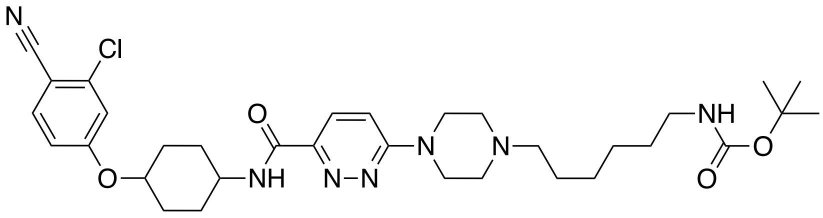

Synthesis was performed based on the general procedure **B**. Following amount of each reagent was used: N-(4-(3-chloro-4-cyanophenoxy)cyclohexyl)-6-(piperazin-1-yl)pyridazine-3-carboxamide (50 mg, 0.11 mmol), DIEA (40 μL, 0.23 mmol), and tert-butyl (6-bromohexyl)carbamate was added (38 mg, 0.14 mmol). After flash column chromatography (DCM : MeOH = 100 : 0 to 90 : 10), **13** was obtained as a white solid (40mg, 55%). $R_f$ = 0.7 (DCM : MeOH = 90 : 10, UV). **$^1$H NMR** (600 MHz, DMSO-d6) δ 8.59 (d, J = 8.2 Hz, 1H), 7.84 (dd, J = 14.6, 9.1 Hz, 2H), 7.38 (d, J = 2.4 Hz, 1H), 7.34 (d, J = 9.6 Hz, 1H), 7.13 (dd, J = 8.7, 2.4 Hz, 1H), 6.74 (t, J = 5.7 Hz, 1H), 4.53 (tt, J = 10.2, 4.6 Hz, 1H), 3.94 – 3.83 (m, 1H), 3.69 (d, J = 5.1 Hz, 4H), 2.89 (t, J = 6.8 Hz,

2H), 2.46 (t, J = 5.1 Hz, 4H), 2.30 (t, J = 7.4 Hz, 2H), 2.14 – 2.04 (m, 2H), 1.94 – 1.84 (m, 2H), 1.71 – 1.57 (m, 2H), 1.52 (td, J = 13.3, 6.9 Hz, 2H), 1.48 – 1.42 (m, 2H), 1.39 – 1.35 (m, 12H), 1.30 – 1.22 (m, 4H). **LC/MS** $[M+H]^+$ m/z calc: 640.3; found: 640.3

***tert*-butyl (7-(4-(6-((4-(3-chloro-4-cyanophenoxy)cyclohexyl)carbamoyl)pyridazin-3-yl)piperazin-1-yl)heptyl)carbamate (14)**

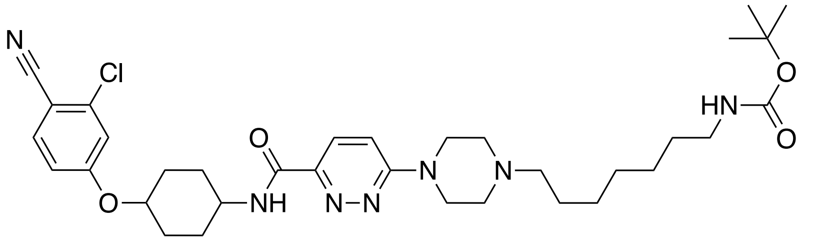

Reaction was performed based on the general procedure **B**. Following amount of each reagent was used: N-(4-(3-chloro-4-cyanophenoxy)cyclohexyl)-6-(piperazin-1-yl)pyridazine-3-carboxamide (50 mg, 0.11 mmol), DIEA (40 μL, 0.23 mmol), and *tert*-butyl (7-bromoheptyl)carbamate was added (40 mg, 0.14 mmol). After flash column chromatography (DCM : MeOH = 100 : 0 to 90 : 10), **14** was obtained as a white solid (35mg, 48%). $R_f$ = 0.7 (DCM : MeOH = 90 : 10, UV). **$^1$H NMR** 1H NMR (600 MHz, DMSO-d6) δ 8.59 (d, J = 8.2 Hz, 1H), 7.94 – 7.71 (m, 2H), 7.38 (d, J = 2.5 Hz, 1H), 7.34 (d, J = 9.6 Hz, 1H), 7.13 (dd, J = 8.8, 2.5 Hz, 1H), 6.74 (t, J = 5.6 Hz, 1H), 4.53 (tt, J = 10.5, 4.3 Hz, 1H), 3.86 (tdt, J = 11.6, 8.1, 4.0 Hz, 1H), 3.75 – 3.63 (m, 4H), 2.89 (q, J = 6.6 Hz, 2H), 2.47 (t, J = 5.2 Hz, 4H), 2.30 (d, J = 7.6 Hz, 2H), 2.15 – 2.06 (m, 2H), 1.90 (dt, J = 13.4, 3.8 Hz, 2H), 1.64 (qd, J = 13.1, 3.2 Hz, 2H), 1.52 (td, J = 11.7, 3.4 Hz, 2H), 1.46 (dt, J = 14.3, 7.7 Hz, 3H), 1.40 – 1.30 (m, 11H), 1.30 – 1.20 (m, 6H). **HRMS** (ESI) m/z: $[M+H]^+$ calc for $C_{34}H_{49}ClN_7O_4^+$ : 654.3534; found: 654.3558

6-(4-(7-(4-(3-((1-acryloylpiperidin-4-yl)methyl)phenoxy)butanamido)heptyl)piperazin-1-yl)-*N*-(4-(3-chloro-4-cyanophenoxy)cyclohexyl)pyridazine-3-carboxamide (**SJH1-62B**)

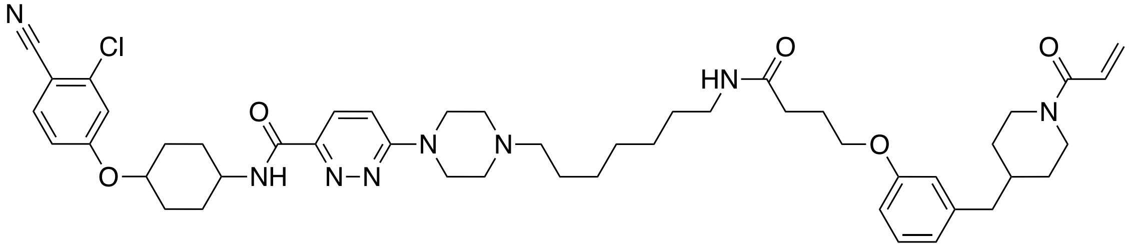

**14** (30 mg,0.05 mmol) was dissolved in 5 mL of DCM:TFA (50 : 50). After 1 hour of stirring at 23 °C, solvent was removed under vacuum. Reactant was redissolved in 5 mL of DCM and the DCM was removed under vacuum in order to remove the residual TFA. TFA removal step was repeated with additional 5 mL of DCM. Next, obtained oil was dissolved in 1 mL DMF with the following reagents: HATU (35 mg, 0.09 mmol), DIEA (40 μL, 0.23 mmol), **3** (18 mg, 0.06 mmol). Reaction was then carried based on the general procedure **C**. After flash column chromatography (DCM : MeOH = 100 : 0 to 85 : 15), another round of purification was performed using RP-HPLC (buffer B : buffer A = 40:60 to 80:20). Fractions were collected and freeze-dried to yield **SJH1-62B** as a white powder (7mg, 18%). **$^1$H NMR** (600 MHz, DMSO-d6) δ 8.67 (d, *J* = 8.2 Hz, 1H), 7.95 (d, *J* = 9.4 Hz, 1H), 7.86 (d, *J* = 8.8 Hz, 1H), 7.82 (t, *J* = 5.6 Hz, 1H), 7.49 (d, *J* = 9.6 Hz, 1H), 7.38 (d, *J* = 2.4 Hz, 1H), 7.17 (t, *J* = 7.7 Hz, 1H), 7.13 (dd, *J* = 8.8, 2.5 Hz, 1H), 6.80 – 6.69 (m, 4H), 6.06 (dd, *J* = 16.7, 2.4 Hz, 1H), 5.63 (dd, *J* = 10.5, 2.4 Hz, 1H), 4.60 (d, *J* = 14.1 Hz, 2H), 4.53 (tt, *J* = 10.4, 4.2 Hz, 1H), 4.38 (d, *J* = 13.0 Hz, 1H), 4.00 (d, *J* = 13.7 Hz, 1H), 3.92 (t, *J* = 6.4 Hz, 2H), 3.90 – 3.83 (m, 1H), 3.62 (d, *J* = 12.1 Hz, 2H), 3.37 (t, *J* = 12.9 Hz, 2H), 3.16 – 3.06 (m, 4H), 3.06 – 3.01 (m, 2H), 2.97 (t, *J* = 12.8 Hz, 1H), 2.59 – 2.52 (m, 1H), 2.47 (d, *J* = 7.2 Hz, 2H), 2.22 (t, *J* = 7.4 Hz, 2H), 2.14 – 2.08 (m, 2H), 1.95 – 1.85 (m, 4H), 1.80 – 1.73 (m, 1H), 1.71 – 1.62 (m, 4H), 1.62 – 1.56 (m, 2H), 1.52 (dt, *J* = 13.0, 9.5 Hz, 2H), 1.39 (p, *J* = 7.2 Hz, 2H), 1.32 – 1.22 (m, 6H), 1.10 – 0.96 (m, 2H). **$^{13}$C NMR** (151 MHz, DMSO-d6) δ 171.24, 163.97, 162.15, 161.70, 159.52, 158.42, 145.76, 141.51, 136.92, 135.66, 129.03, 128.50, 126.67 (d, *J* = 8.7 Hz), 121.15, 116.71, 116.30, 115.39, 115.12, 113.48, 111.59, 103.15, 75.49, 66.64, 55.46, 50.12, 47.08, 45.09, 41.91, 41.51, 38.26, 37.18, 32.36,

31.65, 31.27, 29.76, 29.28, 28.96, 28.03, 26.02, 25.81, 24.89, 22.99. **HRMS** (ESI) m/z: $[M+H]^+$ calc for $C_{48}H_{64}ClN_8O_5^+$ : 867.4688; found: 867.4727

N-(4-(3-chloro-4-cyanophenoxy)cyclohexyl)-6-(4-(7-(4-(3-((1-propionylpiperidin-4 yl)methyl)phenoxy) butanamido)heptyl)piperazin-1-yl)pyridazine-3-carboxamide (**SJH1-62B-NC**)

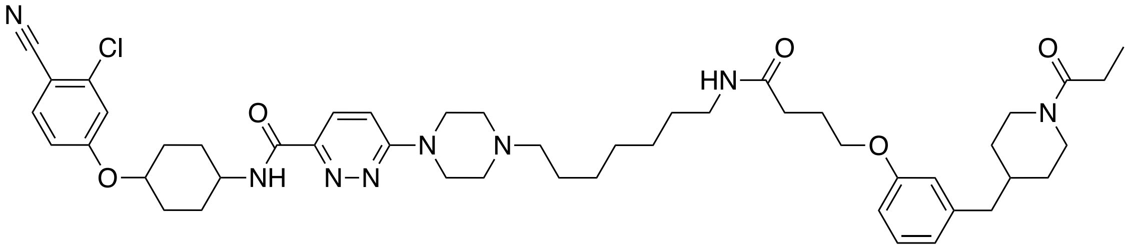

**14** (30 mg,0.05 mmol) was dissolved in 5 mL of DCM:TFA (50 : 50). After 1 hour of stirring at 23 °C, solvent was removed under vacuum. Reactant was redissolved in 5 mL of DCM and the DCM was removed under vacuum in order to remove the residual TFA. TFA removal step was repeated with additional 5 mL of DCM. Next, obtained oil was dissolved in 1 mL DMF with the following reagents: HATU (35 mg, 0.09 mmol), DIEA (40 μL, 0.23 mmol), **4** (18 mg, 0.06 mmol). Reaction was then carried based on the general procedure **C**. After flash column chromatography (DCM : MeOH = 100 : 0 to 85 : 15), another round of purification was performed using RP-HPLC (buffer B : buffer A = 40:60 to 80:20). Fractions were collected and freeze-dried to yield **SJH1-62B** as a white powder (7mg, 18%) **$^1$H NMR** (600 MHz, DMSO-d6) δ 8.66 (d, J = 8.2 Hz, 1H), 7.95 (d, J = 9.5 Hz, 1H), 7.86 (d, J = 8.8 Hz, 1H), 7.81 (t, J = 5.6 Hz, 1H), 7.49 (d, J = 9.5 Hz, 1H), 7.38 (d, J = 2.4 Hz, 1H), 7.17 (t, J = 7.8 Hz, 1H), 7.13 (dd, J = 8.8, 2.5 Hz, 1H), 6.76 – 6.68 (m, 3H), 4.60 (d, J = 14.2 Hz, 2H), 4.54 (td, J = 10.3, 5.3 Hz, 1H), 4.34 (d, J = 13.0 Hz, 1H), 3.92 (t, J = 6.4 Hz, 2H), 3.89 – 3.84 (m, 1H), 3.80 (d, J = 13.6 Hz, 2H), 3.62 (d, J = 12.2 Hz, 2H), 3.36 (t, J = 13.0 Hz, 2H), 3.17 – 3.07 (m, 4H), 3.04 (q, J = 6.6 Hz, 2H), 2.94 – 2.86 (m, 1H), 2.48 – 2.40 (m, 3H), 2.31 – 2.25 (m, 2H), 2.22 (t, J = 7.4 Hz, 2H), 2.15 – 2.08 (m, 2H), 1.96 – 1.84 (m, 4H), 1.76 – 1.69 (m, 1H), 1.69 – 1.62 (m, 3H), 1.61 – 1.46 (m, 4H), 1.44 – 1.34 (m, 2H), 1.33 – 1.21 (m, 6H), 1.06 (qd, J = 12.5, 4.1 Hz, 2H), 0.96 (t, J = 7.4 Hz, 3H). **$^{13}$C NMR** (151 MHz, DMSO-d6) δ 171.29, 170.86, 162.20, 161.76, 159.57, 158.46, 145.80, 141.61, 136.97, 135.72, 129.08, 126.71, 121.21, 116.77, 115.45, 115.17, 111.63, 103.20, 75.54, 66.68, 55.52, 50.20, 47.13, 44.81, 42.07, 41.58, 41.08, 38.31, 37.28, 32.16, 31.71, 31.36, 29.81, 29.34, 29.02, 28.08, 26.08, 25.86, 25.55, 24.94, 23.06, 9.49. **HRMS** (ESI) m/z: $[M+H]^+$ calc for $C_{48}H_{66}ClN_8O_5^+$ : 869.4844; found: 869.4821

6-(4-(4-(4-(3-((1-acryloylpiperidin-4-yl)methyl)phenoxy)butanamido)butyl)piperazin-1-yl)-*N*-(4-(3-chloro-4-cyanophenoxy)cyclohexyl)pyridazine-3-carboxamide (**SJH1-62C**)

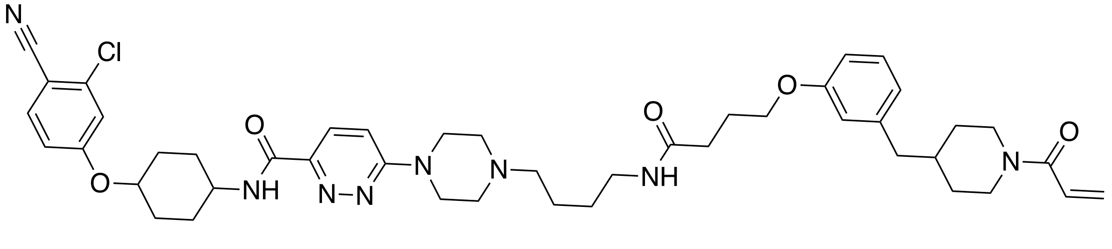

**12** (30 mg,0.05 mmol) was dissolved in 5 mL of DCM:TFA (50 : 50). After 1 hour of stirring at 23 °C, solvent was removed under vacuum. Reactant was redissolved in 5 mL of DCM and the DCM was removed under vacuum in order to remove the residual TFA. TFA removal step was repeated with additional 5 mL of DCM. Next, obtained oil was dissolved in 1 mL DMF with the following reagents: HATU (37 mg, 0.10 mmol), DIEA (43 μL, 0.25 mmol), **3** (20 mg, 0.06 mmol). Reaction was then performed based on the general procedure **C**. After flash column chromatography (DCM : MeOH = 100 : 0 to 90 : 10), another round of purification was performed using HPLC (buffer B : buffer A = 40:60 to 80:20). Targeted mass containing fractions were collected and freeze-dried to yield **SJH1-62C** as a white powder (5.5mg, 14%). **$^{1}$H NMR** (600 MHz, DMSO-D6) δ 8.67 (d, *J* = 8.2 Hz, 1H), 7.95 (d, *J* = 9.4 Hz, 1H), 7.92 (t, *J* = 5.7 Hz, 1H), 7.86 (d, *J* = 8.8 Hz, 1H), 7.49 (d, *J* = 9.5 Hz, 1H), 7.39 (d, *J* = 2.4 Hz, 1H), 7.18 (t, *J* = 7.8 Hz, 1H), 7.14 (dd, *J* = 8.8, 2.4 Hz, 1H), 6.81 – 6.70 (m, 4H), 6.06 (dd, *J* = 16.7, 2.4 Hz, 1H), 5.64 (dd, *J* = 10.5, 2.4 Hz, 1H), 4.60 (d, *J* = 14.2 Hz, 2H), 4.57 – 4.51 (m, 1H), 4.38 (d, *J* = 13.0 Hz, 1H), 4.01 (d, *J* = 13.8 Hz, 1H), 3.94 (t, *J* = 6.4 Hz, 2H), 3.88 (dtd, *J* = 11.4, 7.6, 4.1 Hz, 1H), 3.60 (d, *J* = 12.3 Hz, 2H), 3.38 (t, *J* = 13.2 Hz, 2H), 3.19 – 3.06 (m, 6H), 2.97 (t, *J* = 12.9 Hz, 1H), 2.56 (t, *J* = 13.1 Hz, 1H), 2.48 (d, *J* = 7.2 Hz, 2H), 2.25 (t, *J* = 7.5 Hz, 2H), 2.15 – 2.08 (m, 2H), 1.96 – 1.87 (m, 4H), 1.77 (dqt, *J* = 11.0, 7.1, 3.6 Hz, 1H), 1.71 – 1.64 (m, 4H), 1.63 – 1.57 (m, 2H), 1.56 – 1.48 (m, 2H), 1.48 – 1.42 (m, 2H), 1.03 (tt, *J* = 13.8, 6.8 Hz, 2H). **$^{13}$C NMR** (151 MHz, DMSO-D6) δ 171.5, 164.0, 162.1, 161.7, 159.5, 158.4, 145.8, 141.5, 136.9, 135.7, 129.0, 128.5, 126.7 (d, *J* = 9.7 Hz), 121.2, 116.7, 116.3, 115.4, 115.1, 113.5, 111.6, 103.2, 75.5, 66.6, 55.2, 50.2, 47.1, 45.1, 41.9, 41.5, 37.7, 37.2, 32.4, 31.7, 31.3, 29.8, 29.3, 26.2, 24.9, 20.7. **HRMS** (ESI) m/z: $[M+H]^+$ calc for $C_{45}H_{58}ClN_8O_5^+$ : 825.4218; found: 825.4193

6-(4-(6-(4-(3-((1-acryloylpiperidin-4-yl)methyl)phenoxy)butanamido)hexyl)piperazin-1-yl)-*N*-(4-(3-chloro-4-cyanophenoxy)cyclohexyl)pyridazine-3-carboxamide (**SJH1-62D**)

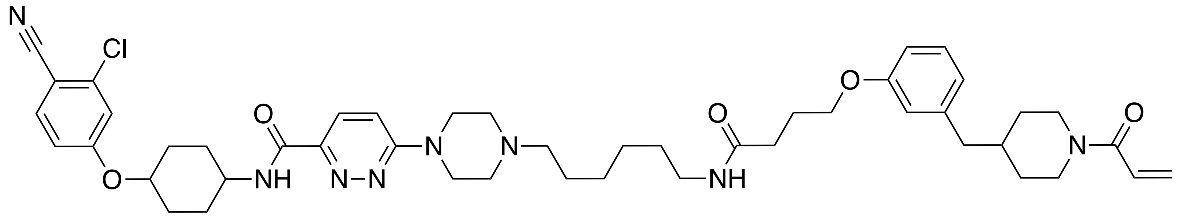

**13** (30 mg,0.05 mmol) was dissolved in 5 mL of DCM:TFA (50 : 50). After 1 hour of stirring at 23 °C, solvent was removed under vacuum. Reactant was redissolved in 5 mL of DCM and the DCM was removed under vacuum in order to remove the residual TFA. TFA removal step was repeated with additional 5 mL of DCM. Next, obtained oil was dissolved in 1 mL DMF with the following reagents: HATU (36 mg, 0.09 mmol), DIEA (41 μL, 0.23 mmol), **3** (19 mg, 0.06 mmol). Reaction was then carried out based on the general procedure **C**. After flash column chromatography (DCM : MeOH = 100 : 0 to 90 : 10), another round of purification was performed using HPLC (buffer B : buffer A = 40:60 to 80:20). Fractions were collected and freeze-dried to yield **SJH1-62D** as a white powder (10.1mg, 28%) **$^{1}$H NMR** (600 MHz, DMSO-D6) δ 8.67 (d, *J* = 8.2 Hz, 1H), 7.95 (d, *J* = 9.5 Hz, 1H), 7.86 (d, *J* = 8.8 Hz, 1H), 7.83 (d, *J* = 5.6 Hz, 1H), 7.49 (d, *J* = 9.5 Hz, 1H), 7.38 (d, *J* = 2.4 Hz, 1H), 7.17 (t, *J* = 7.7 Hz, 1H), 7.13 (dd, *J* = 8.8, 2.5 Hz, 1H), 6.80 – 6.68 (m, 4H), 6.06 (dd, *J* = 16.7, 2.4 Hz, 1H), 5.63 (dd, *J* = 10.5, 2.4 Hz, 1H), 4.60 (d, *J* = 14.2 Hz, 2H), 4.53 (tt, *J* = 10.4, 4.3 Hz, 1H), 4.38 (d, *J* = 13.0 Hz, 1H), 4.00 (d, *J* = 13.7 Hz, 1H), 3.93 (t, *J* = 6.4 Hz, 2H), 3.87 (dtd, *J* = 11.4, 7.6, 4.1 Hz, 1H), 3.62 – 3.58 (m, 2H), 3.37 (t, *J* = 13.1 Hz, 2H), 3.15 – 3.07 (m, 4H), 3.05 (q, *J* = 6.6 Hz, 2H), 2.97 (t, *J* = 12.8 Hz, 1H), 2.56 (t, *J* = 12.4 Hz, 1H), 2.47 (d, *J* = 7.2 Hz, 2H), 2.23 (t, *J* = 7.4 Hz, 2H), 2.14 – 2.08 (m, 2H), 1.94 – 1.87 (m, 4H), 1.77 (dqd, *J* = 11.0, 7.3, 3.5 Hz, 1H), 1.70 – 1.62 (m, 4H), 1.62 – 1.56 (m, 2H), 1.52 (dt, *J* = 13.0, 9.5 Hz, 2H), 1.41 (p, *J* = 7.0 Hz, 2H), 1.33 – 1.27 (m, 4H), 1.09 – 0.97 (m, 2H). **$^{13}$C NMR** (151 MHz, DMSO-D6) δ 171.3, 164.0, 162.2, 161.7, 159.5, 158.4, 145.7, 141.5, 136.9, 135.7, 129.0, 128.5, 126.7 (d, *J* = 9.4 Hz), 121.2, 116.7, 116.3, 115.4, 115.1, 113.5, 111.6, 103.1, 75.5, 66.6, 55.5, 50.1, 47.1, 45.1, 41.9, 41.5, 38.1, 37.2, 32.4, 31.7, 31.3, 29.8, 29.3, 28.8, 25.8, 25.6, 24.9, 23.0. **HRMS** (ESI) m/z: $[M+H]^+$ calc for $C_{47}H_{62}ClN_8O_5^+$ : 853.4531; found: 853.4503

6-(4-(4-(3-((1-acryloylpiperidin-4-yl)methyl)phenoxy)butanoyl)piperazin-1-yl)-*N*-(4-(3-chloro-4-cyanophenoxy)cyclohexyl)pyridazine-3-carboxamide (**SJH1-63)**

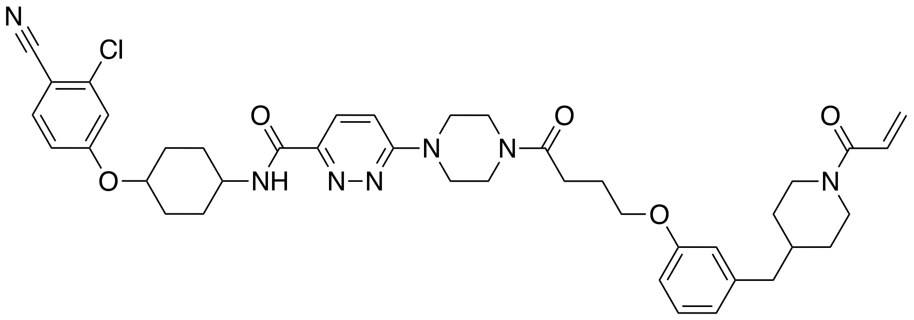


N-(4-(3-chloro-4-cyanophenoxy)cyclohexyl)-6-(piperazin-1-yl)pyridazine-3-carboxamide (50 mg, 0.11 mmol), DIEA (40 μL, 0.23 mmol), and **3** (44 mg, 0.13 mmol) was added in 2 mL DMF. Reaction was stirred vigorously at 23 °C for 4 hours. The reaction was quenched by addition of 10 mL water, and the product was extracted five times using 40 mL of 4:1 $CHCl_3$:IPA. The organic layer was dried with $Na_2SO_4$ and concentrated under vacuum. After 2 rounds of flash column chromatography (DCM:MeOH = 100 : 0 to 90:10), **SJH1-63** was obtained as a white powder (18 mg, 21 %). **$^1$H NMR** (600 MHz, DMSO-d6) δ 8.61 (d, *J* = 8.2 Hz, 1H), 7.86 (dd, *J* = 9.1, 8.4 Hz, 2H), 7.41 – 7.34 (m, 2H), 7.22 – 7.12 (m, 2H), 6.81 – 6.69 (m, 4H), 6.05 (dd, *J* = 16.7, 2.5 Hz, 1H), 5.63 (dd, *J* = 10.4, 2.4 Hz, 1H), 4.54 (tt, *J* = 10.3, 4.2 Hz, 1H), 4.37 (d, *J* = 13.1 Hz, 2H), 3.99 (t, *J* = 6.4 Hz, 3H), 3.86 (tdt, *J* = 11.6, 8.0, 4.0 Hz, 2H), 3.79 – 3.73 (m, 2H), 3.73 – 3.68 (m, 2H), 3.66 – 3.58 (m, 4H), 2.96 (t, *J* = 12.9 Hz, 1H), 2.56 – 2.52 (m, 2H), 2.47 (d, *J* = 7.2 Hz, 2H), 2.15 – 2.06 (m, 2H), 2.00 – 1.94 (m, 2H), 1.94 – 1.86 (m, 2H), 1.77 (dqd, *J* = 11.1, 7.3, 3.5 Hz, 1H), 1.70 – 1.56 (m, 4H), 1.56 – 1.45 (m, 2H), 1.07 – 0.99 (m, 2H). **$^{13}$C NMR** (151 MHz, DMSO-d6) δ 170.4, 164.0, 162.3, 161.7, 159.9, 158.4, 144.9, 141.5, 136.9, 135.7, 129.0, 128.5, 126.7, 126.3, 121.1, 116.7, 116.3, 115.4, 115.1, 112.8, 111.7, 103.1, 75.5, 66.5, 47.0, 45.1, 44.1, 44.0, 41.9, 41.5, 40.4, 37.2, 32.3, 31.3, 29.8, 29.3, 28.6, 24.4. **HRMS** (ESI) m/z: $[M+Na]^+$ calc for $C_{41}H_{48}ClN_7O_5Na^+$: 776.3303; found: 776.3300

**Synthetic References**

(1) Henning, N. J.; Boike, L.; Spradlin, J. N.; Ward, C. C.; Liu, G.; Zhang, E.; Belcher, B. P.; Brittain, S. M.; Hesse, M. J.; Dovala, D.; et al. Deubiquitinase-targeting chimeras for targeted protein stabilization. *Nature Chemical Biology* **2022**, *18* (4), 412-421. DOI: 10.1038/s41589-022-00971-2.

(2) Forte, N.; Dovala, D.; Hesse, M. J.; McKenna, J. M.; Tallarico, J. A.; Schirle, M.; Nomura, D. K. Targeted Protein Degradation through E2 Recruitment. *ACS Chemical Biology* **2023**, *18* (4), 897-904. DOI: 10.1021/acschembio.3c00040.

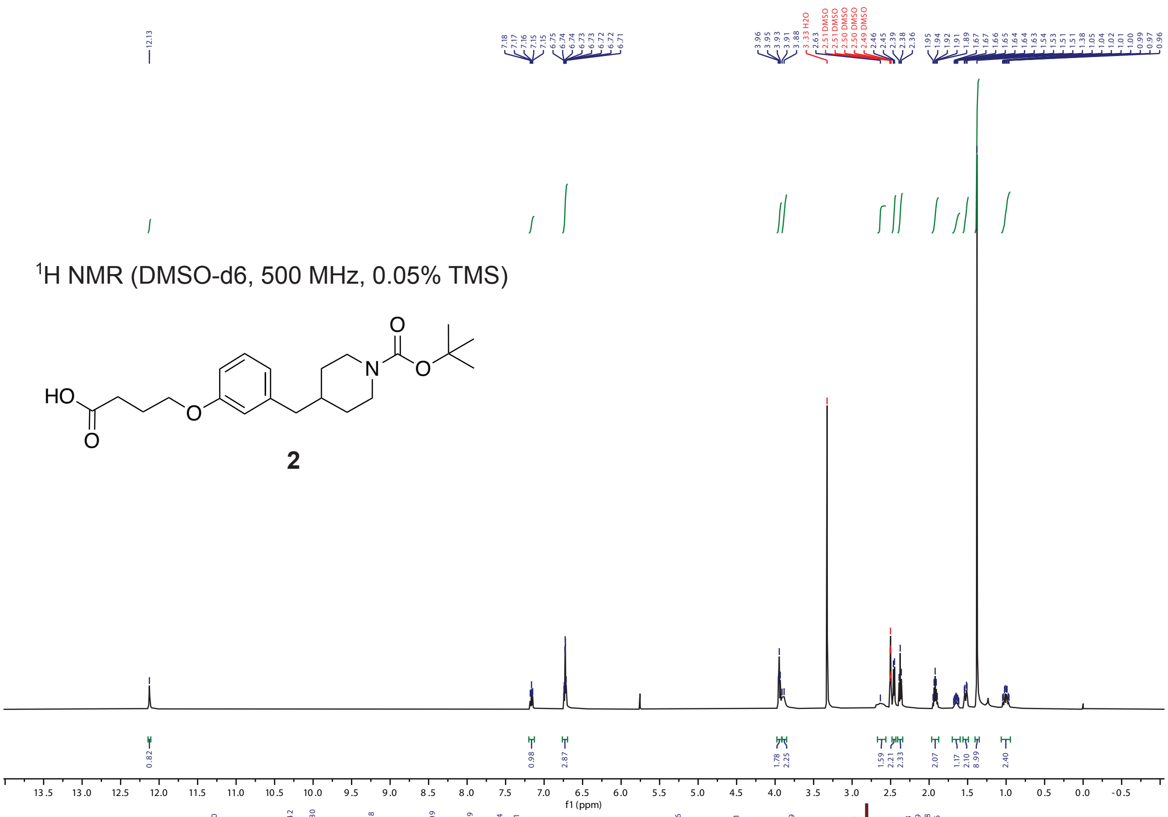


$^{1}$H NMR (DMSO-d6, 500 MHz, 0.05% TMS)

**2**

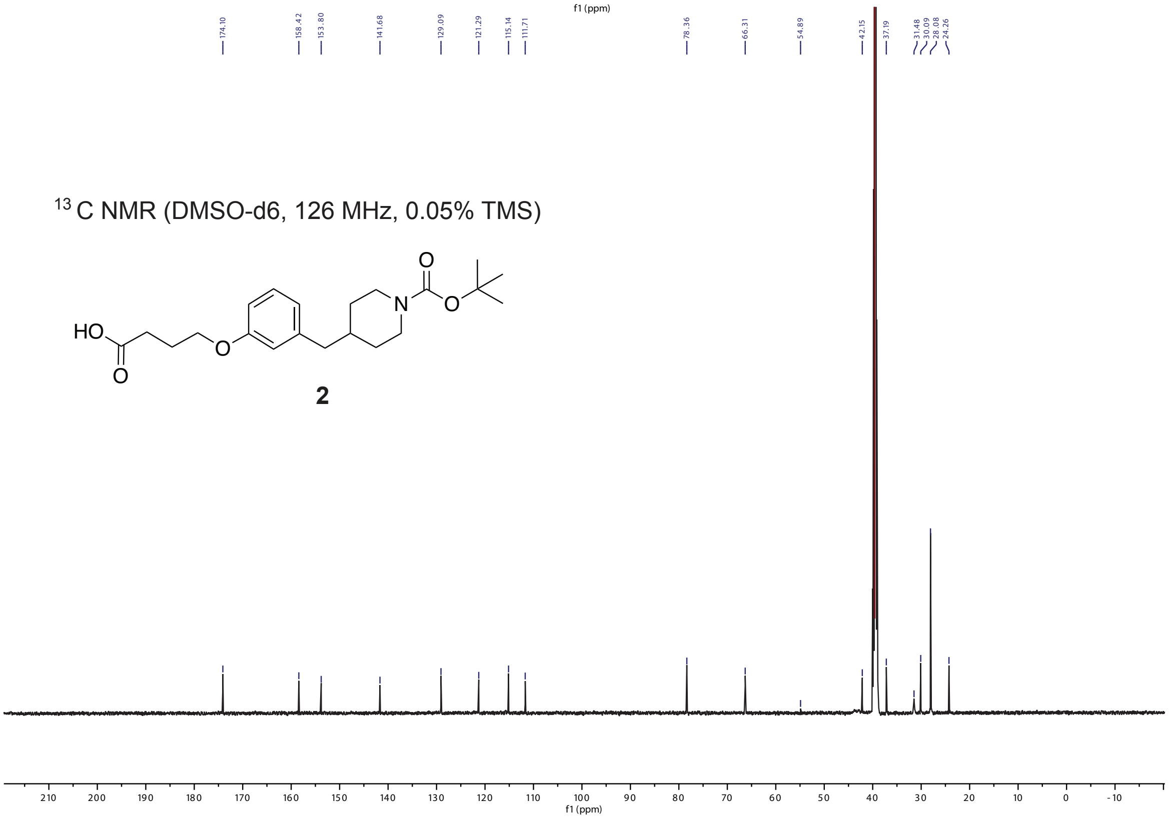


$^{13}$C NMR (DMSO-d6, 126 MHz, 0.05% TMS)

**2**

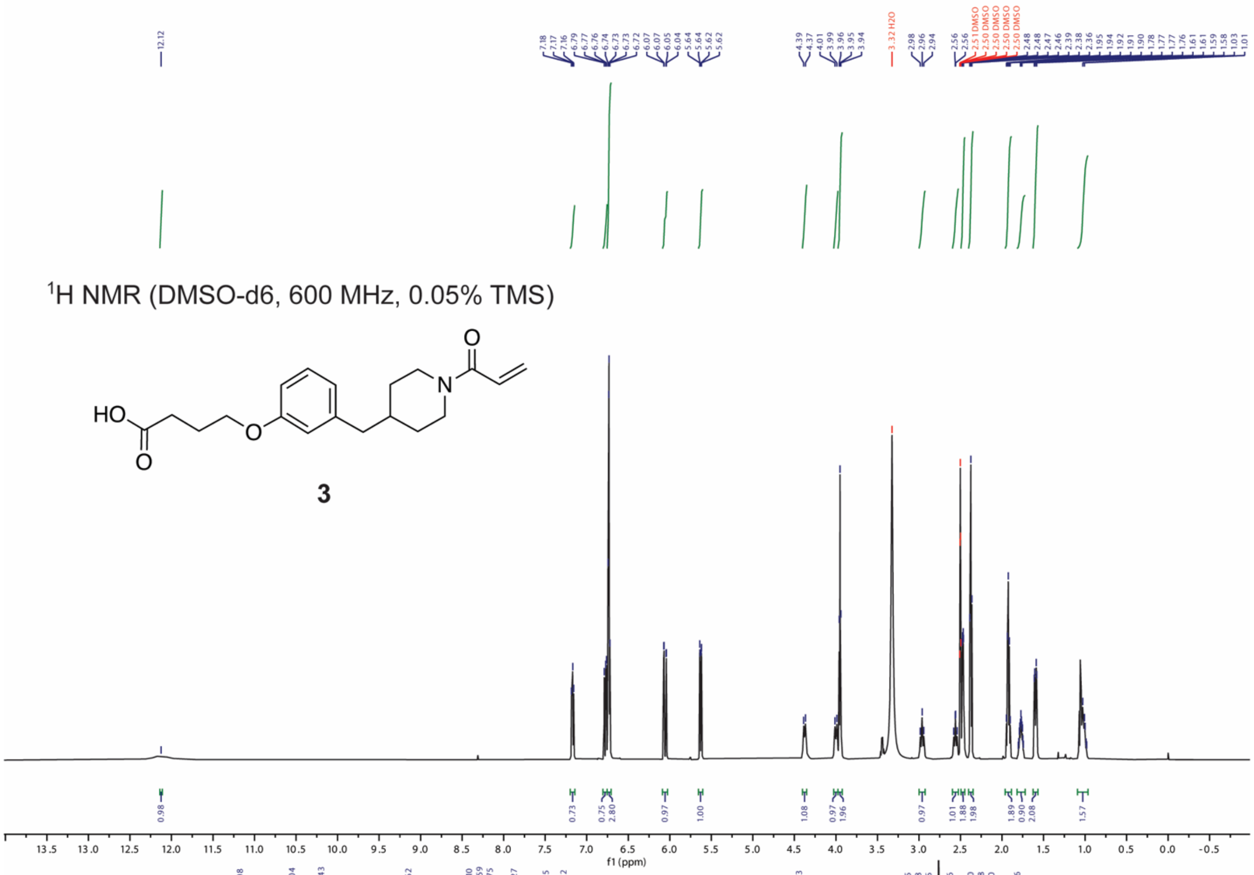

1H NMR (DMSO-d6, 600 MHz, 0.05% TMS)
HO
O
O
N
O
3
f1 (ppm)


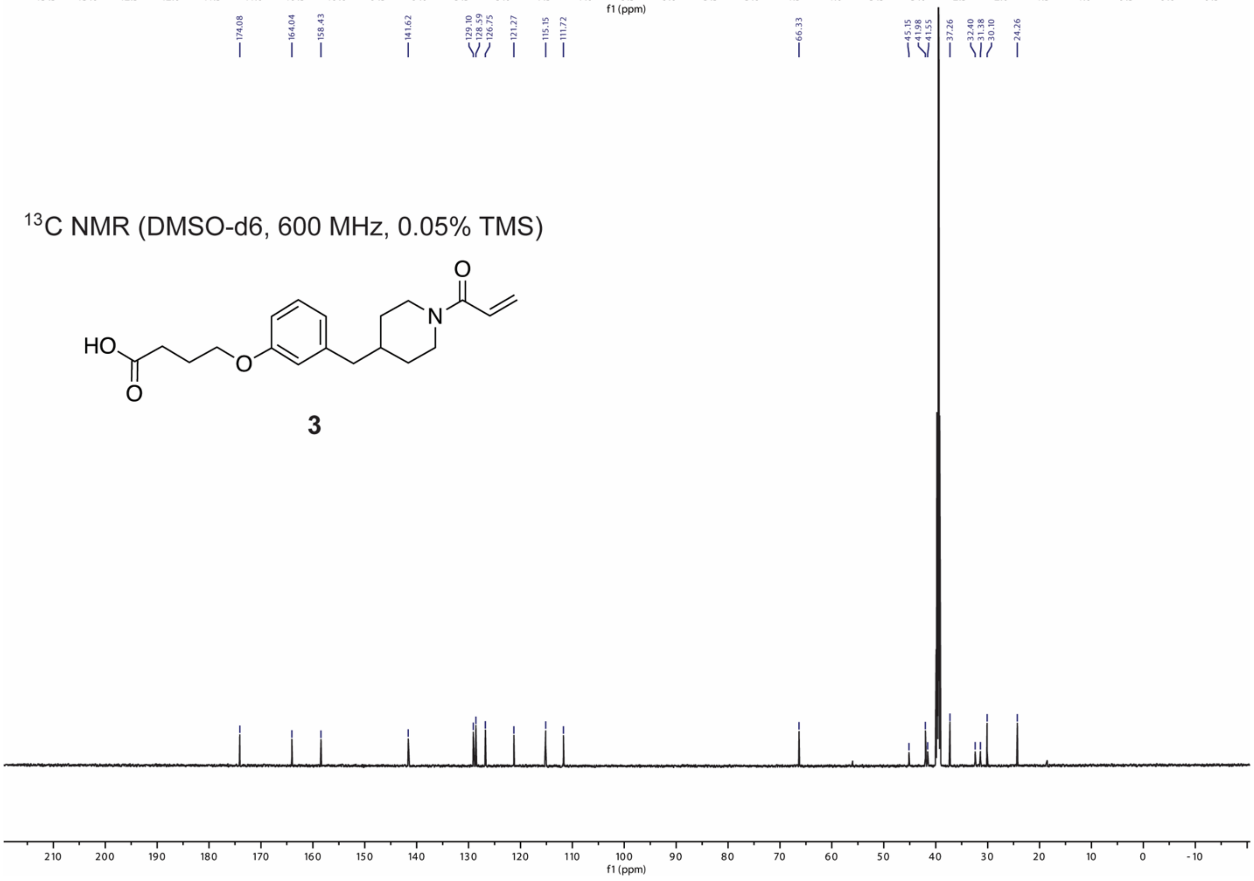

13C NMR (DMSO-d6, 600 MHz, 0.05% TMS)
HO
O
O
N
O
3
f1 (ppm)

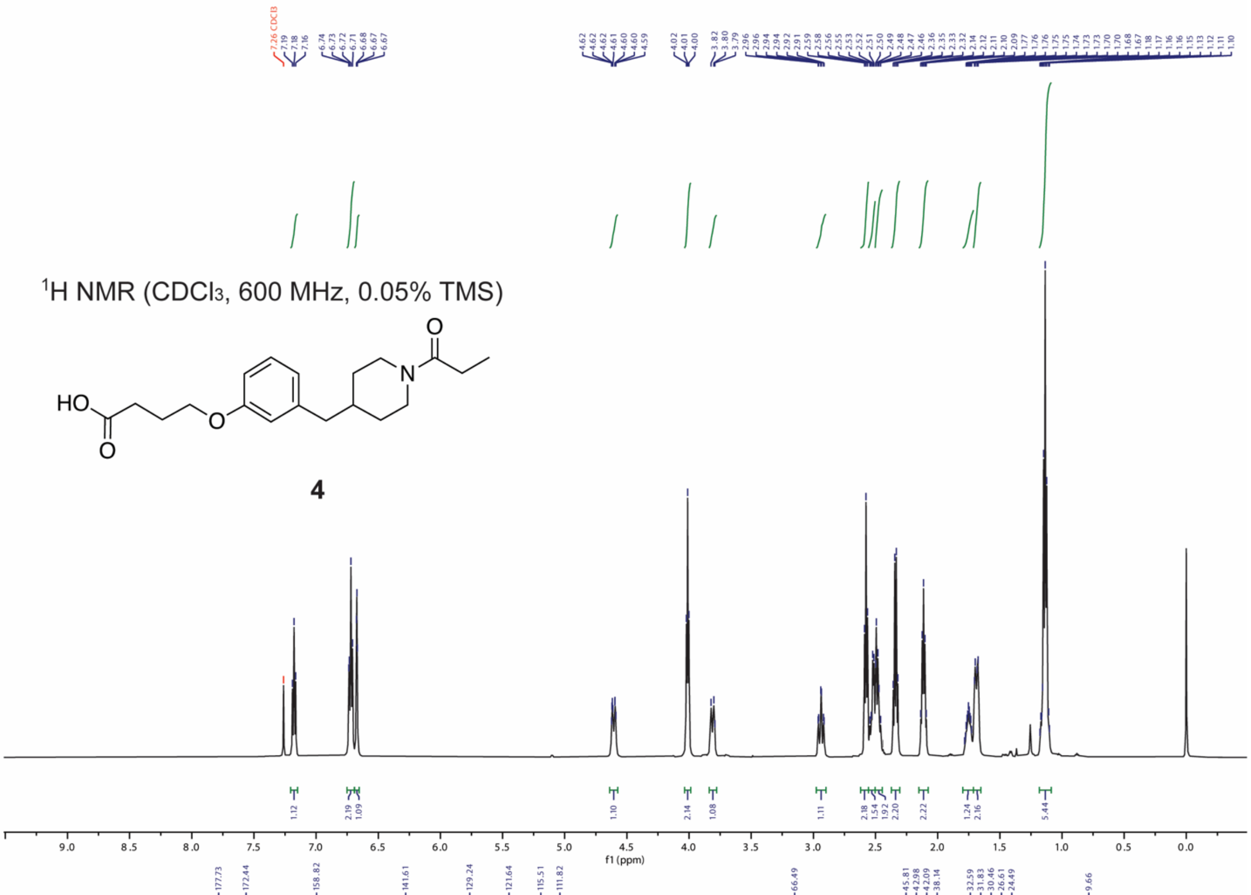

$^{1}$H NMR ($CDCl_3$, 600 MHz, 0.05% TMS)

**4**

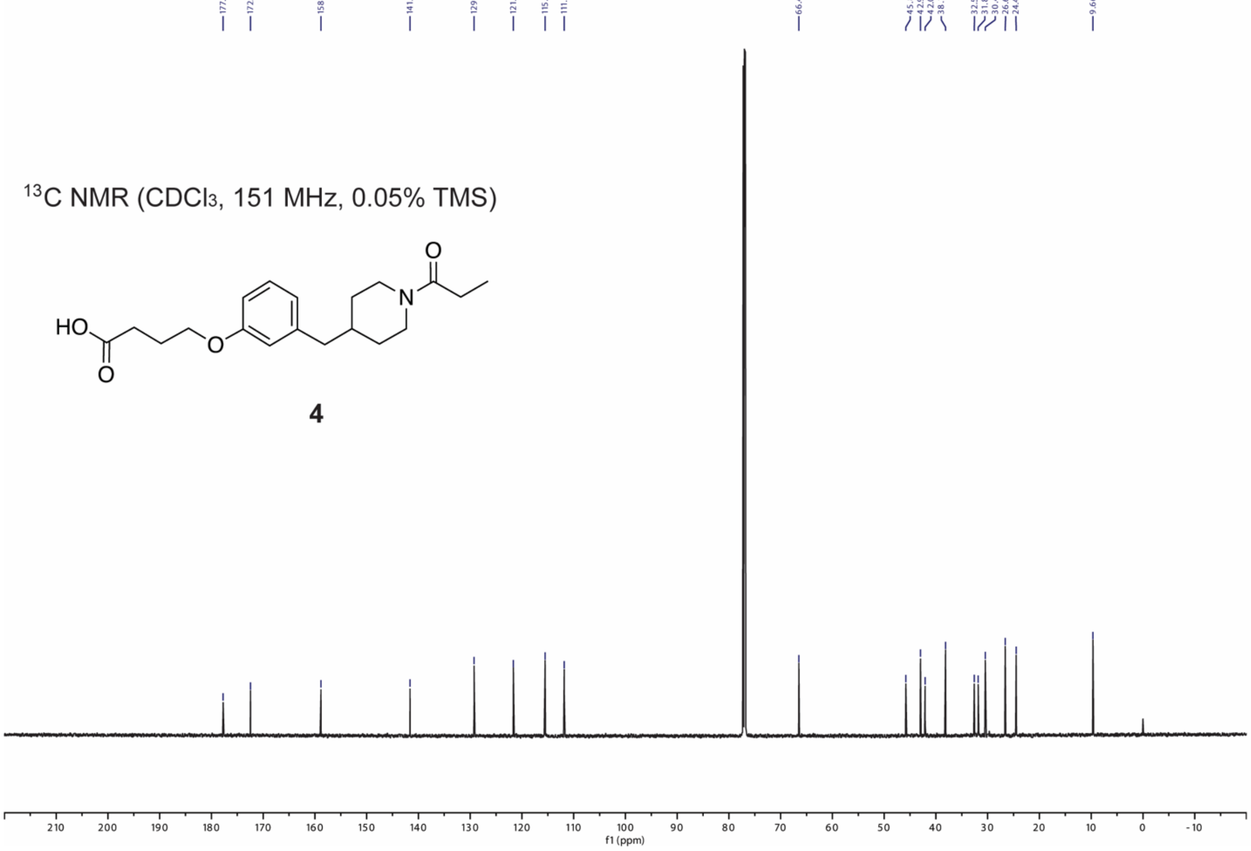

$^{13}$C NMR ($CDCl_3$, 151 MHz, 0.05% TMS)

**4**

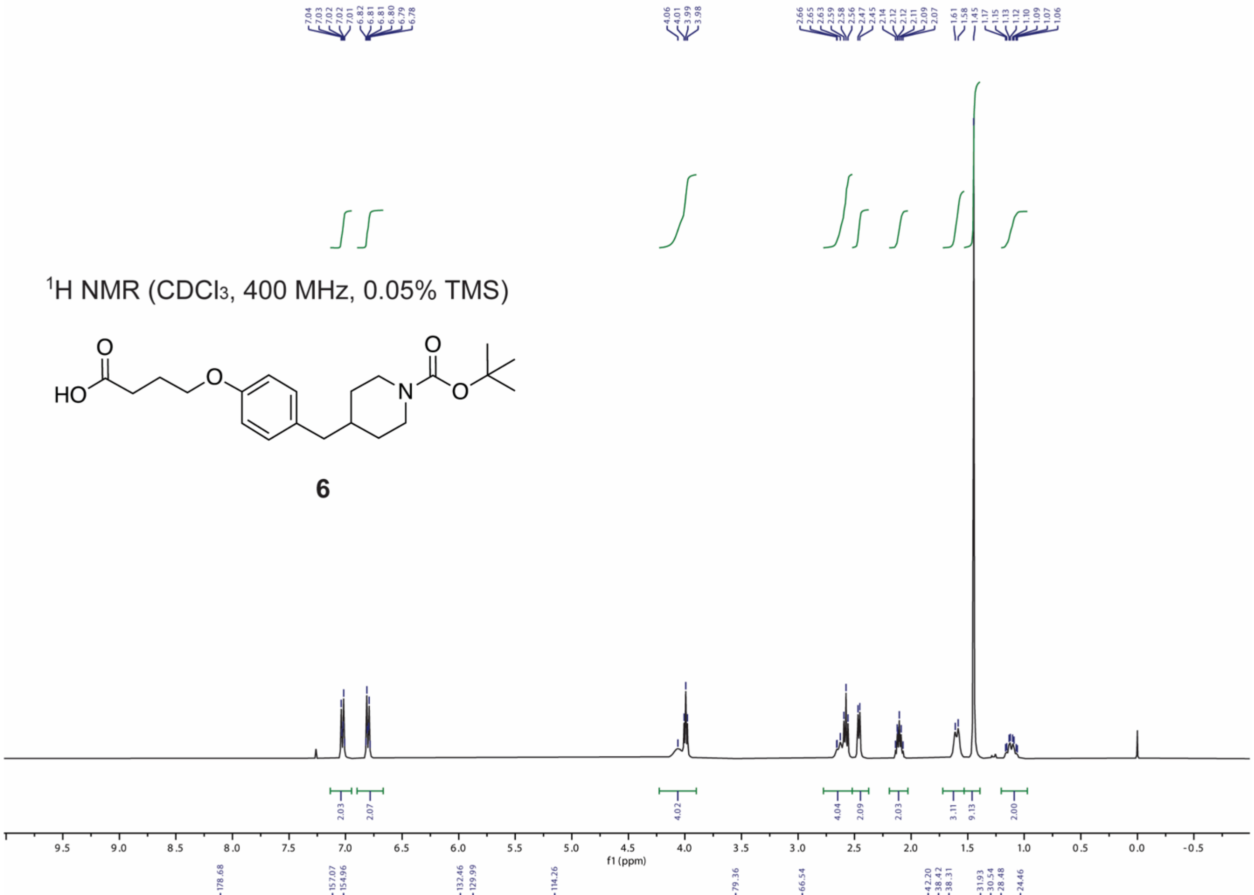

7.04
7.03
7.02
7.02
7.01
6.82
6.81
6.81
6.80
6.79
6.78
4.06
4.01
3.99
3.98
2.66
2.65
2.63
2.59
2.58
2.56
2.47
2.45
2.14
2.12
2.12
2.11
2.09
2.07
1.61
1.58
1.45
1.17
1.15
1.13
1.12
1.10
1.09
1.07
1.06
$^{1}$H NMR ($CDCl_3$, 400 MHz, 0.05% TMS)
HO
O
O
O
N
O
6
2.03
2.07
4.02
4.04
2.09
2.03
3.11
9.13
2.00
f1 (ppm)


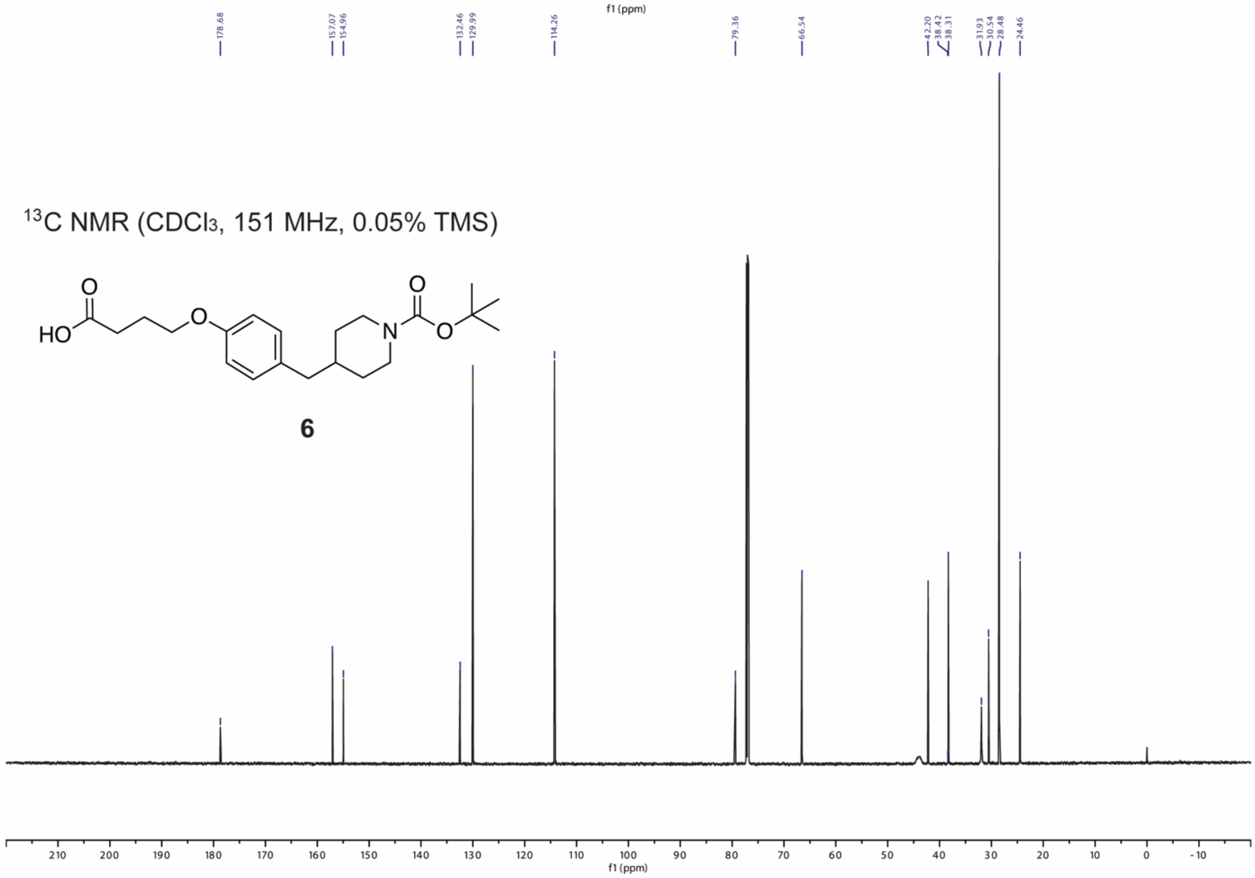

178.68
157.07
154.96
132.46
129.99
114.26
79.36
66.54
42.20
38.42
38.31
31.93
30.54
28.48
24.46
$^{13}$C NMR ($CDCl_3$, 151 MHz, 0.05% TMS)
HO
O
O
O
N
O
6
f1 (ppm)

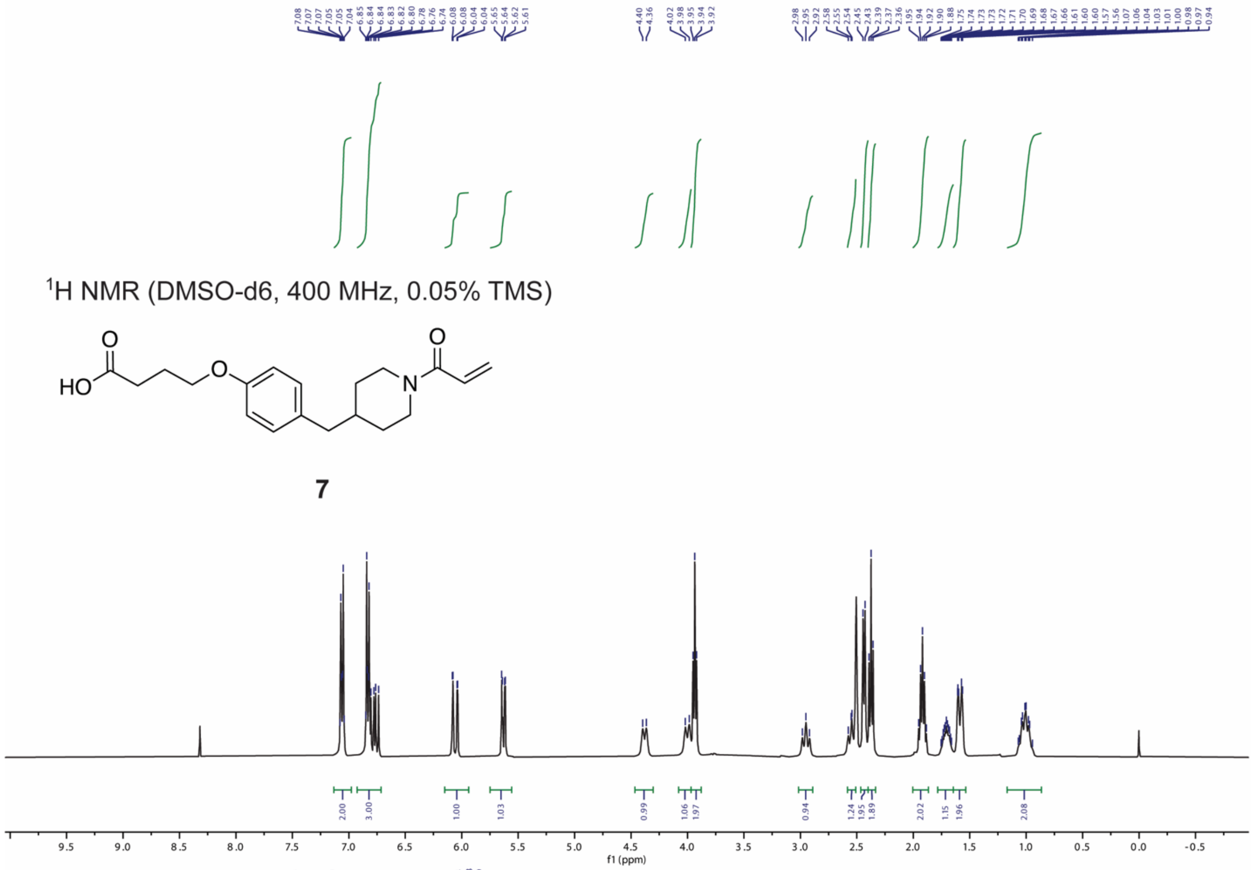


[1]H NMR (DMSO-d6, 400 MHz, 0.05% TMS)

**7**

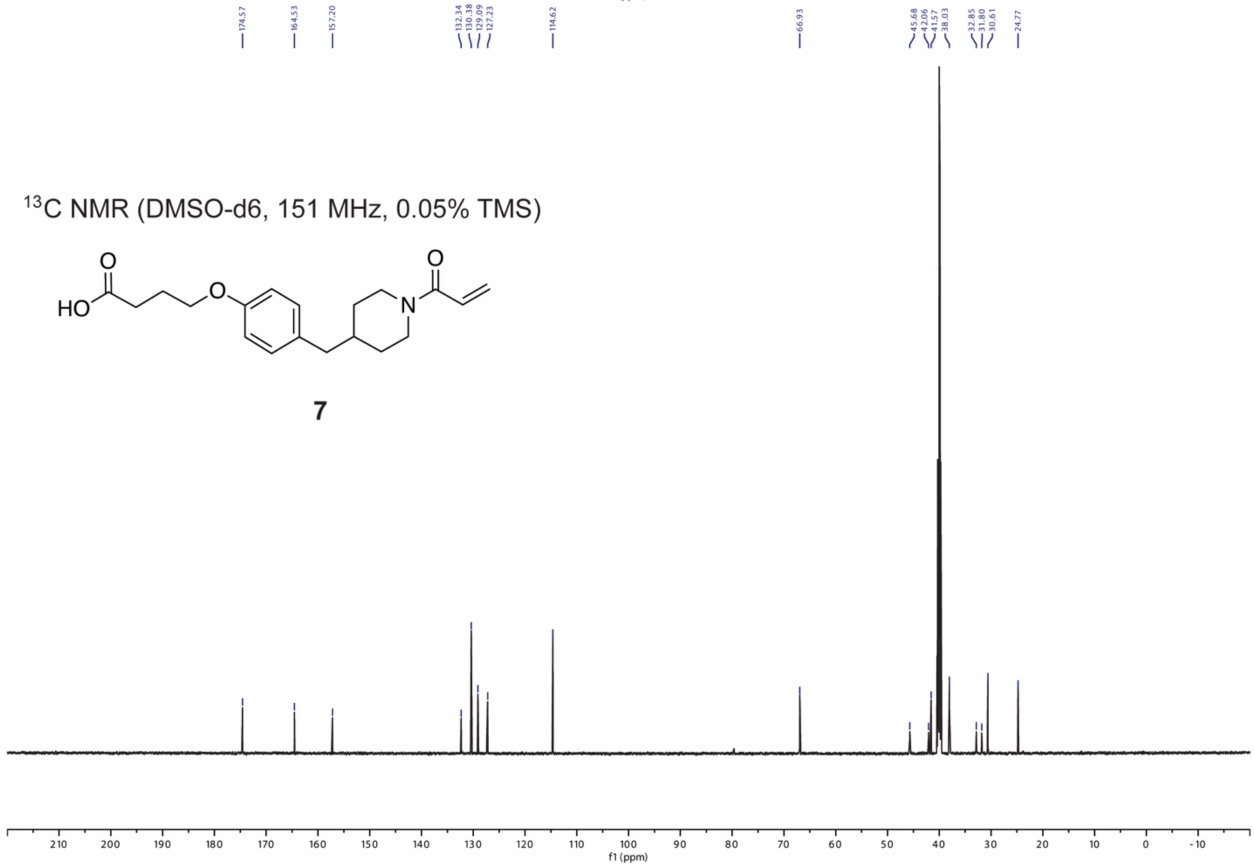


[13]C NMR (DMSO-d6, 151 MHz, 0.05% TMS)

**7**

[1]H NMR (DMSO-d6, 600 MHz, 0.05% TMS)

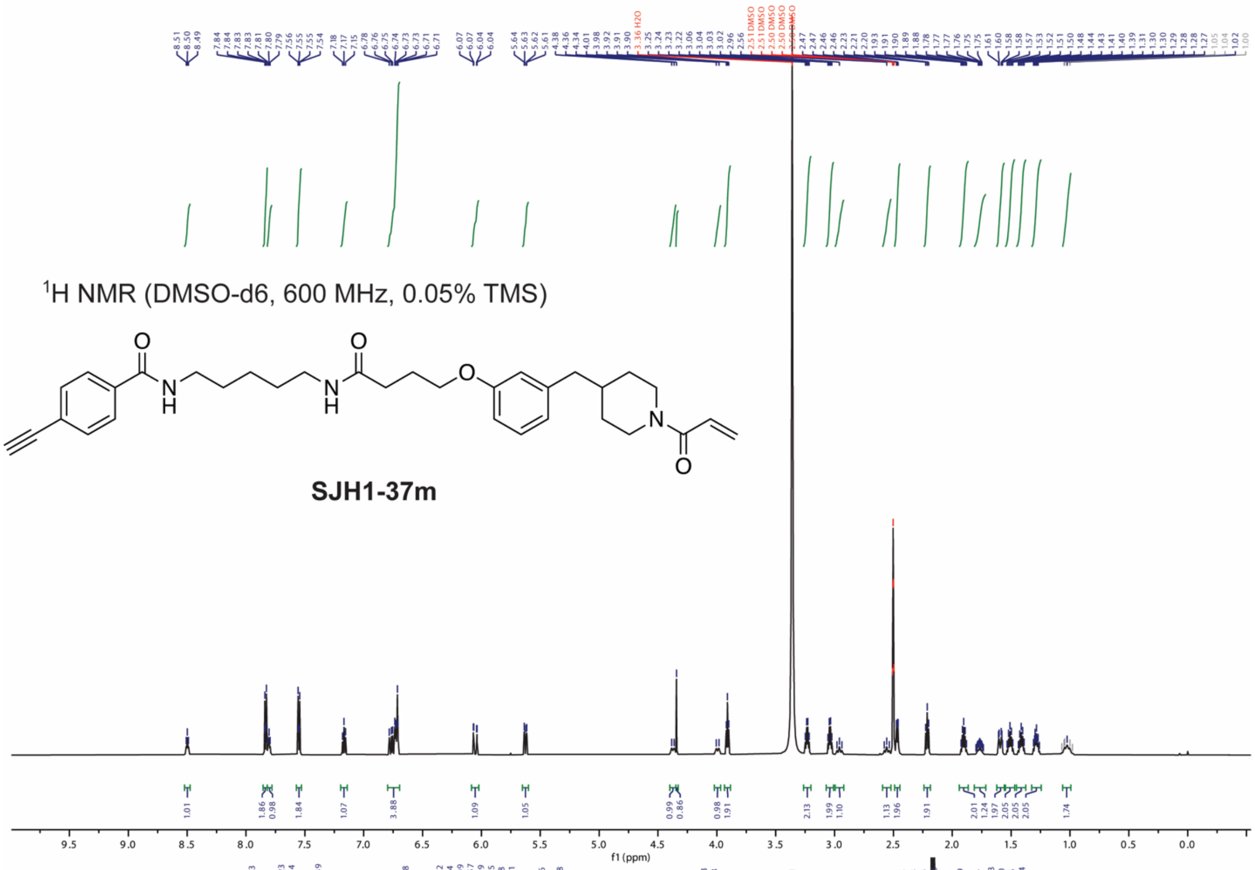


**SJH1-37m**

[13]C NMR (DMSO-d6, 151 MHz, 0.05% TMS)

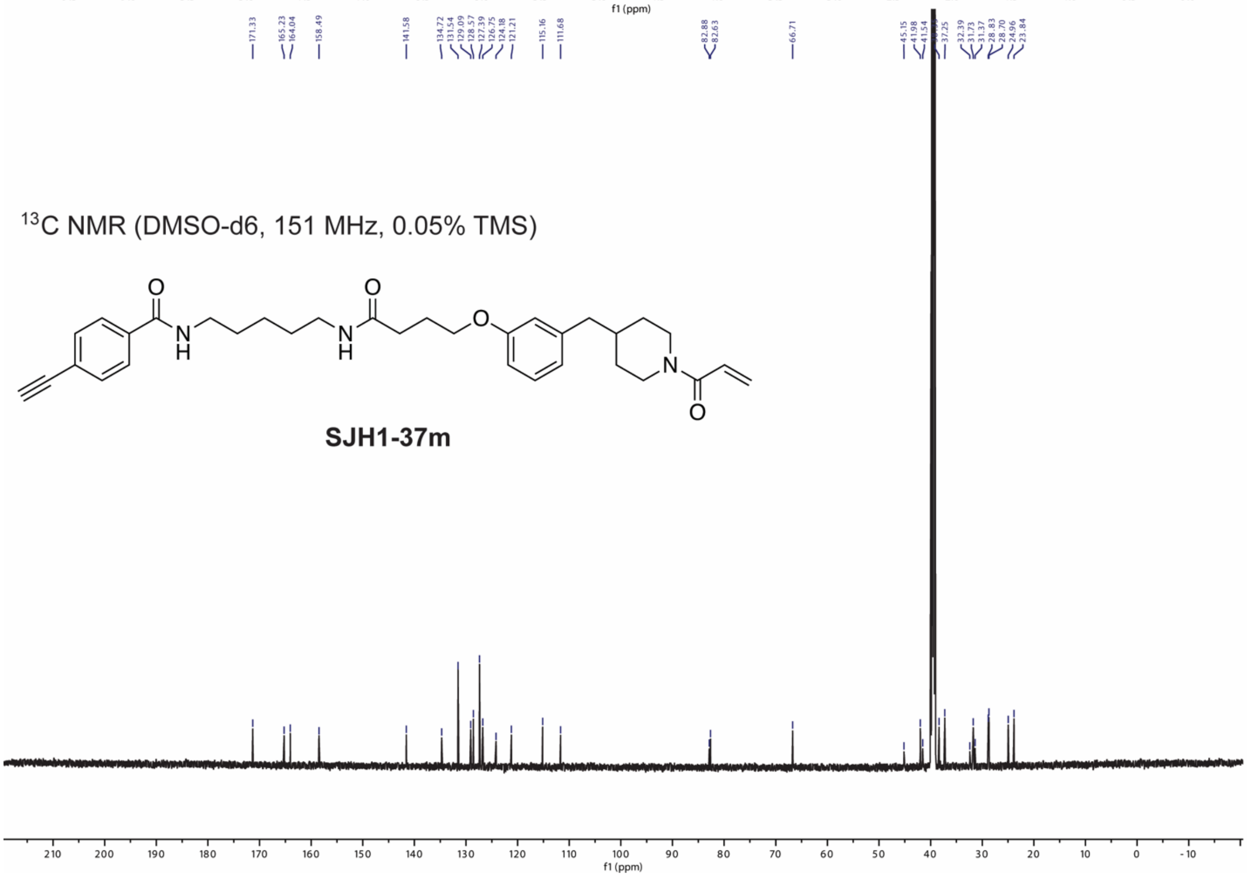


**SJH1-37m**

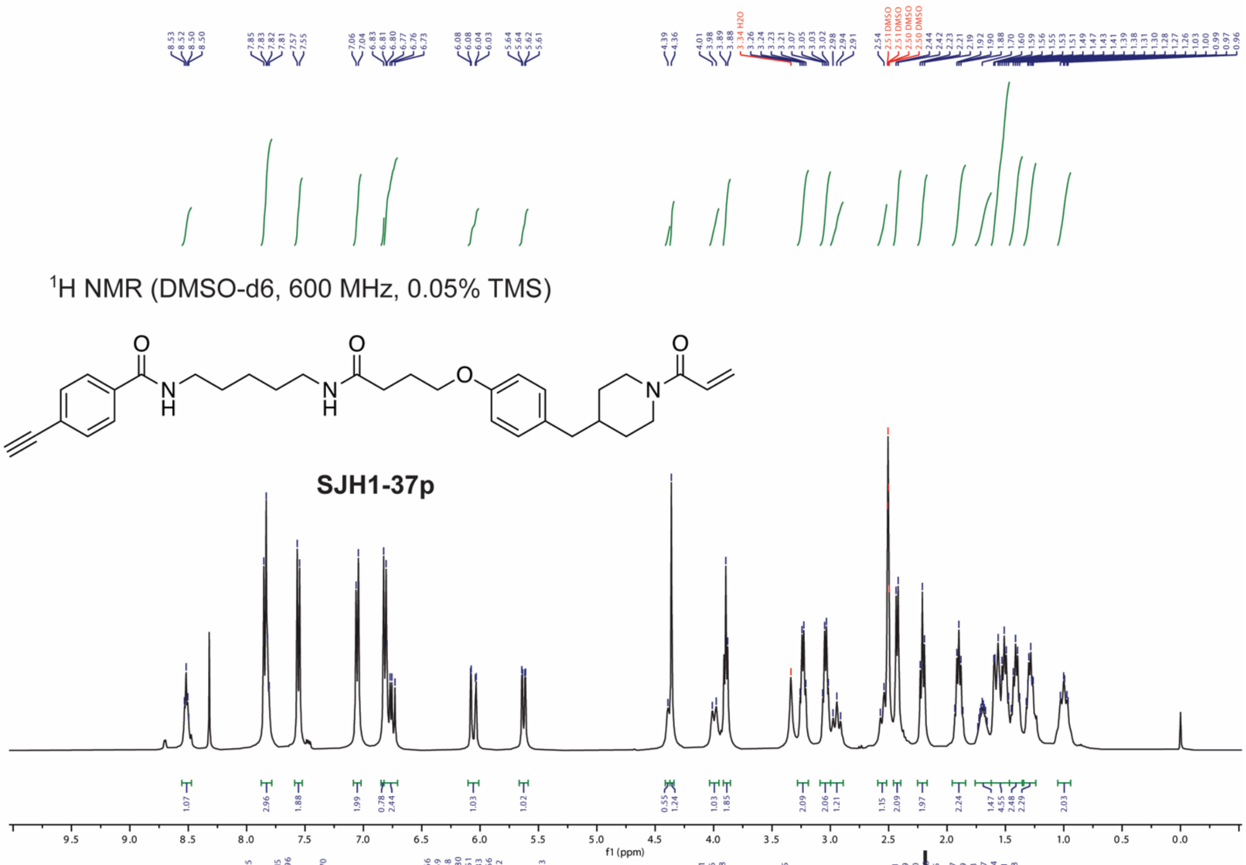
8.53 8.52 8.50 8.50 7.85 7.83 7.82 7.81 7.57 7.55 7.06 7.04 6.83 6.81 6.80 6.77 6.76 6.73 6.08 6.08 6.04 6.03 5.64 5.64 5.62 5.61 4.39 4.36 4.01 3.98 3.89 3.88 3.34 H2O 3.26 3.24 3.23 3.21 3.07 3.05 3.03 3.02 2.98 2.94 2.91 2.54 2.51 DMSO 2.51 DMSO 2.50 DMSO 2.50 DMSO 2.44 2.42 2.23 2.21 2.19 1.92 1.90 1.88 1.70 1.60 1.59 1.56 1.55 1.53 1.51 1.49 1.47 1.43 1.41 1.39 1.38 1.31 1.30 1.28 1.27 1.26 1.03 1.00 0.99 0.97 0.96
$^1$H NMR (DMSO-d6, 600 MHz, 0.05% TMS)
O
N
H
O
N
H
O
O
N
SJH1-37p
1.07 2.96 1.88 1.99 0.78 2.44 1.03 1.02 0.55 1.24 1.03 1.85 2.09 2.06 1.21 1.15 2.09 1.97 2.24 1.47 4.55 2.48 2.29 2.03
9.5 9.0 8.5 8.0 7.5 7.0 6.5 6.0 5.5 5.0 4.5 4.0 3.5 3.0 2.5 2.0 1.5 1.0 0.5 0.0
f1 (ppm)

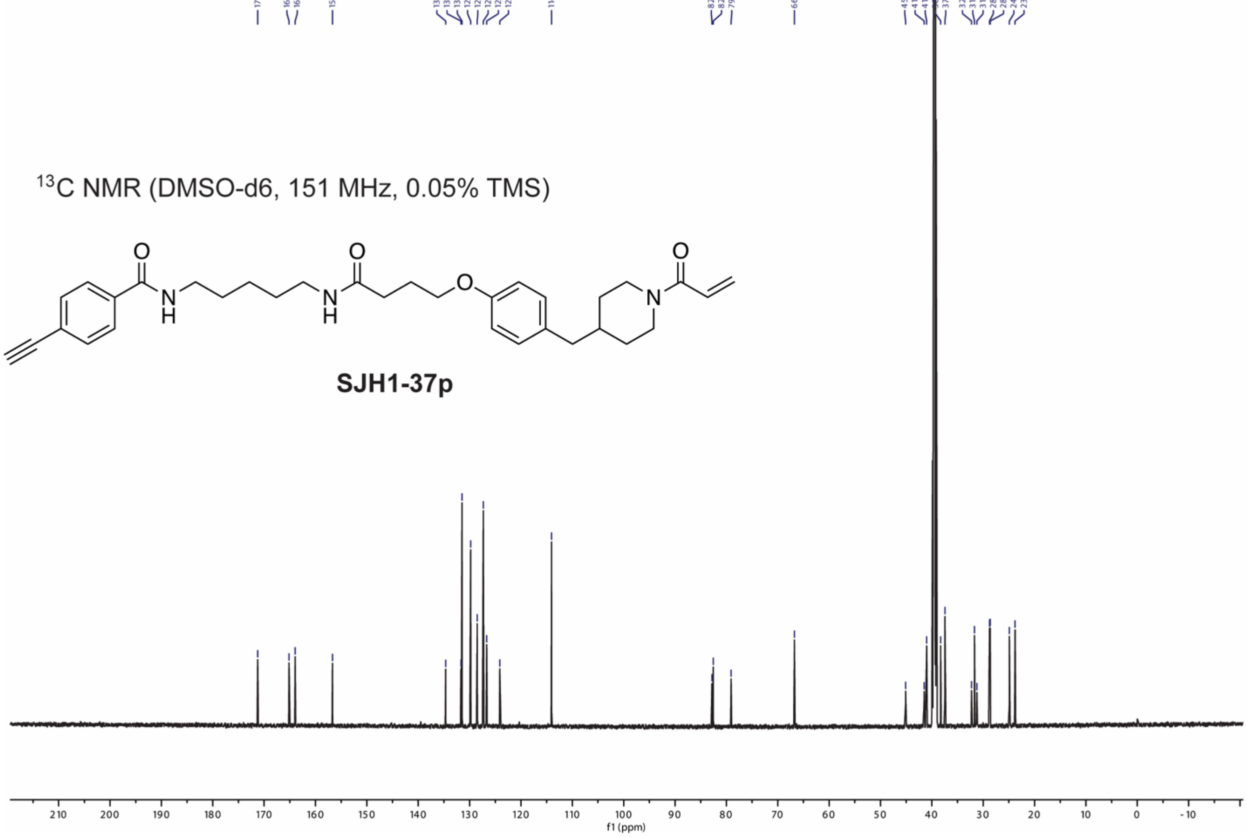
171.25 165.15 163.96 156.70 134.66 131.69 131.48 129.80 128.51 127.33 126.66 124.12 114.03 82.81 82.56 79.08 66.75 45.11 41.49 41.00 37.45 32.27 31.69 31.23 28.77 28.64 24.91 23.78
$^{13}$C NMR (DMSO-d6, 151 MHz, 0.05% TMS)
O
N
H
O
N
H
O
O
N
SJH1-37p
210 200 190 180 170 160 150 140 130 120 110 100 90 80 70 60 50 40 30 20 10 0 -10
f1 (ppm)

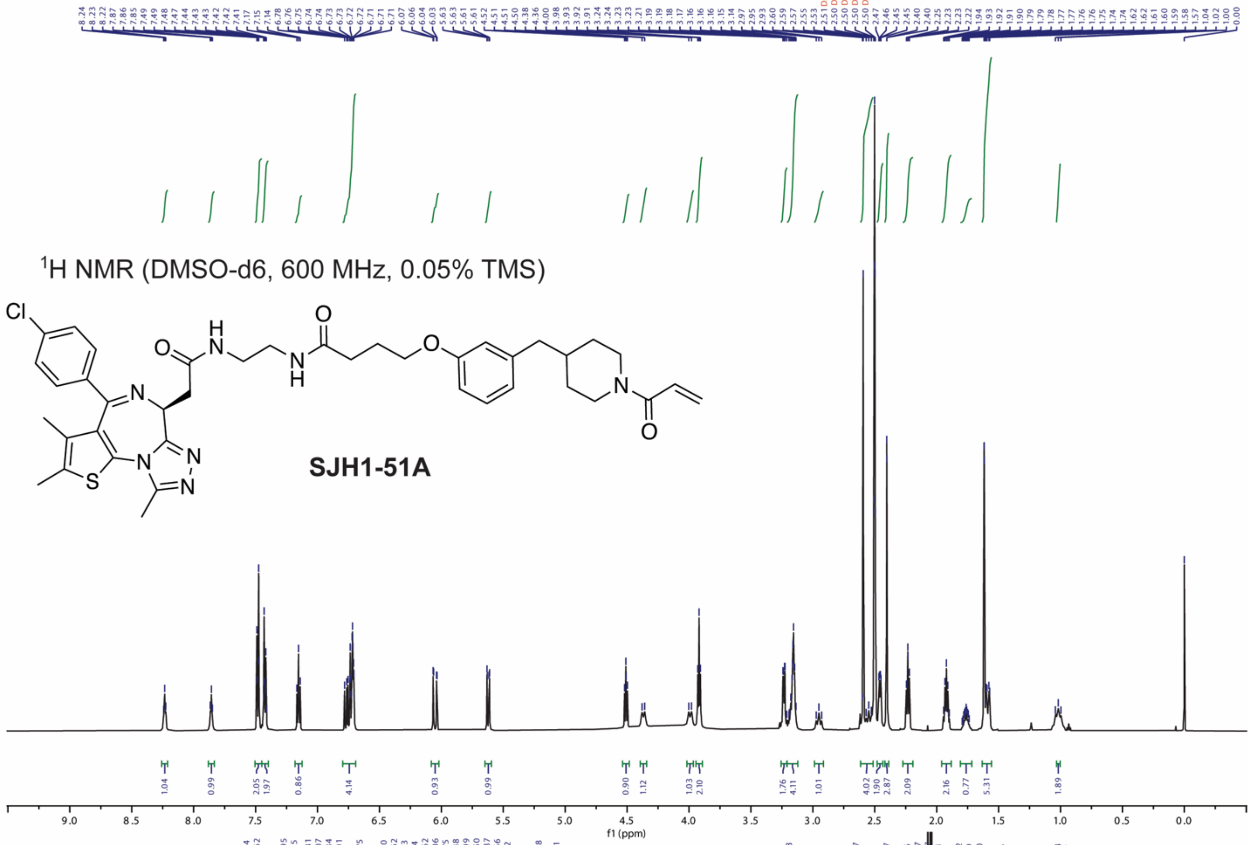

1H NMR (DMSO-d6, 600 MHz, 0.05% TMS)
Cl
O
H
N
N
H
O
O
N
O
N
N
N
S
N
SJH1-51A
f1 (ppm)


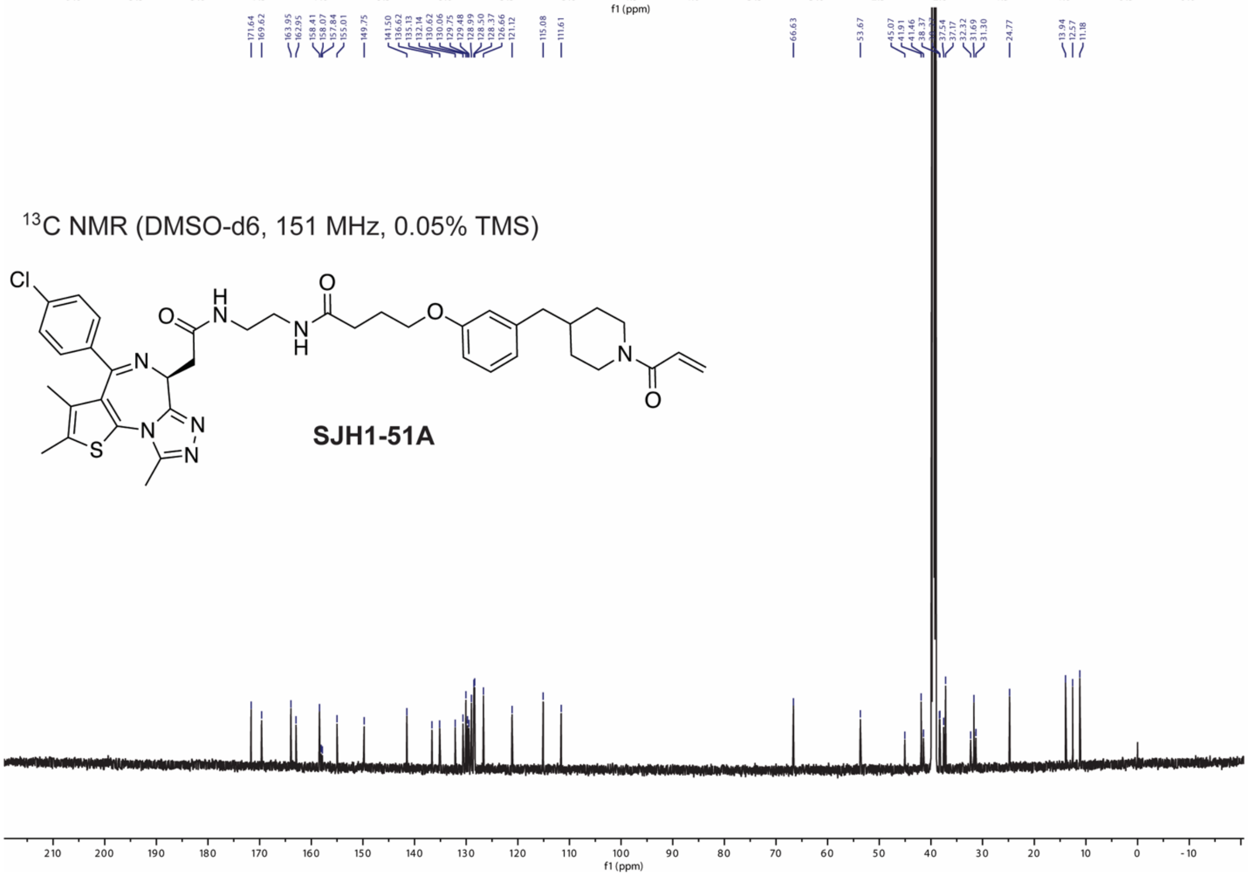

13C NMR (DMSO-d6, 151 MHz, 0.05% TMS)
Cl
O
H
N
N
H
O
O
N
O
N
N
N
S
N
SJH1-51A
f1 (ppm)

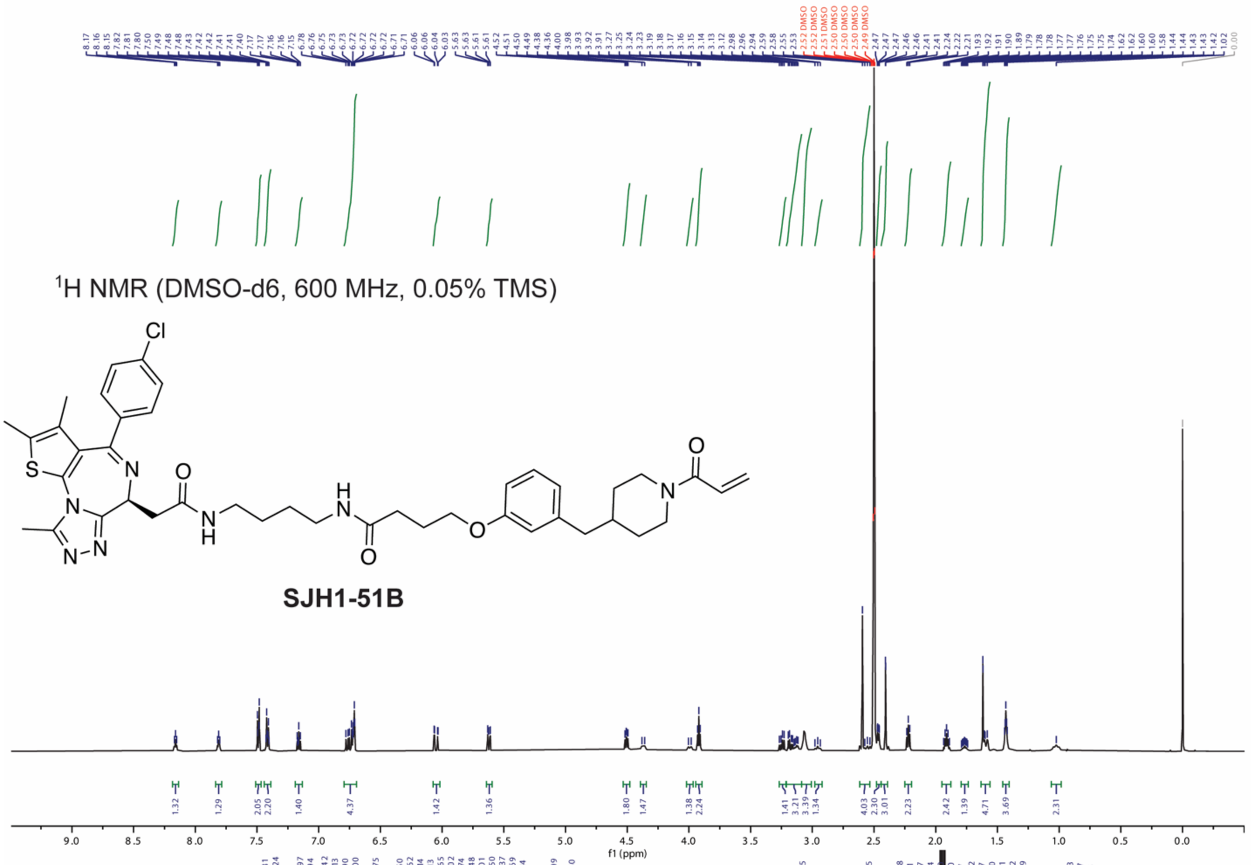

$^{1}$H NMR (DMSO-d6, 600 MHz, 0.05% TMS)
Cl
S
N
O
H
N
O
N
SJH1-51B
f1 (ppm)


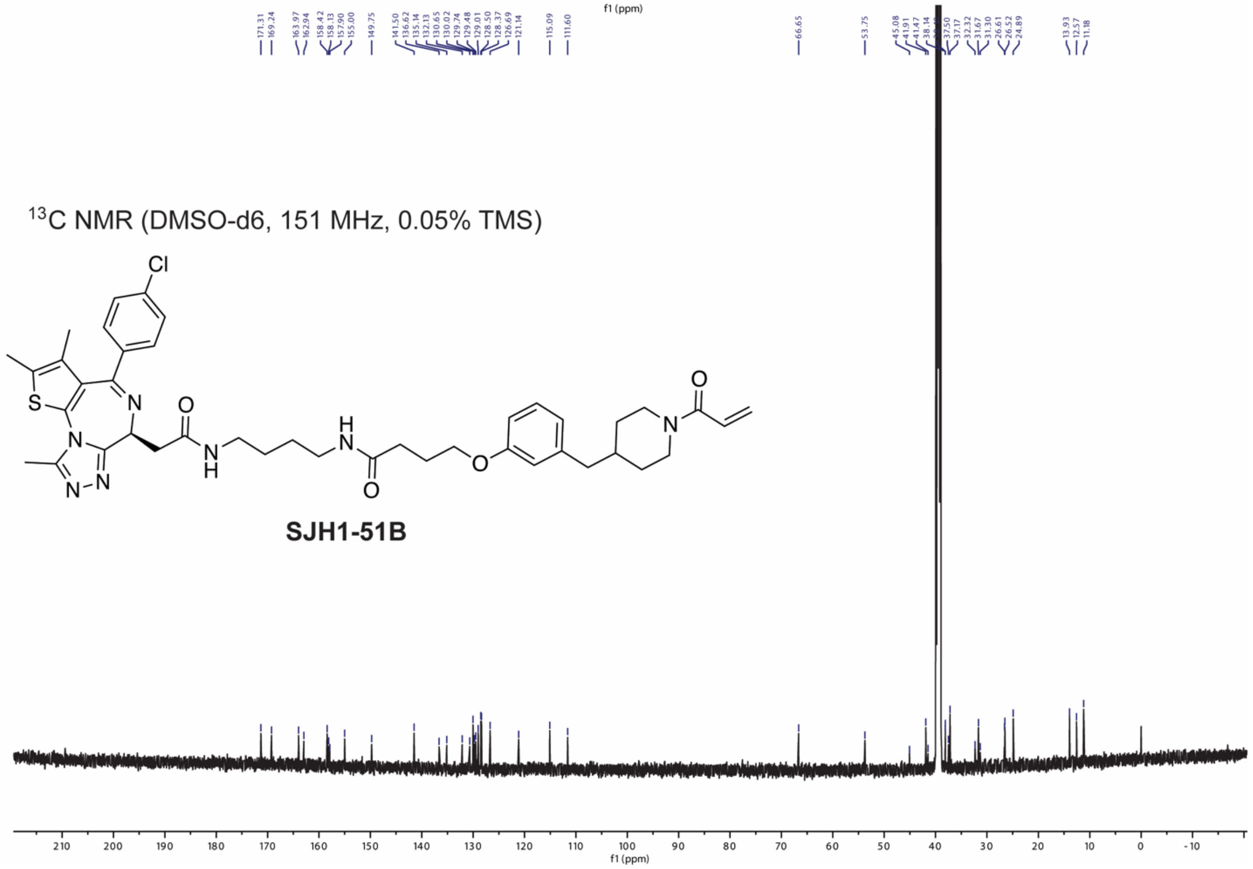

$^{13}$C NMR (DMSO-d6, 151 MHz, 0.05% TMS)
Cl
S
N
O
H
N
O
N
SJH1-51B
f1 (ppm)

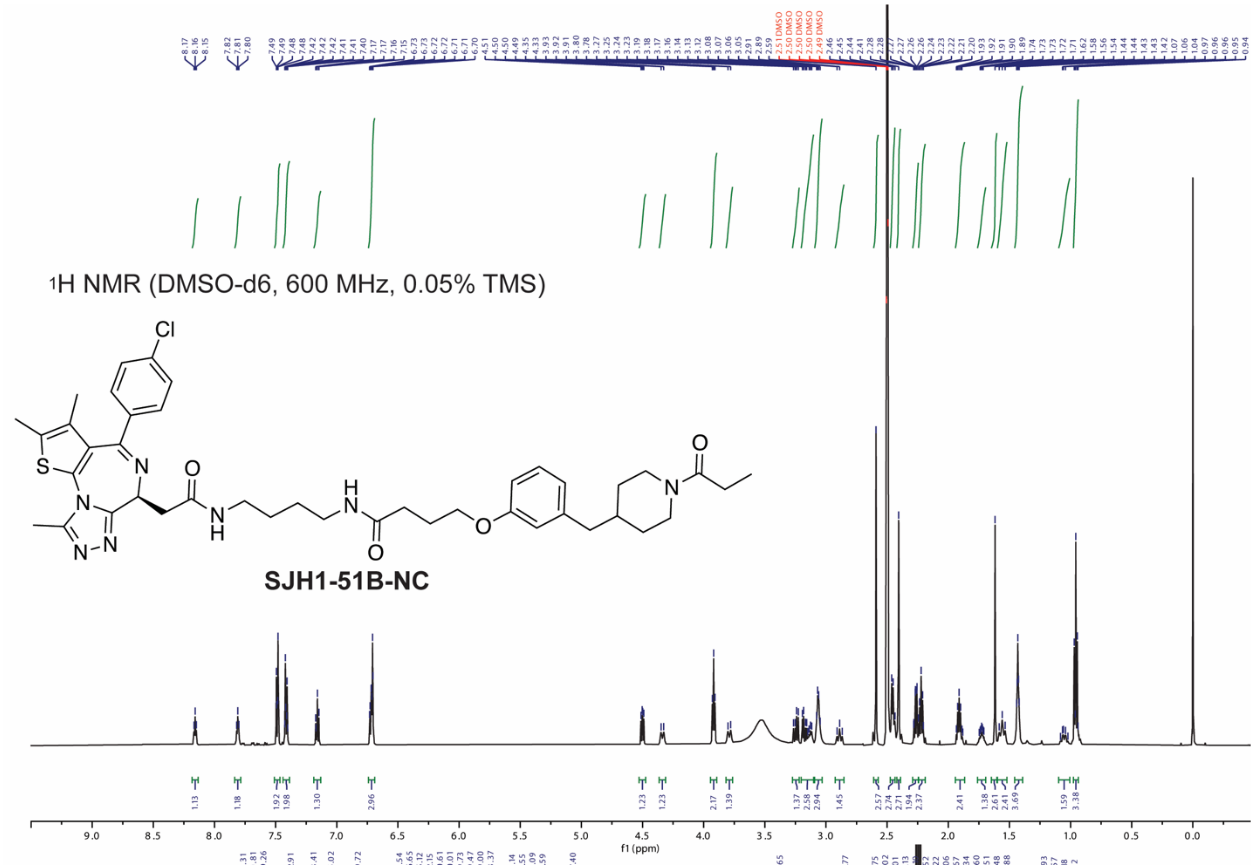

1H NMR (DMSO-d6, 600 MHz, 0.05% TMS)
SJH1-51B-NC
f1 (ppm)


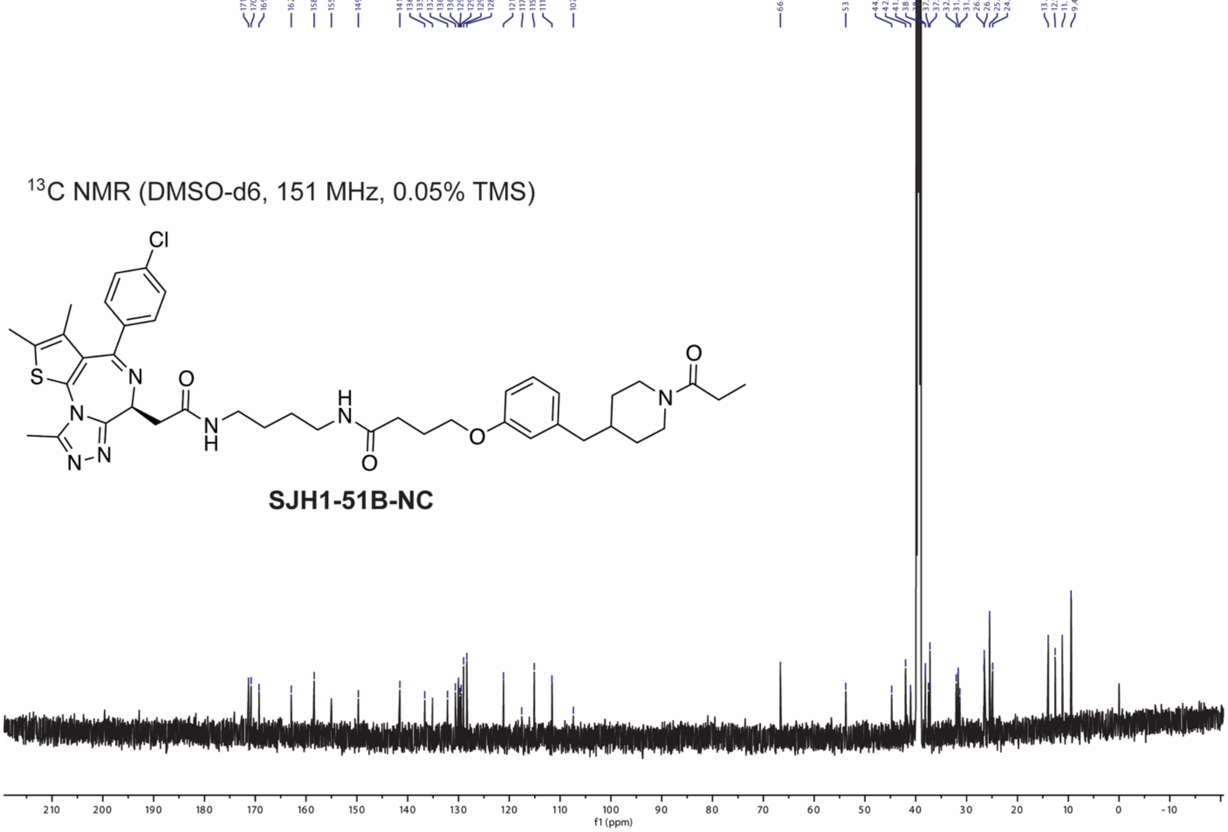

13C NMR (DMSO-d6, 151 MHz, 0.05% TMS)
SJH1-51B-NC
f1 (ppm)

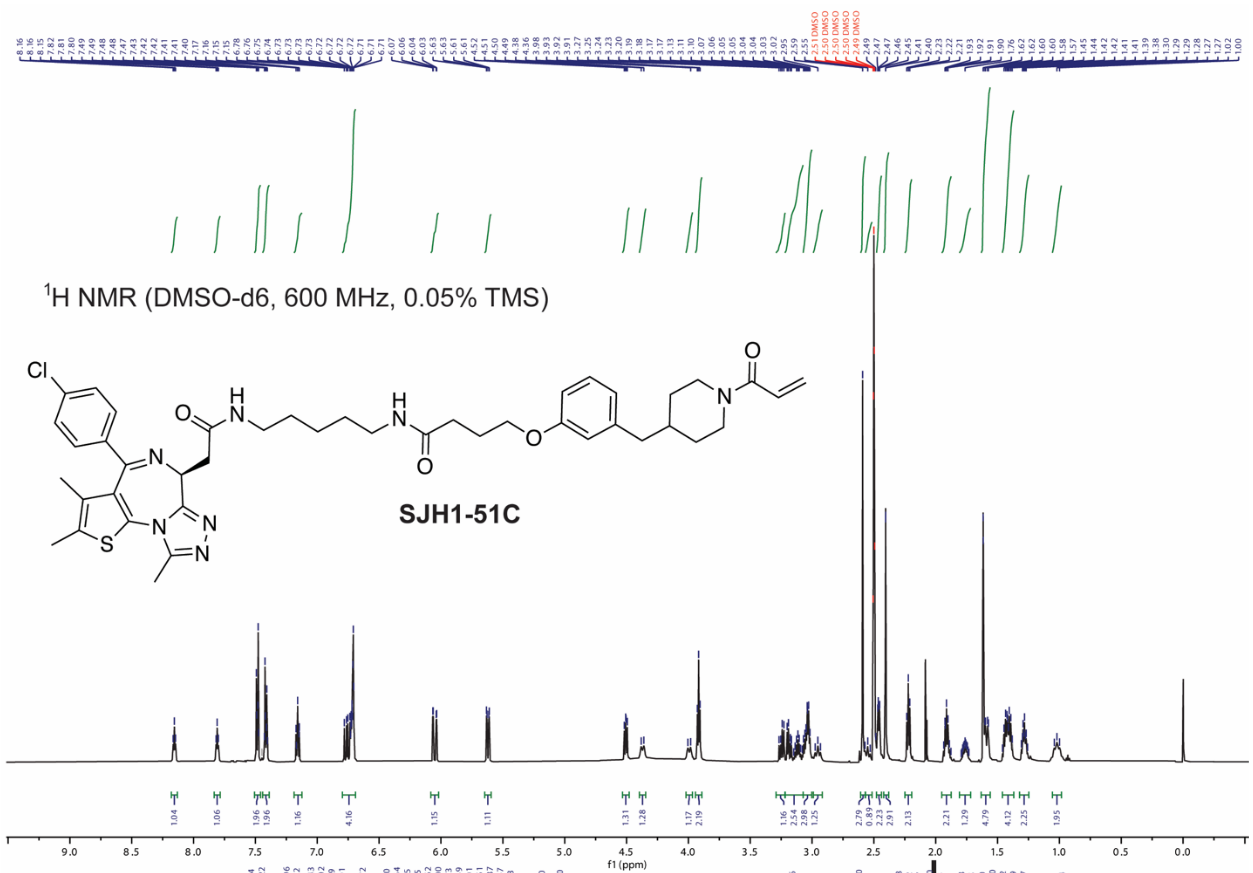

1H NMR (DMSO-d6, 600 MHz, 0.05% TMS)
Cl
O
H
N
N
O
O
N
N
S
N
N
N
SJH1-51C
f1 (ppm)


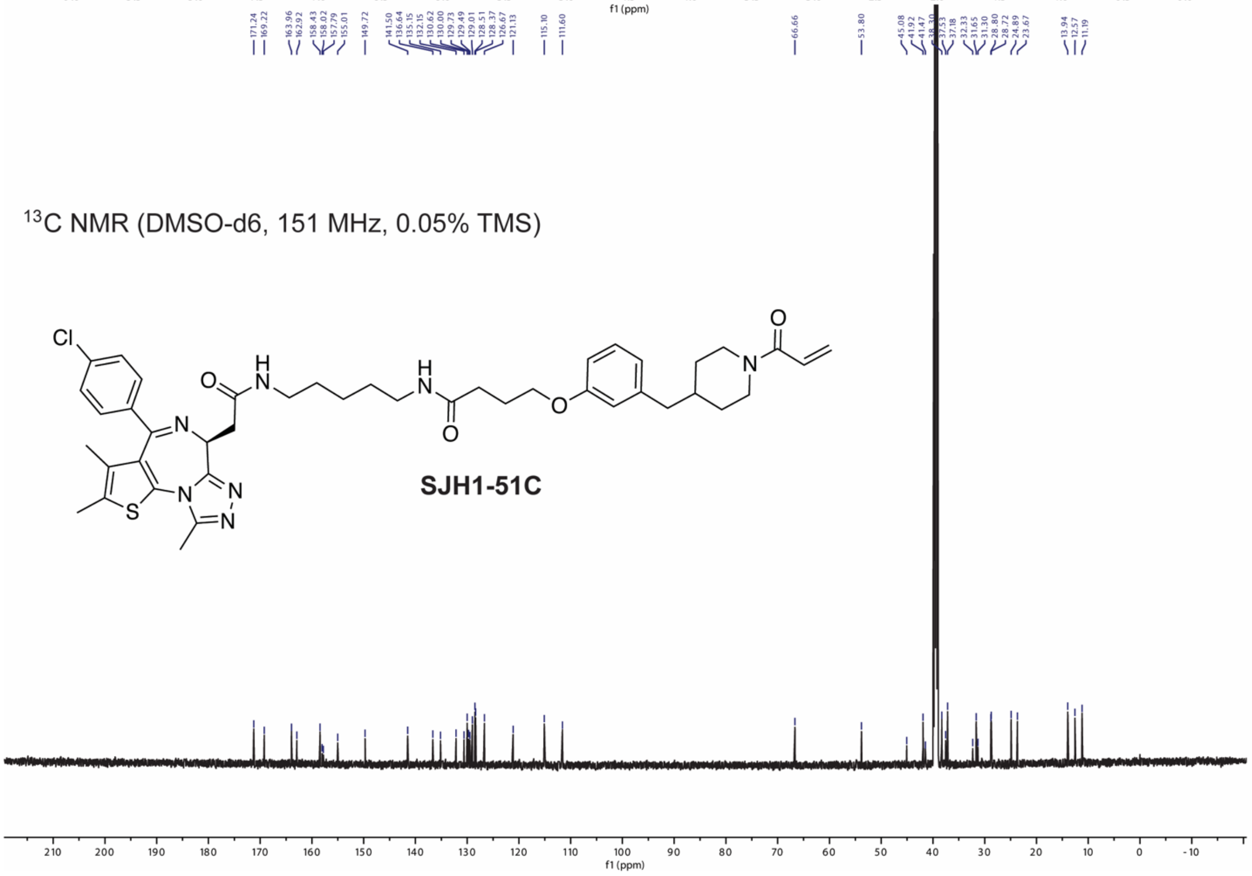

13C NMR (DMSO-d6, 151 MHz, 0.05% TMS)
Cl
O
H
N
N
O
O
N
N
S
N
N
N
SJH1-51C
f1 (ppm)

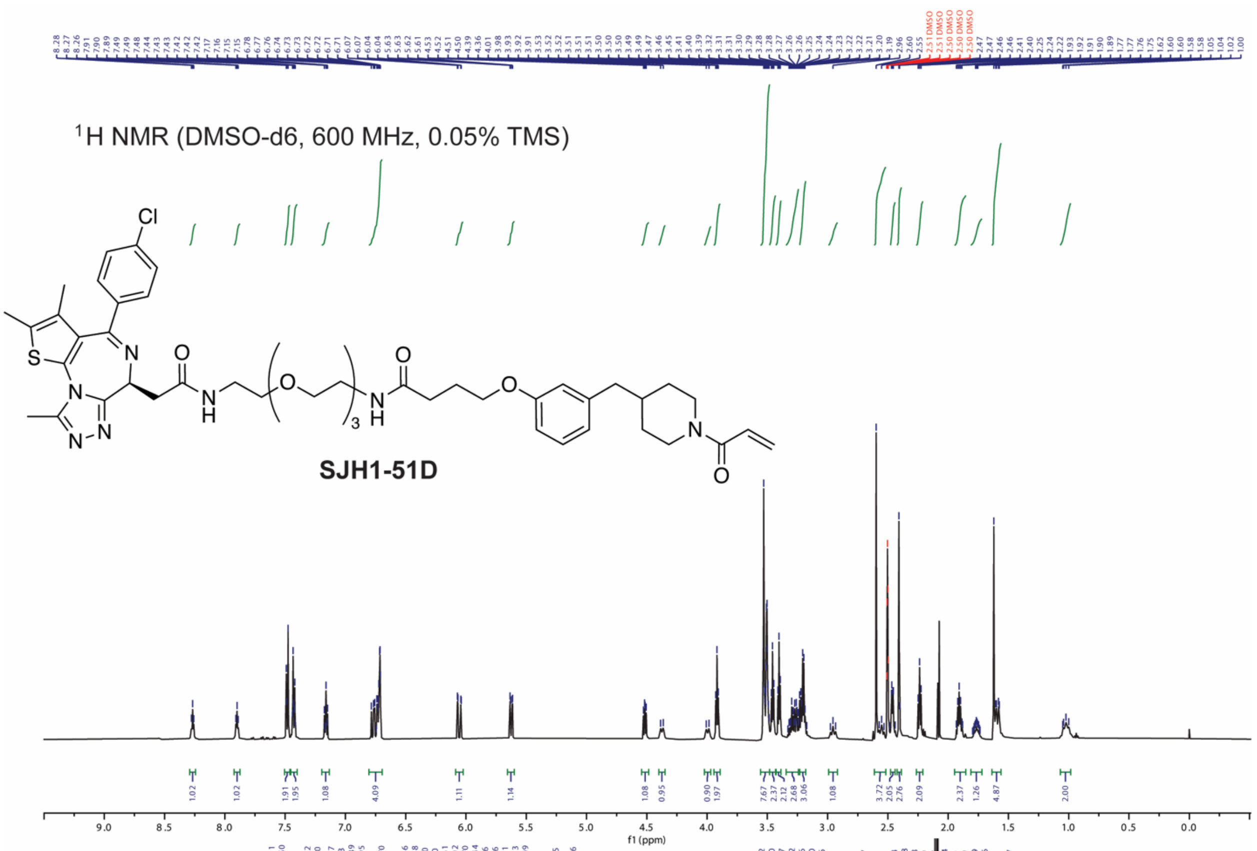

$^{1}$H NMR (DMSO-d6, 600 MHz, 0.05% TMS)
Cl
S
N
O
NH
O
3
H
O
N
O
N
O
N
N
SJH1-51D
f1 (ppm)


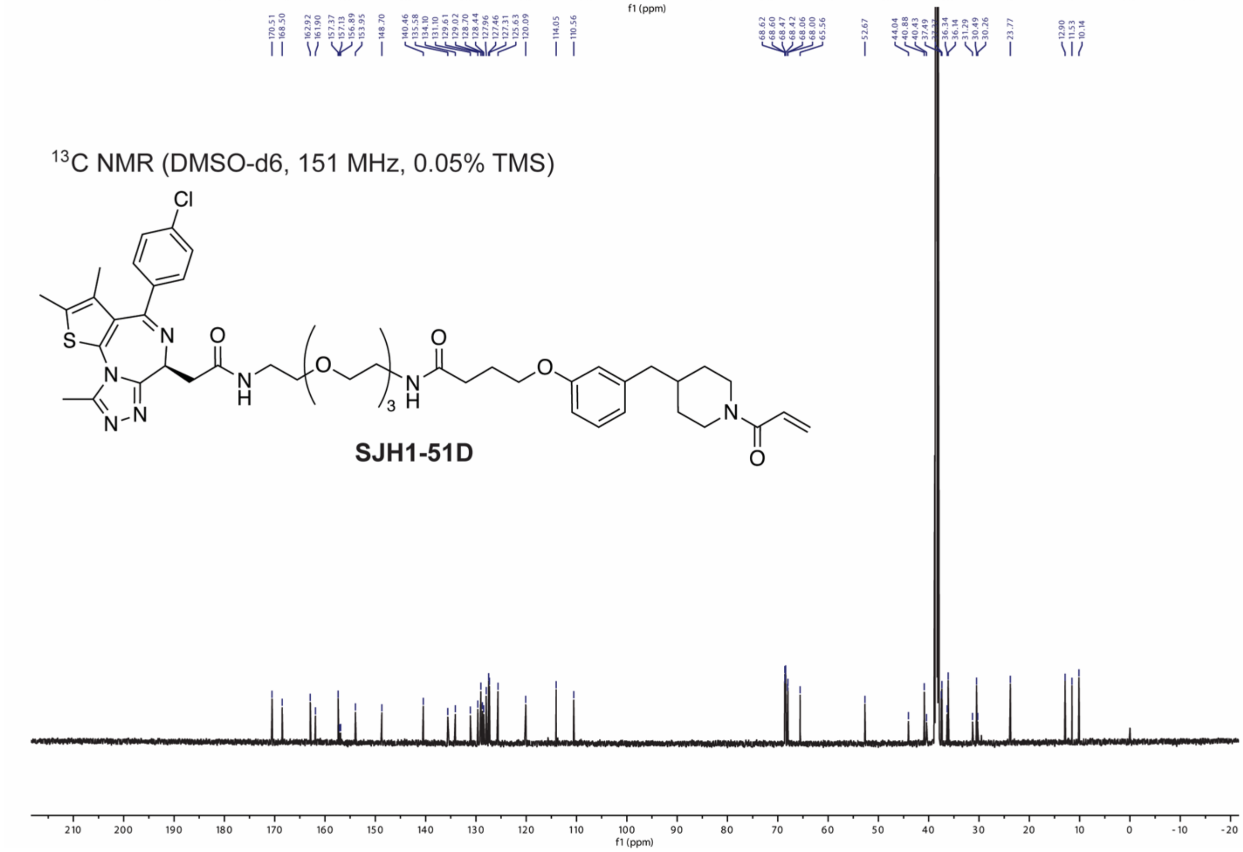

$^{13}$C NMR (DMSO-d6, 151 MHz, 0.05% TMS)
Cl
S
N
O
NH
O
3
H
O
N
O
N
O
N
N
SJH1-51D
f1 (ppm)

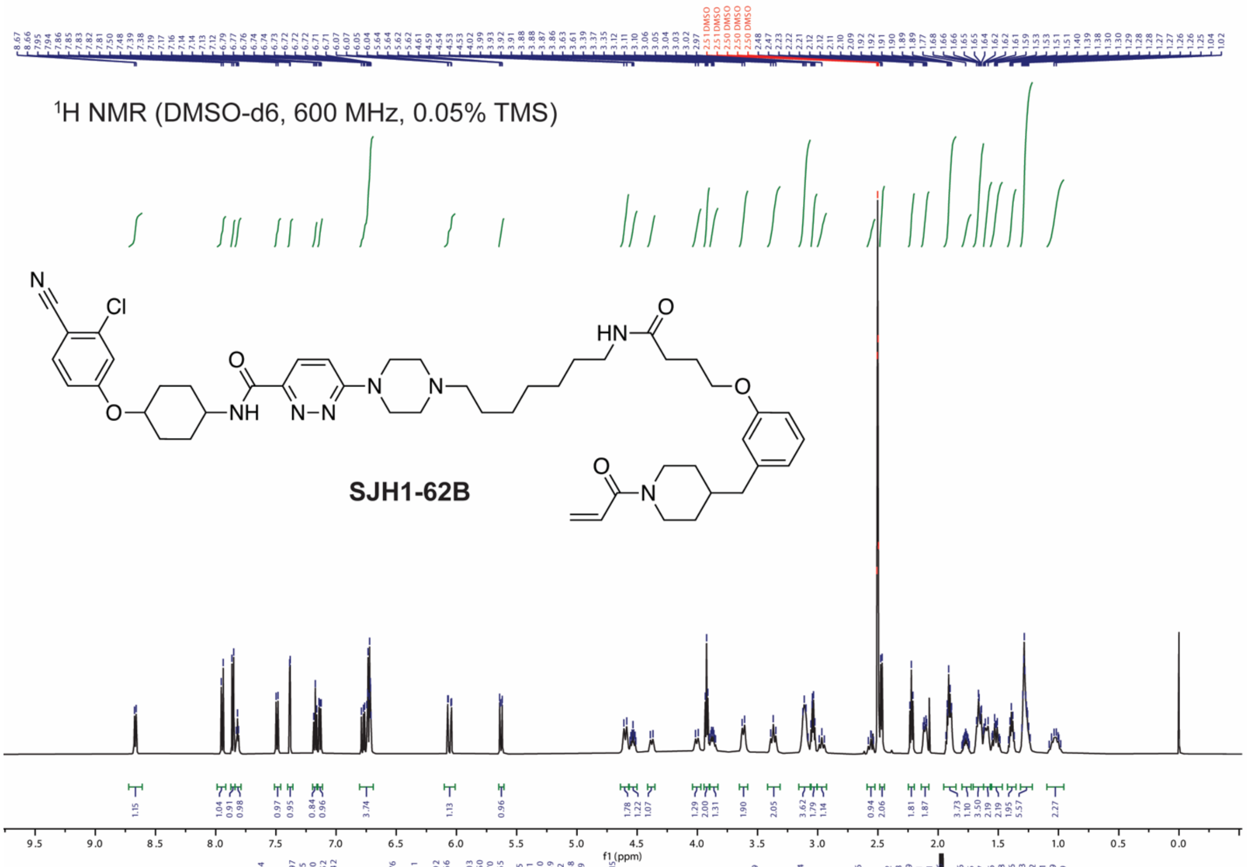

1H NMR (DMSO-d6, 600 MHz, 0.05% TMS)
SJH1-62B
f1 (ppm)


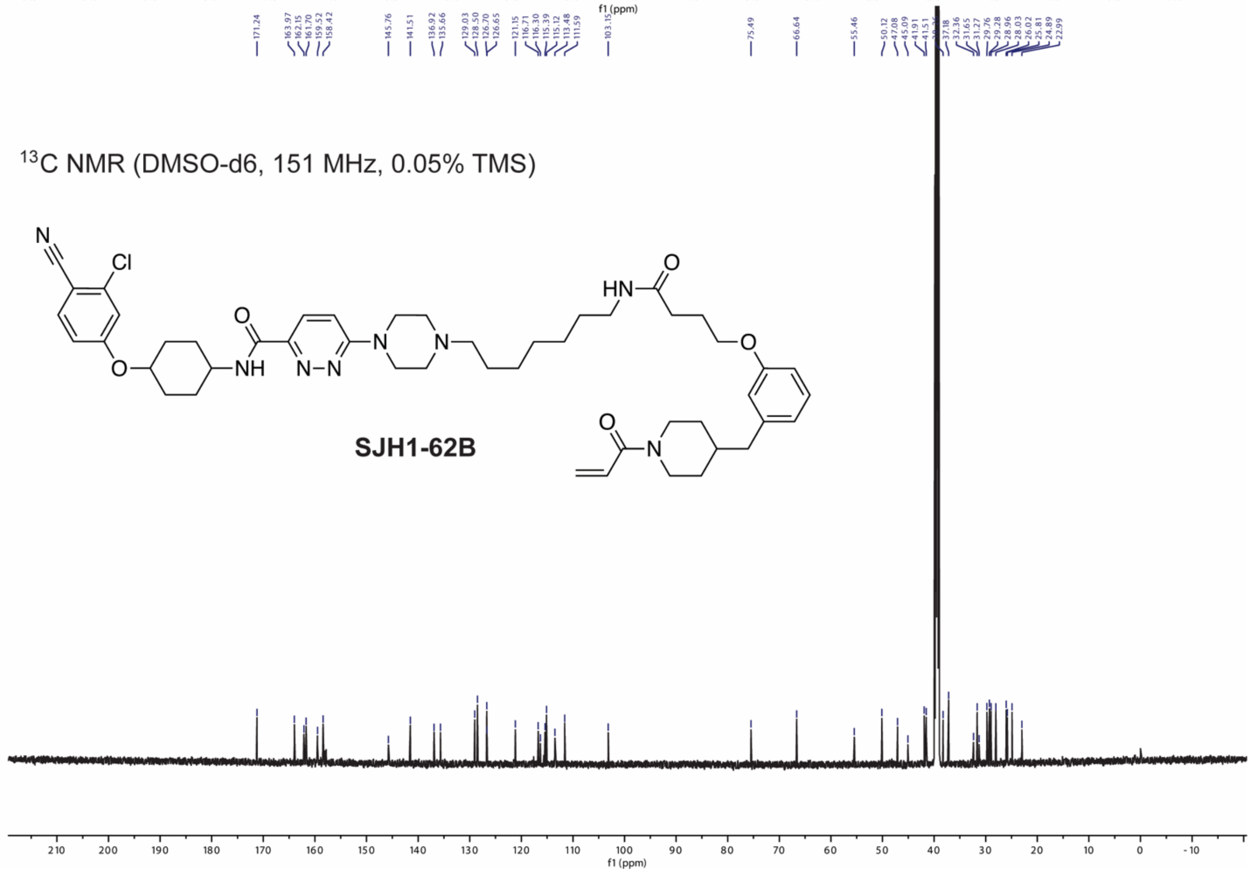

13C NMR (DMSO-d6, 151 MHz, 0.05% TMS)
SJH1-62B
f1 (ppm)

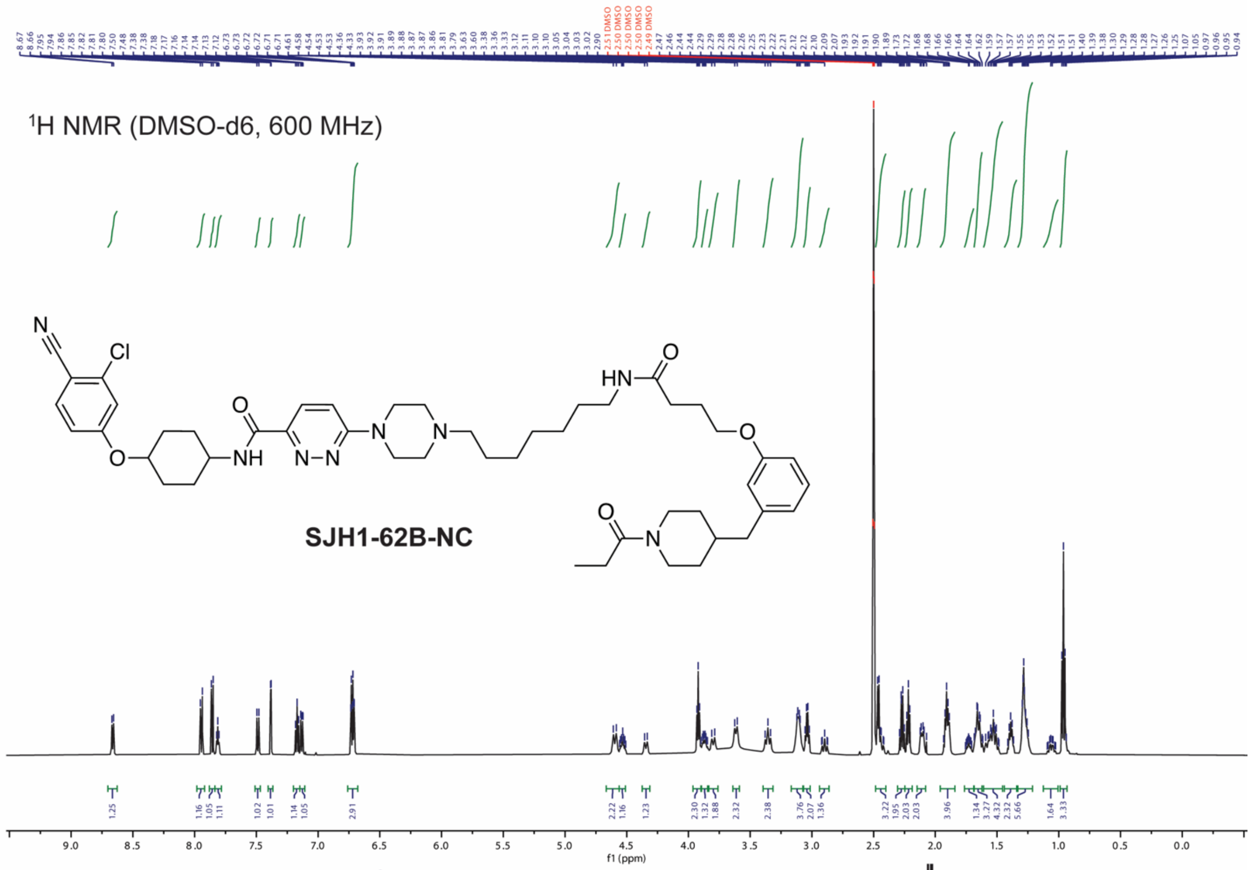

1H NMR (DMSO-d6, 600 MHz)
SJH1-62B-NC
f1 (ppm)


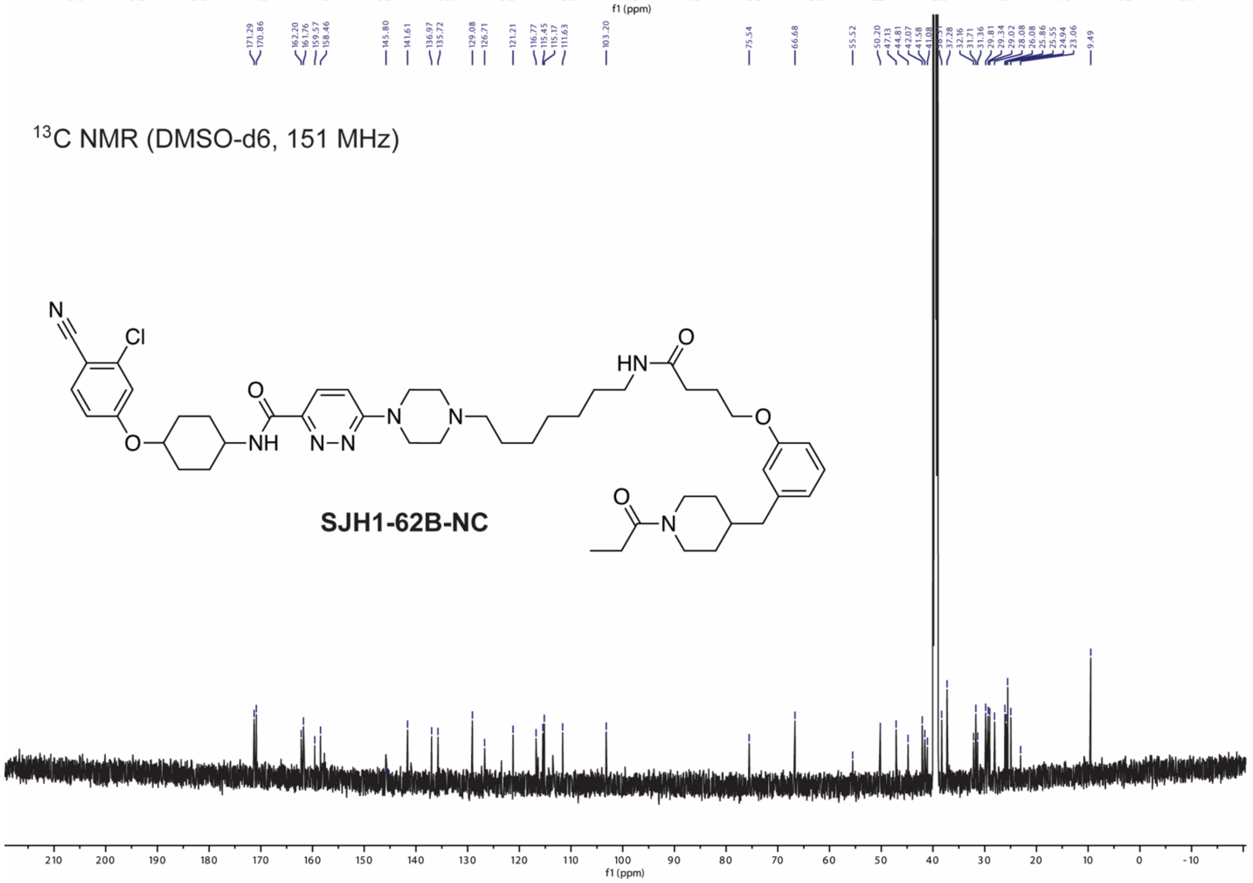

13C NMR (DMSO-d6, 151 MHz)
SJH1-62B-NC
f1 (ppm)

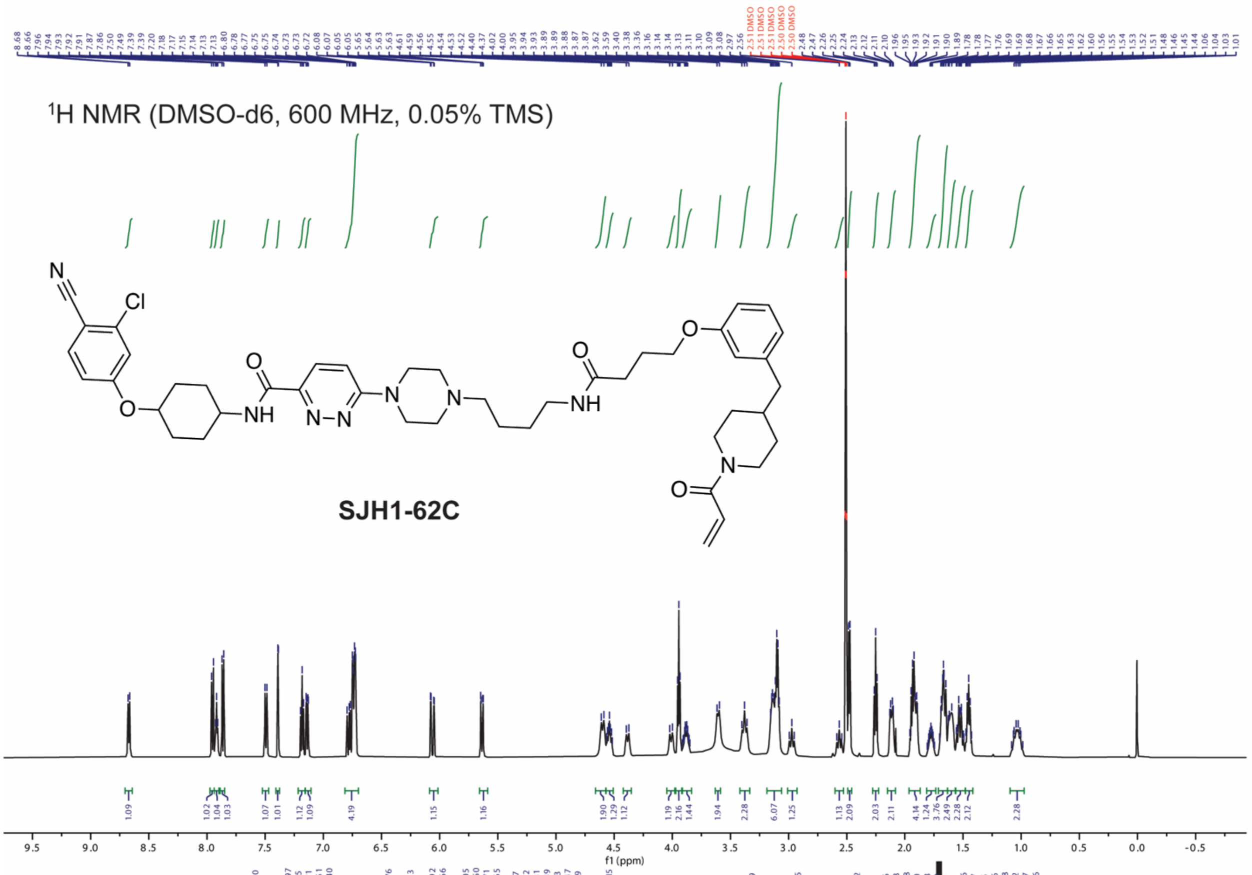

$^1$H NMR (DMSO-d6, 600 MHz, 0.05% TMS)
SJH1-62C
f1 (ppm)


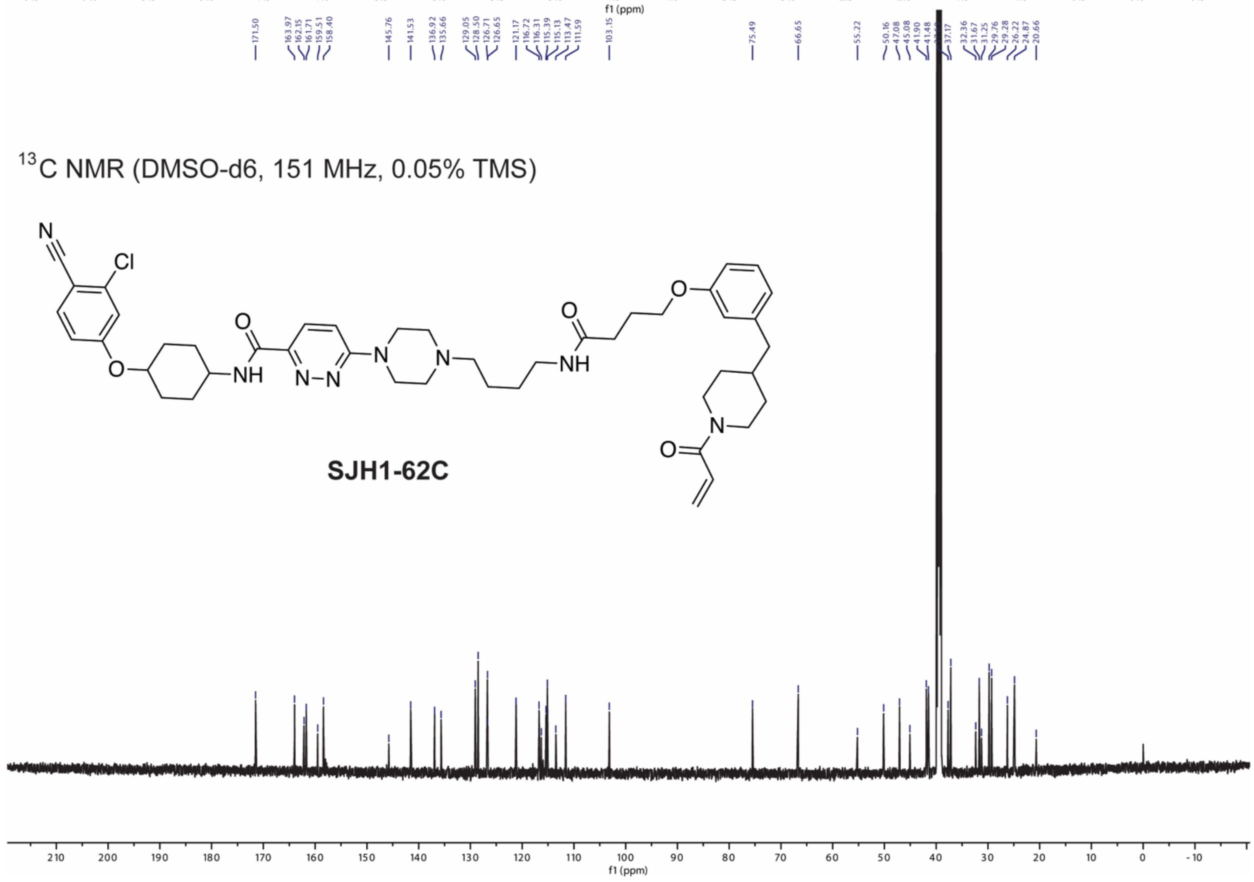

$^{13}$C NMR (DMSO-d6, 151 MHz, 0.05% TMS)
SJH1-62C
f1 (ppm)

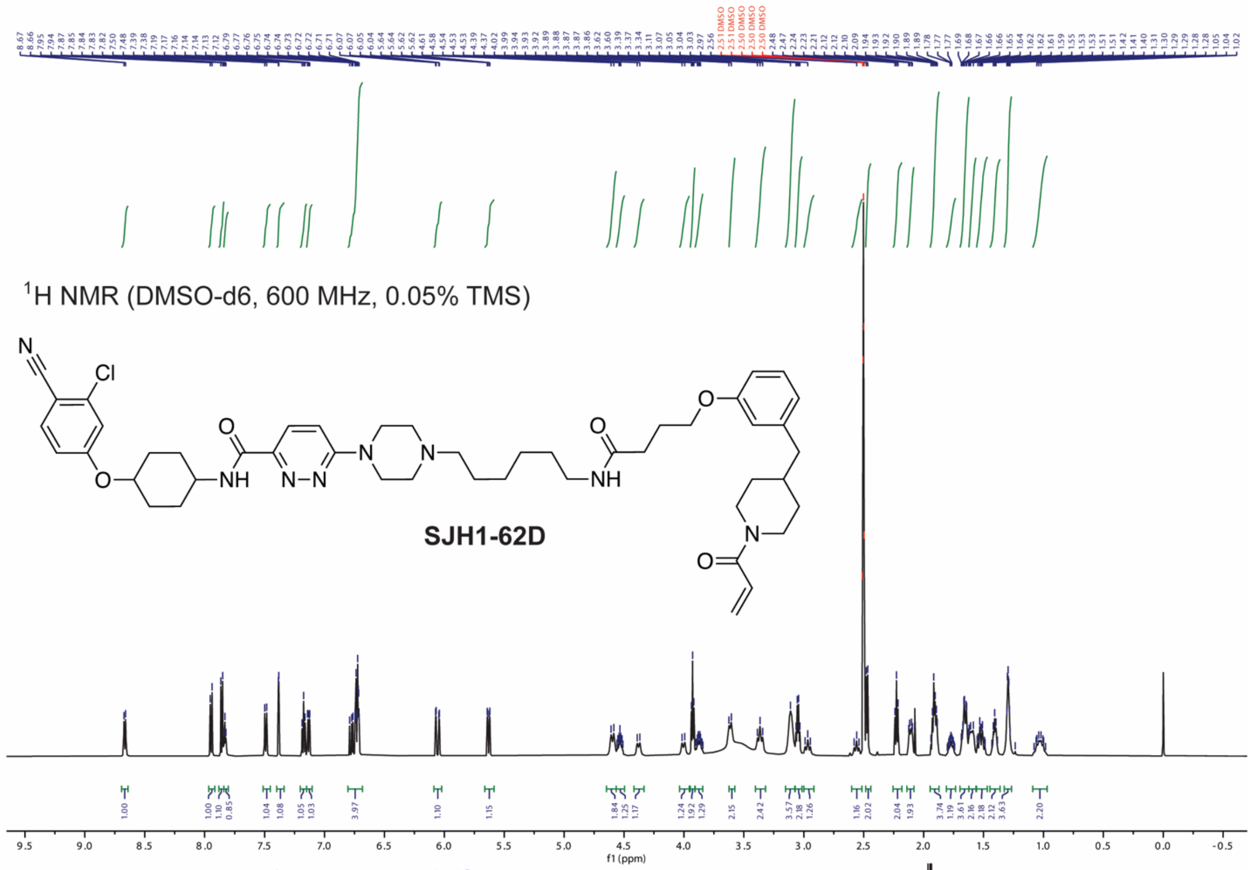

$^{1}$H NMR (DMSO-d6, 600 MHz, 0.05% TMS)
SJH1-62D
f1 (ppm)


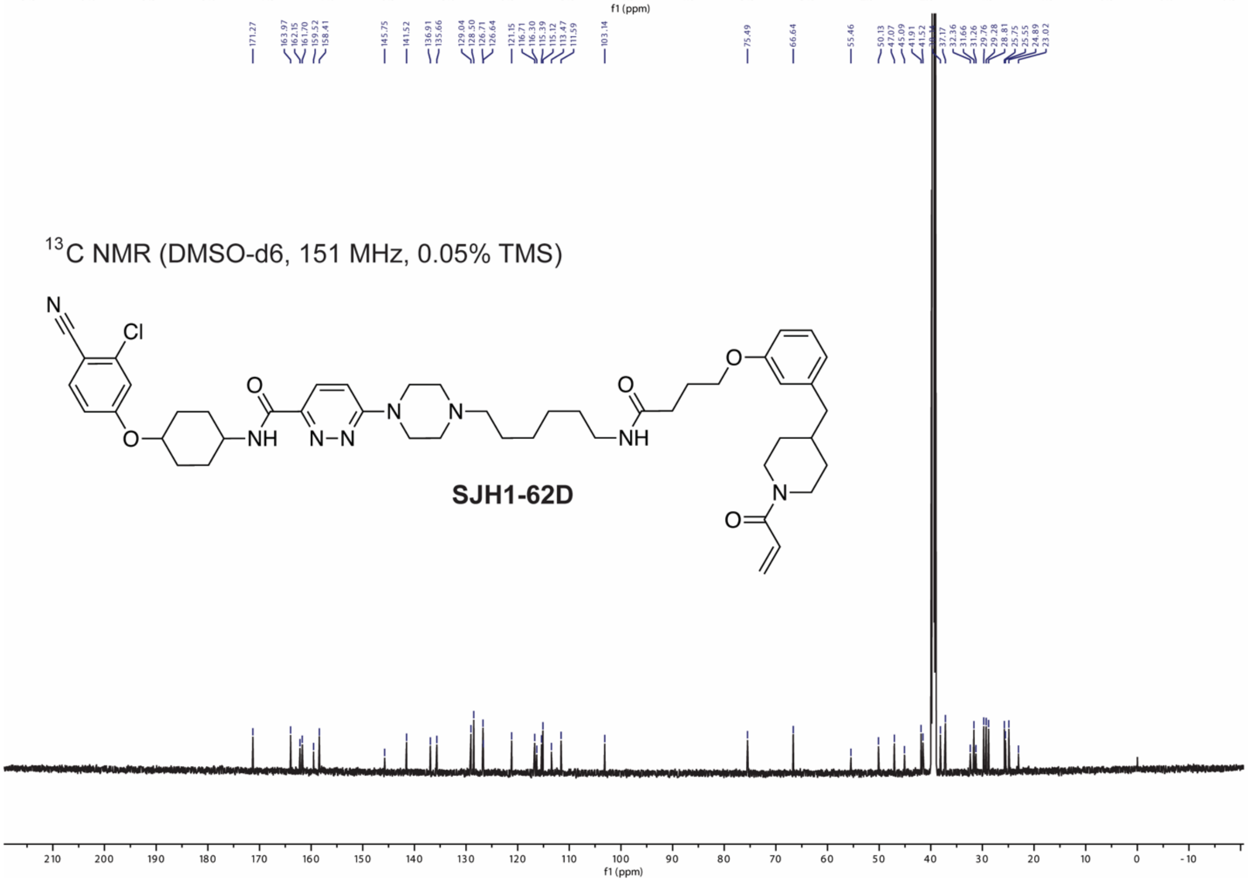

$^{13}$C NMR (DMSO-d6, 151 MHz, 0.05% TMS)
SJH1-62D
f1 (ppm)

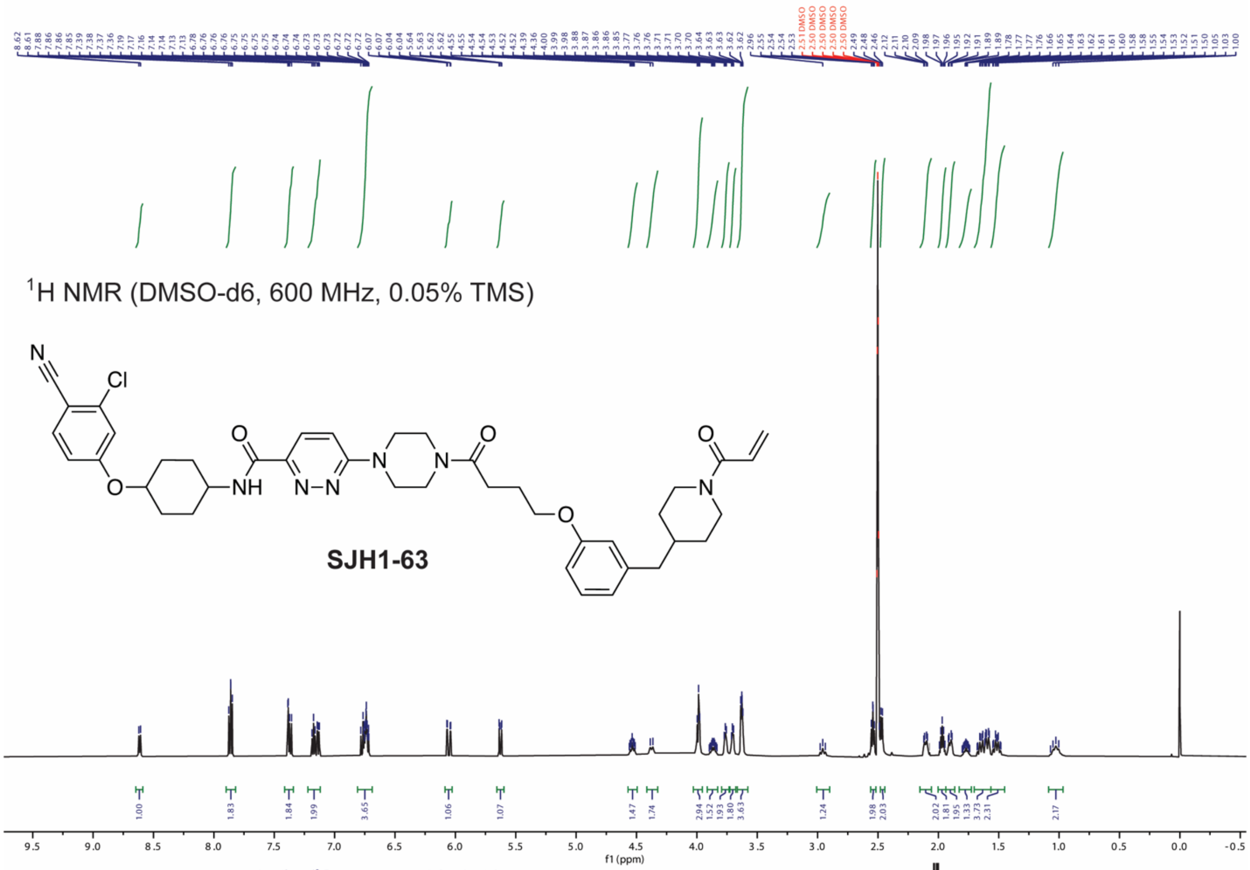
$^{1}$H NMR (DMSO-d6, 600 MHz, 0.05% TMS)
SJH1-63
f1 (ppm)

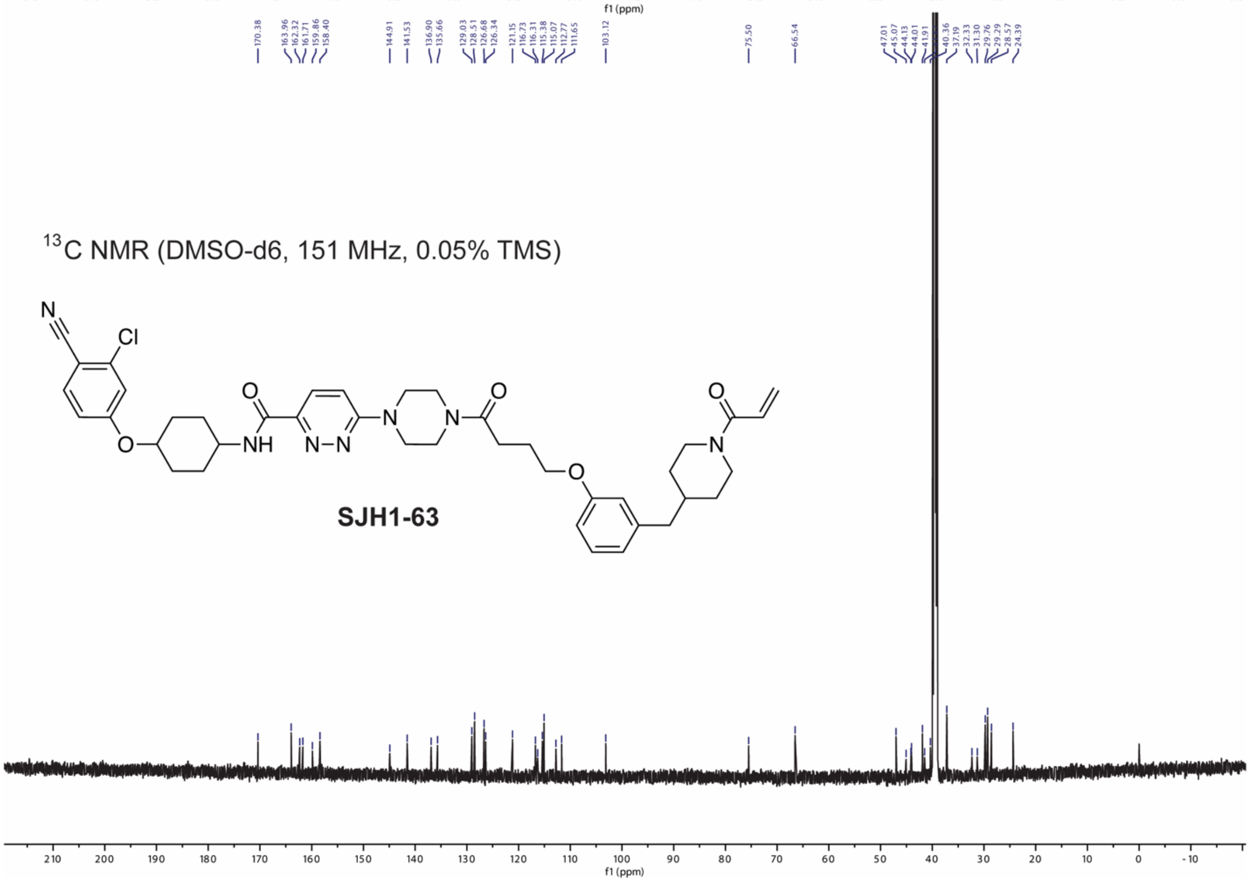
170.38
163.96
162.32
161.71
159.86
158.40
144.91
141.53
136.90
135.66
129.03
128.51
126.68
126.34
121.15
116.73
116.31
115.38
115.07
112.77
111.65
103.12
75.50
66.54
47.01
45.07
44.13
44.01
41.91
40.36
37.19
32.33
31.30
29.76
29.29
28.57
24.39
$^{13}$C NMR (DMSO-d6, 151 MHz, 0.05% TMS)
SJH1-63
f1 (ppm)
